# Translational products encoded by novel ORFs may form protein-like structures and have biological functions

**DOI:** 10.1101/567800

**Authors:** Chaitanya Erady, David Chong, Narendra Meena, Shraddha Puntambekar, Ruchi Chauhan, Yagnesh Umrania, Adam Andreani, Jean Nel, Matthew T. Wayland, Cristina Pina, Kathryn S. Lilley, Sudhakaran Prabakaran

## Abstract

Translation products encoded by non canonical or novel open reading frame (ORF) genomic regions are generally considered too small to play any significant biological role, and dismissed as inconsequential. In this study, we show that mutations mapping to novel ORFs have significantly higher pathogenicity scores than mutations in protein-coding regions. Importantly, novel ORFs can translate into protein-like structures with putative independent biological functions that can be of relevance in disease states, including cancer. We thus provide strong evidence to support the systematic study of novel ORFs to gain new insights into normal biological and disease processes.

**One Sentence Summary:** Non coding regions may encode protein-like products that are important to understand diseases.

## Main Text

All known human proteins are coded by just 1.5% of the genome; however, genome-wide association studies (GWAS) have indicated that ∼93% of disease-and trait-associated variants map to noncoding regions [1]. Recent work has not only shown that transcription and translation can occur pervasively over the entire genome, but these non canonical transcriptional and translated products encoded by as yet undefined or novel ORFs could also be biologically regulated [2–4] and may even be used as the main source of targetable tumor-specific antigens [5]. Analysis of all the GWAS associated variants and mutations in the Catalogue of Somatic Mutations in Cancer (COSMIC) and Human Gene Mutation Database (HGMD) databases has revealed that a significant proportion of the variants and mutations map to apparent non coding regions of the human genome (**Fig. 1A.**). Many of these mutations and variants map to regulatory regions of known genes, such as promoters, enhancers, and transcription factor binding sites, and are implicated in disease. For the majority of these mutations, however, their functional consequences cannot be interpreted and a potential relationship with novel ORFs has not been assessed.

**Fig. 1.**
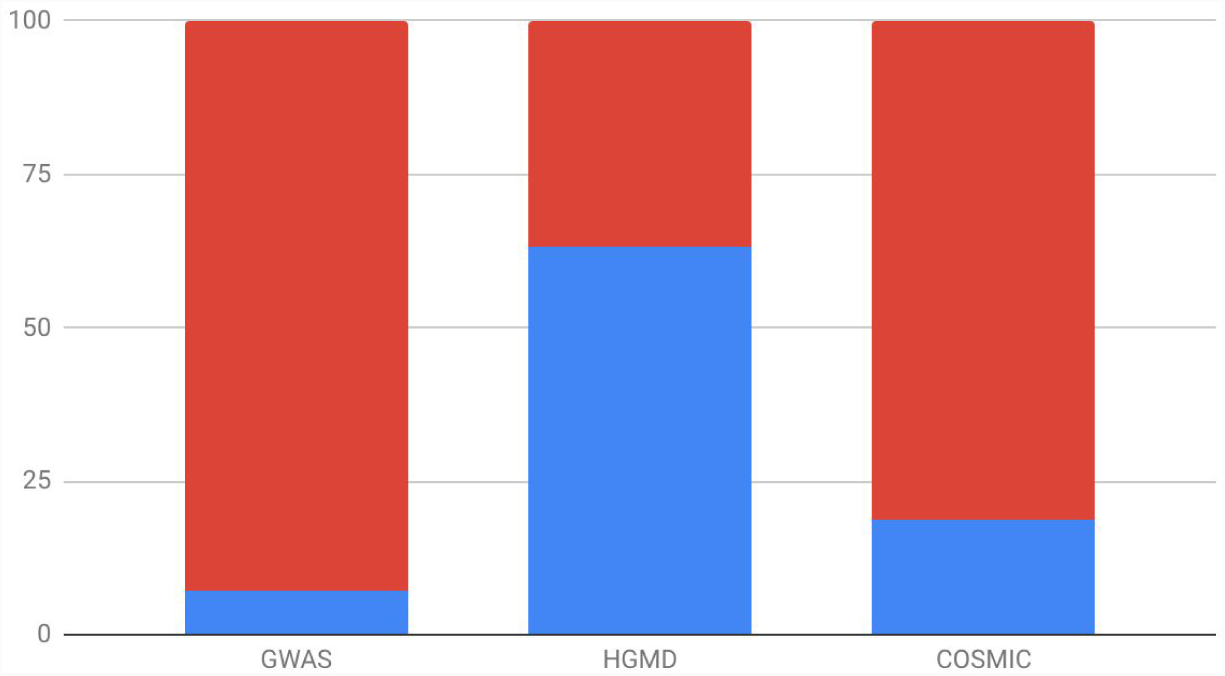
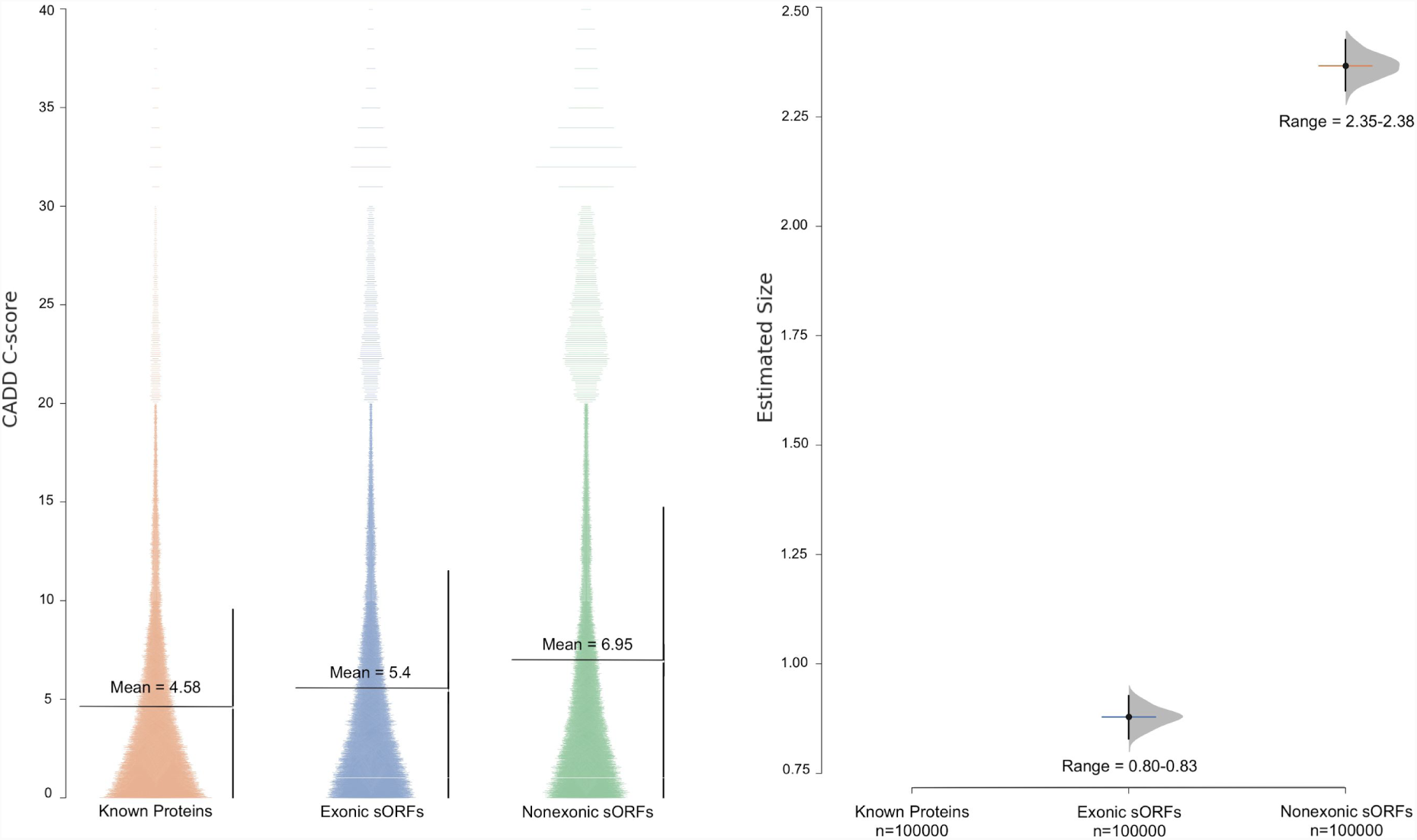
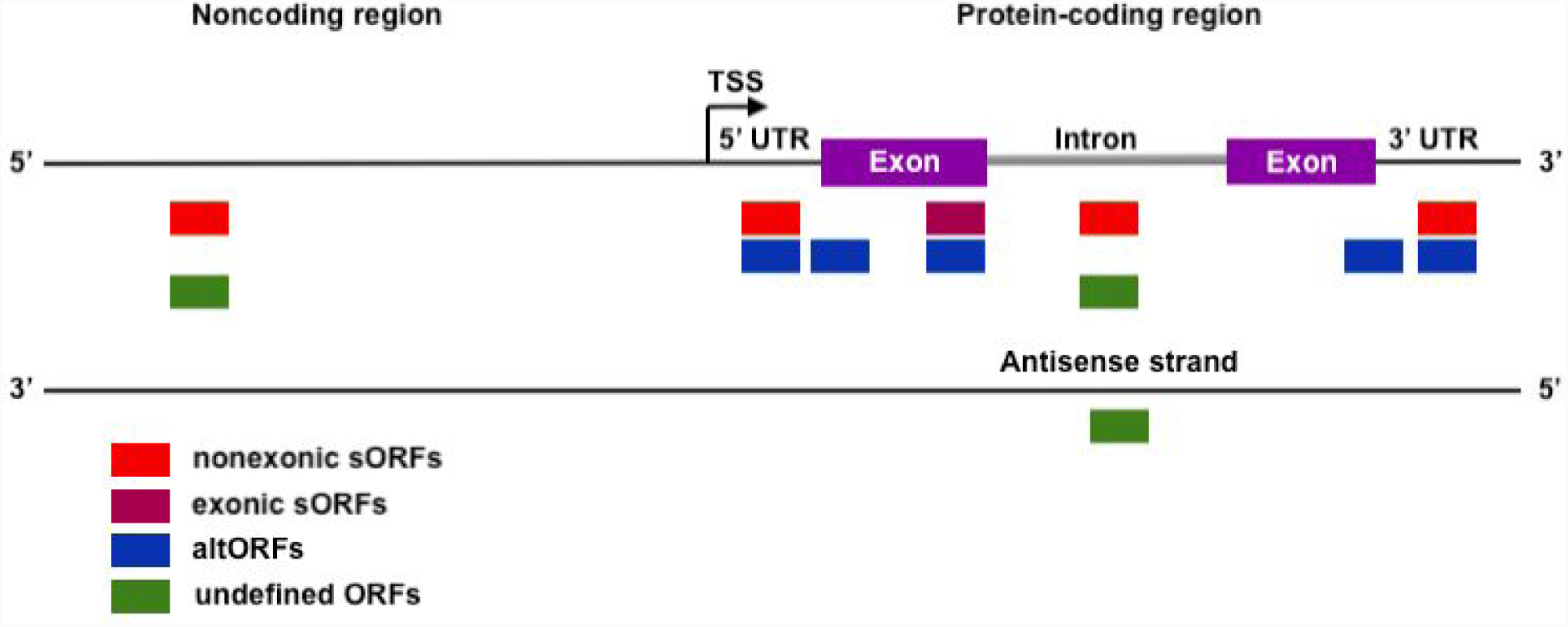
Reasons to investigate novel ORFs. (**A**) Proportion of coding (blue) vs non coding (red) disease-associated variants in GWAS, HGMD, and COSMIC datasets. (**B**) Left panel shows the CADD score distribution and their mean values mapped to known proteins, sORFs in the exonic regions, and sORFs in the non exonic, regions. Right panel is the estimation size plot of the CADD scores showing the mean difference with 95% confidence interval of all variants mapped to exonic sORFS (range 0.80-0.83) and non exonic sORFs (range 2.35-2.38) with respect to known proteins. (**C**) Schematic representation of the three types of novel ORFs that were investigated in this study. They include non exonic and exonic sORFs (red and maroon boxes), altORFs (blue box), and as yet undefined ORFs (green box). sORFs and undefined ORFs can be found both within coding (including 5’UTR and 3’UTR) and non coding regions; whereas, altORFs are defined as translation products found only in the alternate frames of known protein coding regions. Sometimes sORFs can be classified as altORFs and vice versa.

In spite of a large proportion of disease- and trait-associated variants mapping to novel ORF regions there has not been a systematic investigation of the products encoded by these novel ORFs. This lack of comprehensive interrogation of novel ORFs has arisen for a variety of reasons. Firstly, non canonical translated products encoded by novel ORFs have been largely dismissed as false positives [6] and as biological noise. Secondly, some products are genuinely too small to play any significant role in biological processes [7]. Finally, improper experimental design that does not allow for the collection of total RNA sequencing and mass spectrometry data from the same samples, use of rigid and conservative genome annotations frameworks [8], and the lack of systematic investigation of their potential structures have contributed to our lack of knowledge about products of novel ORF transcription and translation. Yet, the Combined Annotation Dependent Depletion (CADD) scores or the pathogenicity scores [9] of all variants that map to a subset of well-characterised novel ORFs in the human genome called the short Open Reading Frames (sORFs) [10,11], are marginally more than the CADD scores of variants that map to all the known proteins. **Fig. 1B. left panel**, shows the distribution of CADD scores for variants that map to (a) known proteins encoded by known ORFs, (b) sORFs that overlap known ORFs, known as exonic sORFs, and (c) sORFs that are present in non coding regions. **Fig. 1B. right panel** demonstrates that the distribution of mean CADD scores of variants of sORFs in the non coding regions that are significantly higher than the mean CADD scores of variants that map to exonic sORFs and known proteins. This indicates that the deleterious effects of variants that map to non exonic sORFs in the non coding regions are greater than the deleterious effects of variants on known proteins. It is thus of paramount importance to identify and investigate the role of all novel ORFs in biological and disease processes.

In this study to add insight to the potential functionality of novel ORFs, we take a proteogenomics approach combining total RNA sequencing data of naive B and T cells (GEO94671 [12] from the Blueprint consortium with in-house generated proteomics data from a similar experimental design. We perform comprehensive and systematic analyses of this data to identify novel ORFs in the naive B and T cells from inbred mice that have not been exposed to antigens. It was crucial to use naive B and T cells because we wanted each of the cellular transcriptome and proteome in the population to be identical. We also investigated whether the functions of human orthologs of the identified novel ORFs are disrupted in diseases by predicting their structures [13] and mapping disease- and trait-associated mutations from COSMIC and HGMD on to them

## Results

We first identified novel ORFs in naive mouse B and T cells and inferred potential functions of the products encoded by them by (a) using their sequences, (b) predicting their structures, and (c) inferring whether they compromise biological functions in diseases by mapping disease-associated mutations from HGMD and COSMIC databases. Briefly, total RNA was extracted from naive B and T cells isolated from spleen of six male and six female C57BL/6J mice and subjected to total RNA sequencing (GEO accession GSE94671) (Table S1). Similarly, proteins were extracted from naive B and T cells isolated from spleen of a different set of six male and six female BL6 mice and analysed using mass spectrometry (Fig. S1). The novel ORFs regions that were systematically investigated are schematically represented in **Fig. 1C.** and include (a) sORFs, (b) altORFs [14], ORFs in the alternate frame of known protein coding genes, and (c) all other as yet undefined novel ORFs. Transcriptional evidence alone, or translational evidence alone, or both, were used to identify novel ORFs.

### Identification of novel ORFs with transcriptional evidence

HISAT2 [15] was used to align ∼73 million sequenced reads to the reference genome, which resulted in ∼50 million aligned reads (Fig. S2-S4). An average of ∼194,000 transcripts were then assembled for each of the cell types by StringTie [15] (Fig. S5-S7). Merging the transcripts across the 12 samples resulted in a total of ∼164,000 transcripts, which after filtering gave 109,441 transcripts (Fig. S8), of these, 101,767 transcripts were expressed in B cells and 99,552 transcripts were expressed in T cells. These transcripts were then used to construct cell-type specific nucleotide databases for subsequent proteogenomic analysis (**Fig. 2A**). Of the 109,441 transcripts, 91,878 transcripts are common to B and T cells, 9,889 are unique to B cells, 7,674 are unique to T cells (Fig. S9). 30,270 transcripts are known protein coding transcripts in B cells, and 29,235 are known protein coding transcripts in T cells.

**Fig. 2.**
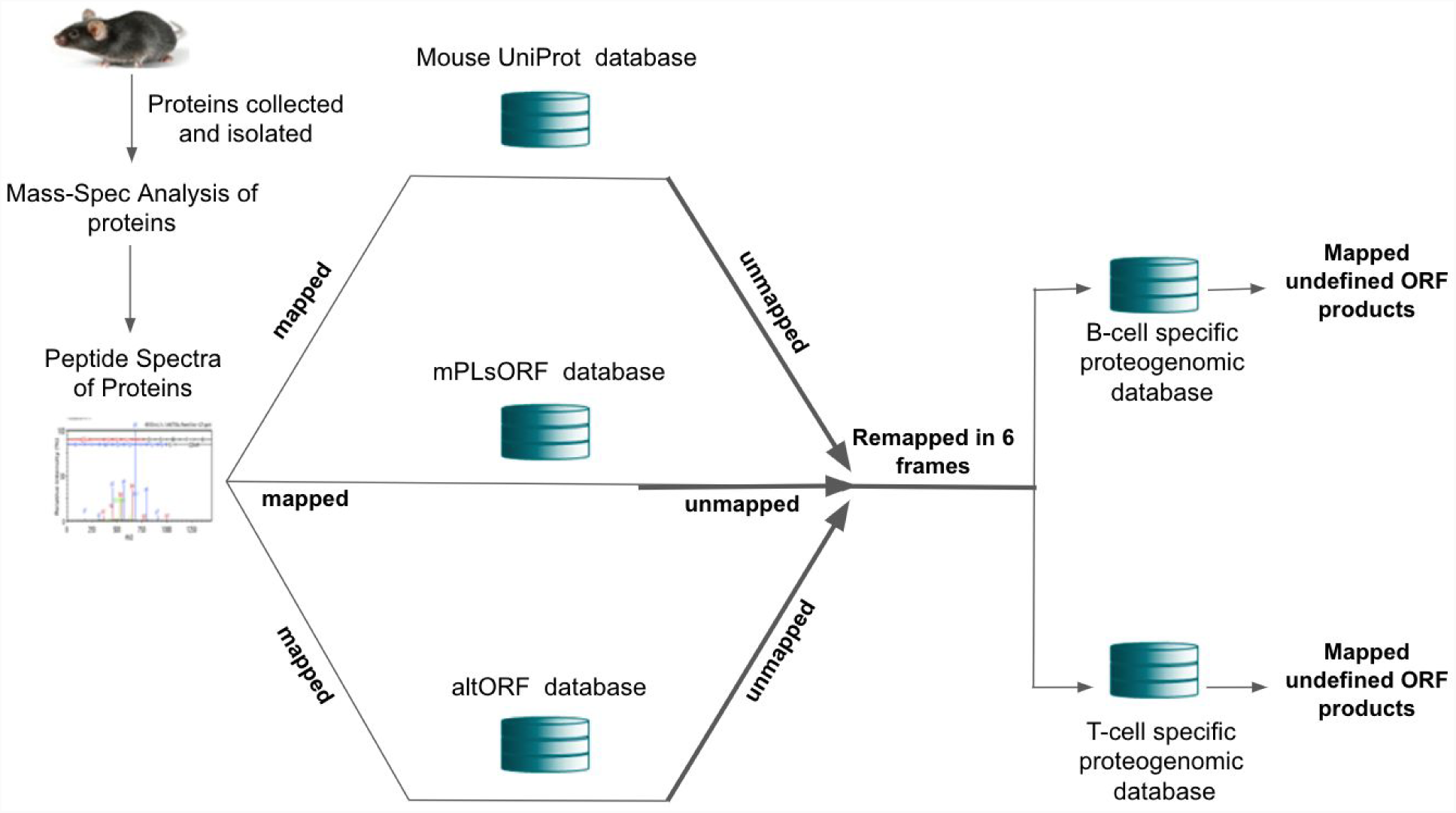
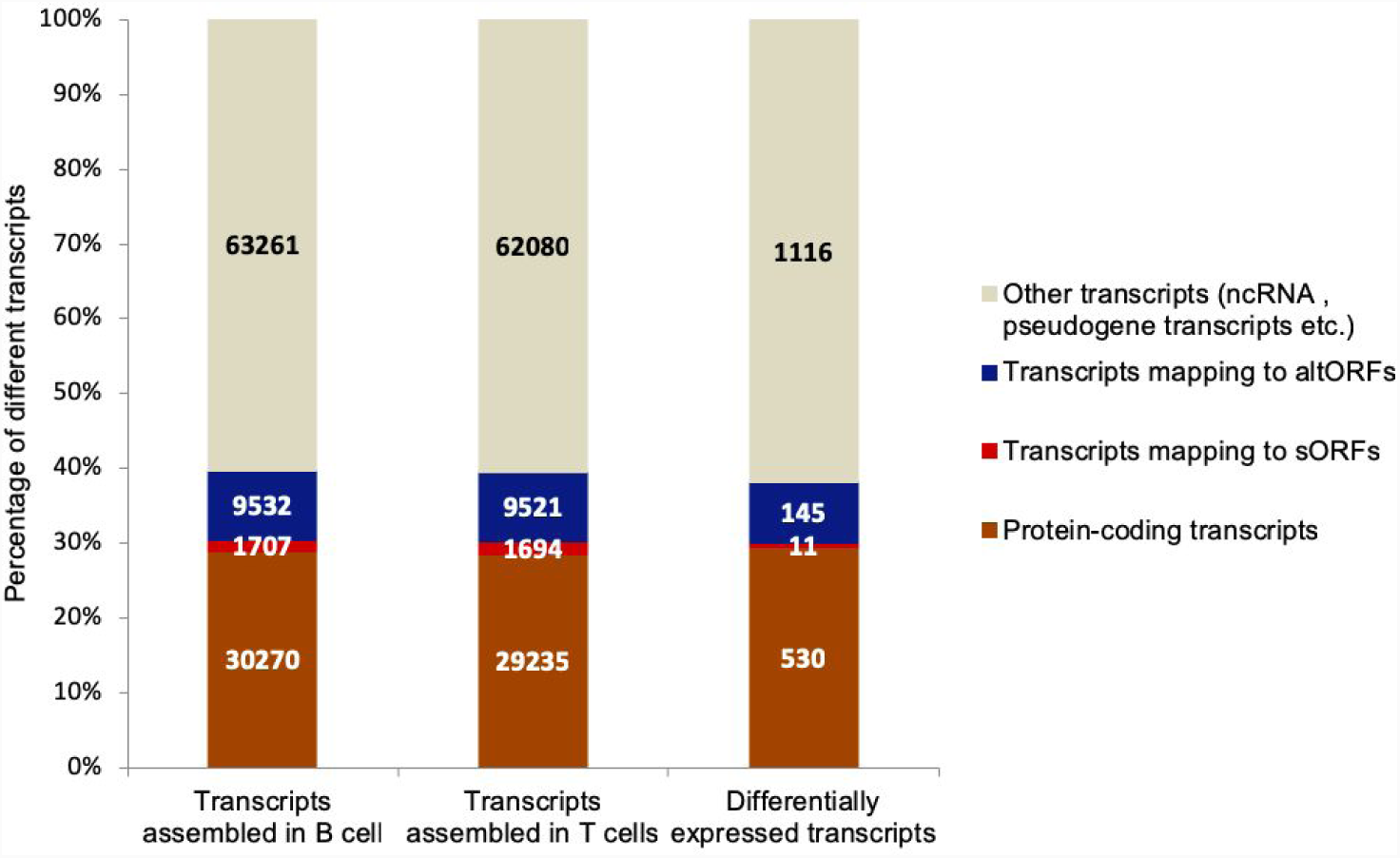
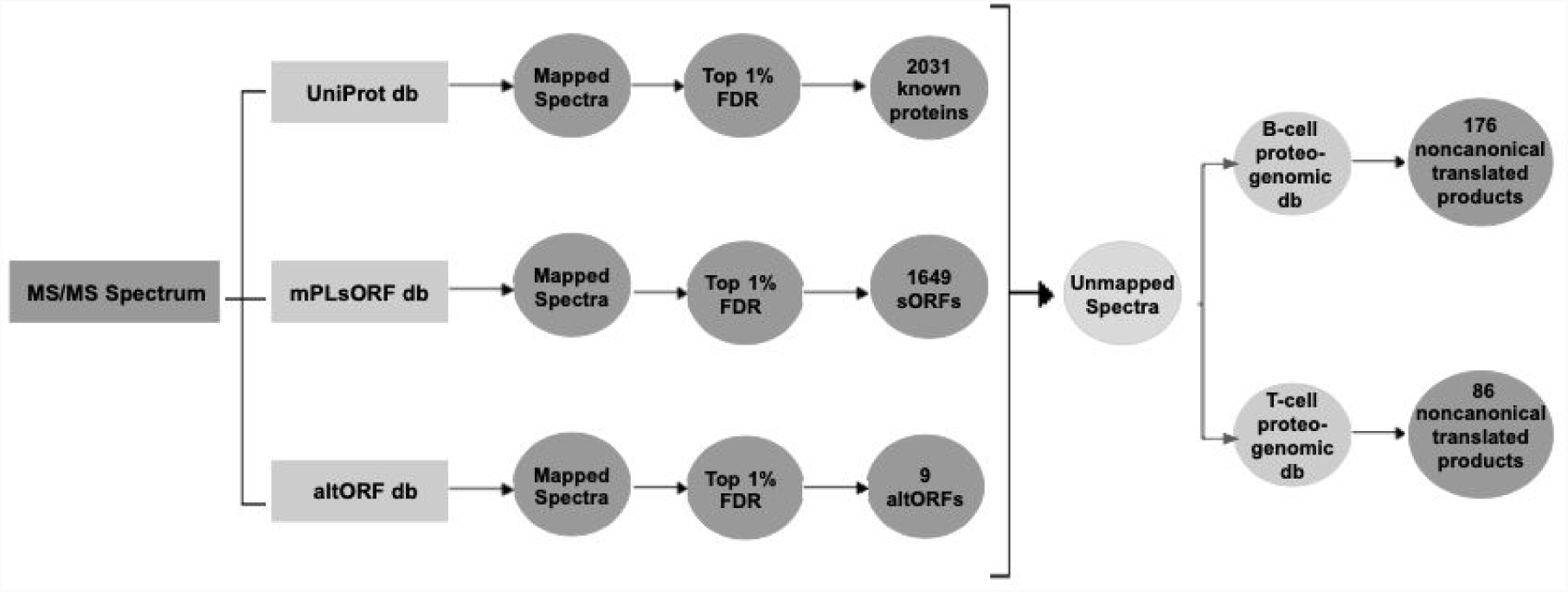
Proteogenomic workflow to identify non canonical translated products in mouse B and T cells. **(A)** Illustrates the schematic workflow of our proteogenomic analysis. Briefly, mass spectra of proteins obtained from mouse B and T cells were independently mapped to (a) mouse UniProt database, (b) mPLsORF database, and (c) the mouse altORF database. Unmapped peptides were remapped to a B and T cell-specific proteogenomic nucleotide database to identify any other undefined ORFs. (**B**) Different categories of transcripts, expressed in percentage (y-axis), identified from different transcript datasets (x-axis). **(C)** Schematically illustrates the results from the proteogenomic analysis. Briefly, 2,031 known proteins, 9 altORFs, 1,649 sORFs, and 259 undefined novel ORF translated products were identified in mouse B and T cells.

As putative novel ORFs, sORFs, and altORFs have been identified by other research groups [10,11,14], we investigated whether we could first identify any established sORF and altORFs in our mouse B and T cells data. We collated all known mouse sORFs from sORFs.org [11] and SmProt [10] containing 1,127,154 and 15,581 mouse sORFs, processed them, and retained only those that had the strongest evidence for coding in our sORF database. The database finally contained a total of 454,120 sORFs assigned with a unique sORF identifier (Fig. S10). We downloaded mouse altORF coordinates from the Roucou laboratory [14] and after processing obtained a final list of 215,320 altORFs (Fig. S11). For more description on how these databases were created please see material and methods. Using these two databases, we identified 2,595 unique sORFs as transcribed in B cells and 2,535 unique sORFs as transcribed in T cells (Fig. S12). Similarly, we identified 3,007 unique altORFs as transcribed B cells and 2,953 unique altORFs as transcribed in T cells (Fig. S12). The identity of ∼62,600 transcripts from B and T cell transcriptomes could not be ascertained because they did not map to protein coding genes, sORFs, or altORFs. Although we could identify some of them as noncoding RNAs and pseudogenes, for the rest we could not identify any known genomic annotation (Fig 2B)

One thousand seven hundred and sixty seven transcripts were found to be differentially expressed (DE) between B and T cells using Ballgown [15]. Of these, 1,378 transcripts were common to both B and T cells (Fig. S13), 204 DE transcripts were unique to B cells and 185 DE transcripts were unique to T cells. Of note, a total of 530 DE transcripts are already known protein coding transcripts (**Fig. 2B**), of these, 443 were common to both B and T cells, 43 were unique to B cells and 44 were unique to T cells. Out of 1,767 DE transcripts 11 were sORF transcripts, and 145 were altORF transcripts (**Fig. 2B**). Furthermore, we identified one DE sORF transcript in T and 10 in both B and T, and 7 altORF DE transcripts in T, 9 in B and 129 in both B and T cells (Fig. S13).

### Identification of novel ORFs with translational evidence

To identify novel ORFs with translational evidence we used mass spectrometry with the following schema (for more details see material and methods). All mass spectra obtained from naive B and T cell proteome were mapped to the following three databases independently: (a) Uniprot database, which has fasta sequences of all known mouse proteins, and our in-house (b) mouse sORF and (c) mouse altORF databases (**Fig. 2A**). This analysis identified that 2,031 known proteins, 1,649 sORFs and 9 altORFs were translated (**Fig. 2C**). Mass spectra that did not match to any of the three databases and whose identity was unknown were further evaluated by mapping them to the B and T cell-specific nucleotide databases in six frames. From this analysis 259 non canonical transcript regions (176 in B cells and 86 in T cells) were identified to be translated with at least two peptides matching per non-canonical transcript, (**Fig. 2C**). A total of 766 peptides were used to identify 259 non canonical transcript regions as translated. Because the two or three peptides that mapped to each of these 259 regions were separated by vast distances and because we did not have peptide evidence for the intervening distances we decided to identify ‘undefined ORFs’ within these non canonical transcripts based on single peptides. For this we located start and stop codons upstream and downstream of these peptides in these non canonical transcript regions. This resulted in 835 ‘undefined ORFs’ among these 259 non canonical transcript regions.

### Combining novel ORFs with transcriptional and translational evidence

Fig. S14 shows the distribution of transcript abundances of sORFs, altORFs, known proteins, and undefined novel ORFs. Fig. S15 shows the distribution of protein abundances of exonic sORF proteins, non exonic sORF proteins, altORF proteins, and known proteins.

Venn diagrams of known protein coding genes and proteins identified in B and T cells are displayed in **Fig. 3A** (left and right panel). The majority of known protein coding genes and proteins are identified in both cell types. Among the novel ORFs, we found a total of 4,325 sORFs with evidence of transcription or translation, 45 sORFs had evidence for both transcription and translation and (**Fig. 3B**). Genomic annotations of 2,916 sORFs showed that the most abundant sORF annotations are lncRNAs (∼45%) followed by exonic (∼28%) and 5’UTRs (∼23%) (**Fig. 3B**). For altORFs we found 3,205 unique altORFs with evidence of either transcription or translation (**Fig. 3C**), 2 altORFs had evidence for both transcription and translation. Genomic annotations of 2,621 out of 3,205 altORFs revealed that 91% of altORFs are from known protein-coding regions, 6% from noncoding, and 3% from pseudogenes (**Fig. 3C**). Of the 259 undefined novel ORF regions a potential 1,405 with both transcriptional and translational evidence, 65% were from introns of known genes, 4% are found within or overlapping the CDS of a known gene, 25% mapped to antisense of known genes, and the remaining 6% are from non coding regions, pseudogenes, intergenic regions and 3’ UTR (**Fig. 3D**).

**Fig. 3.**
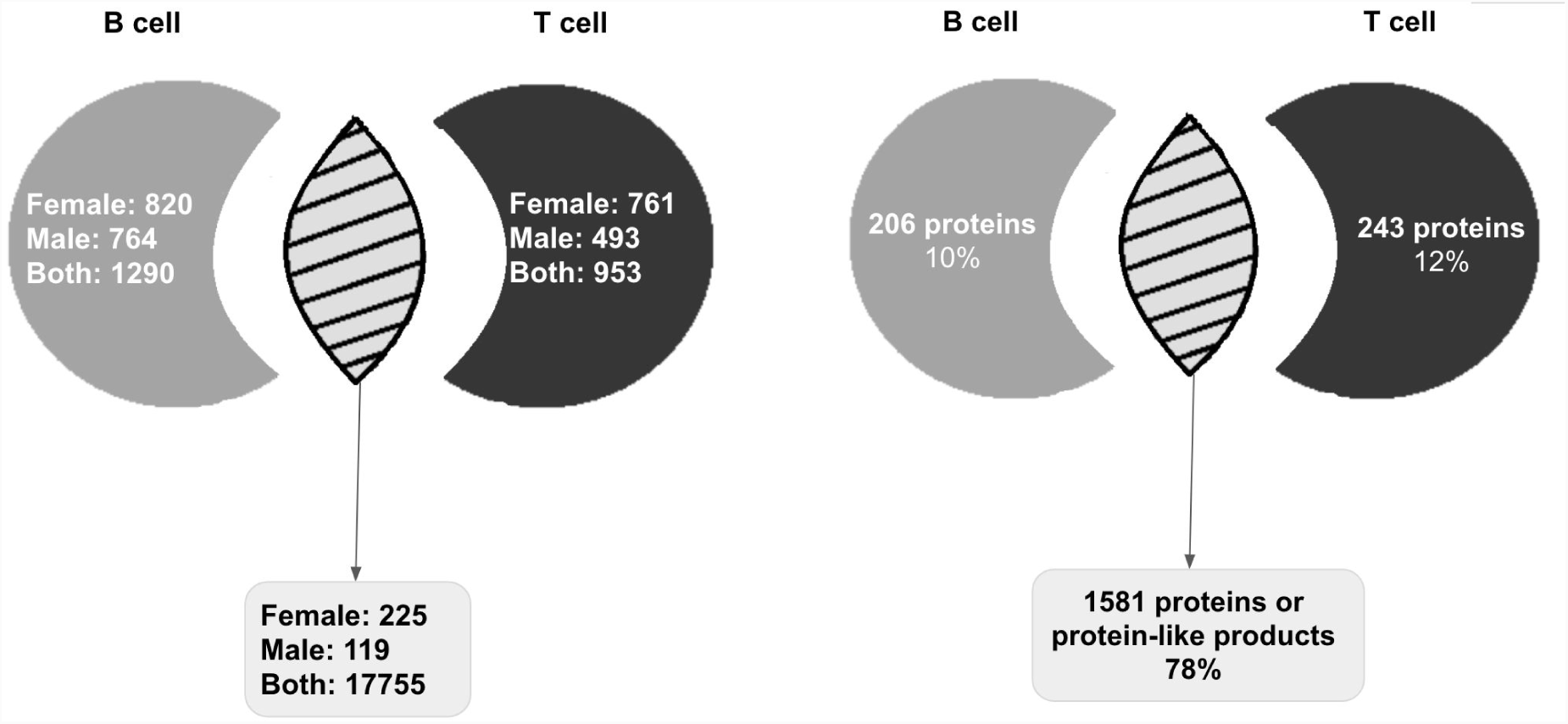
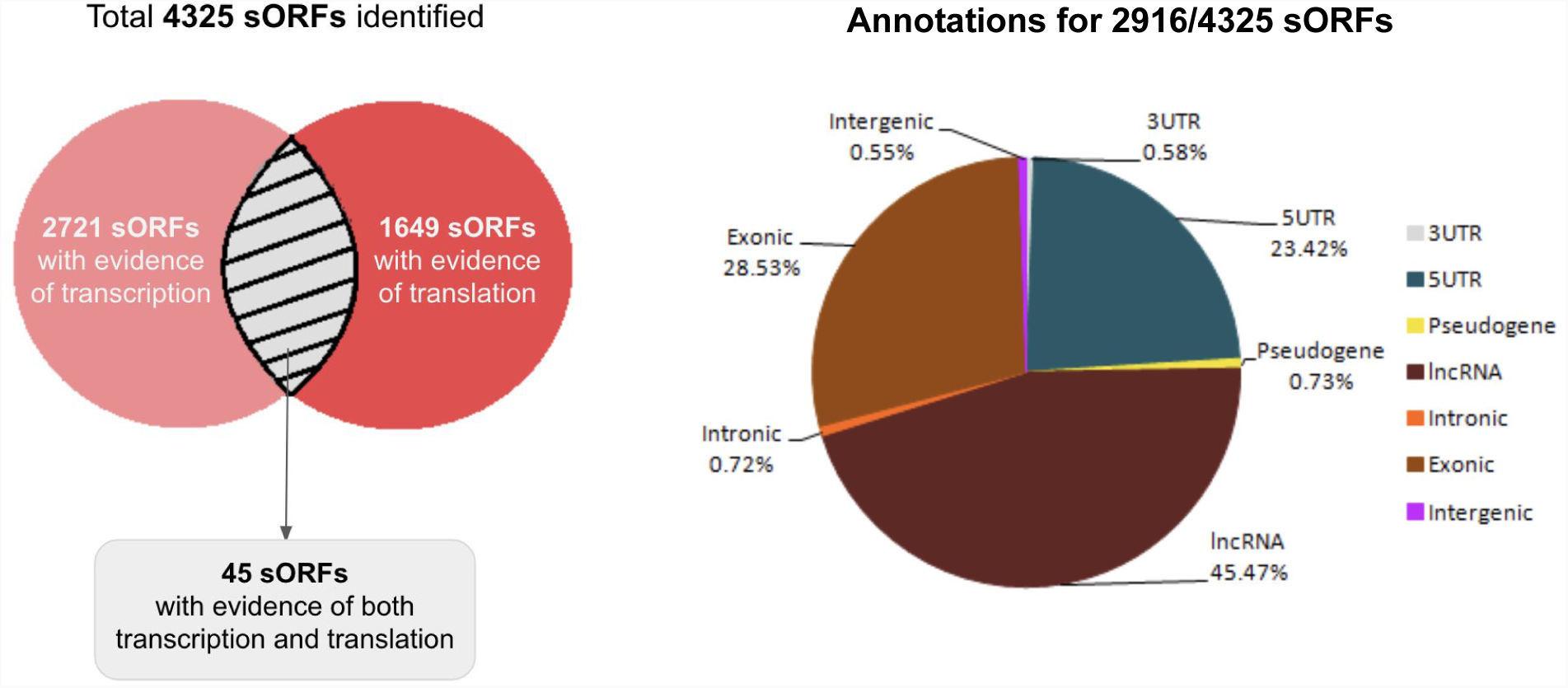
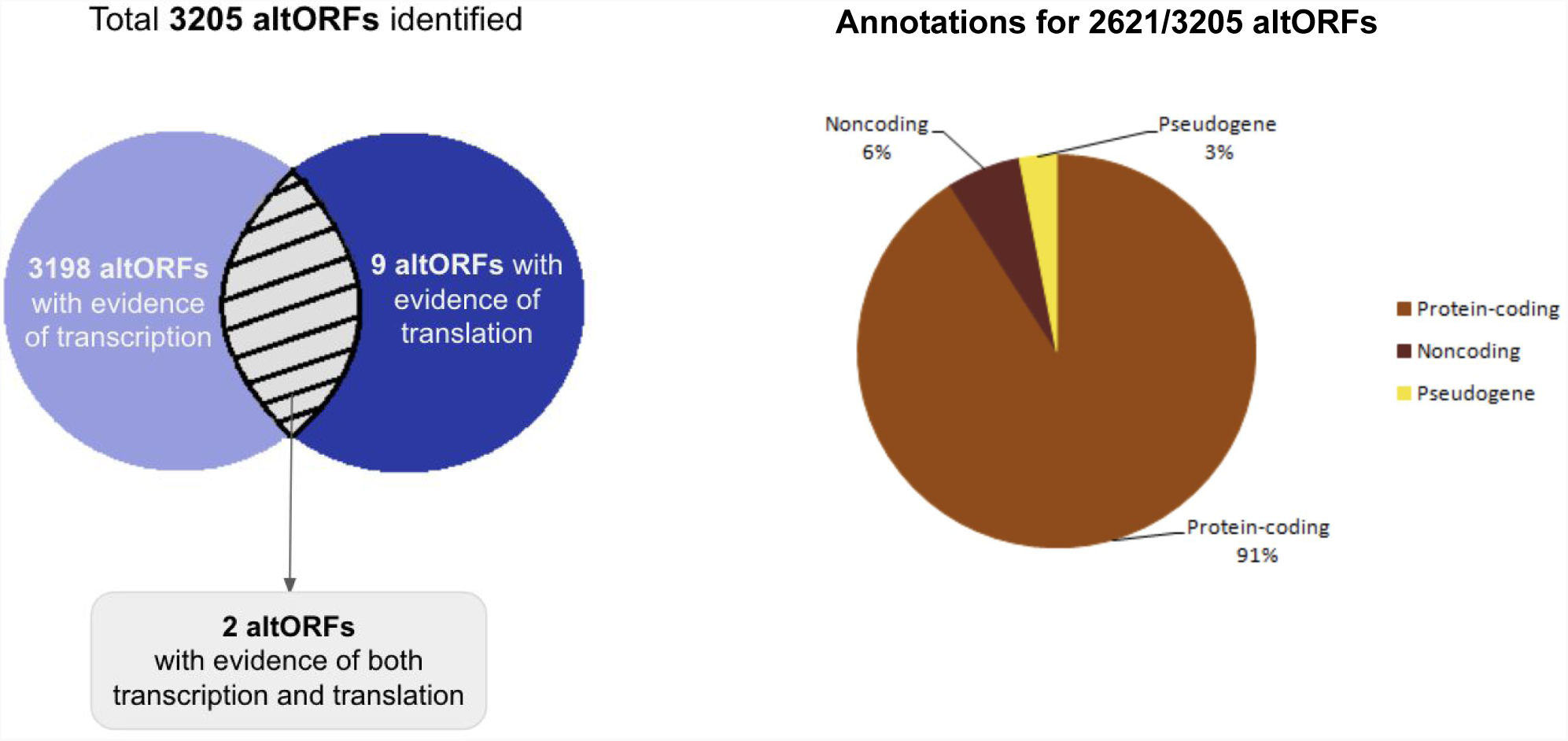
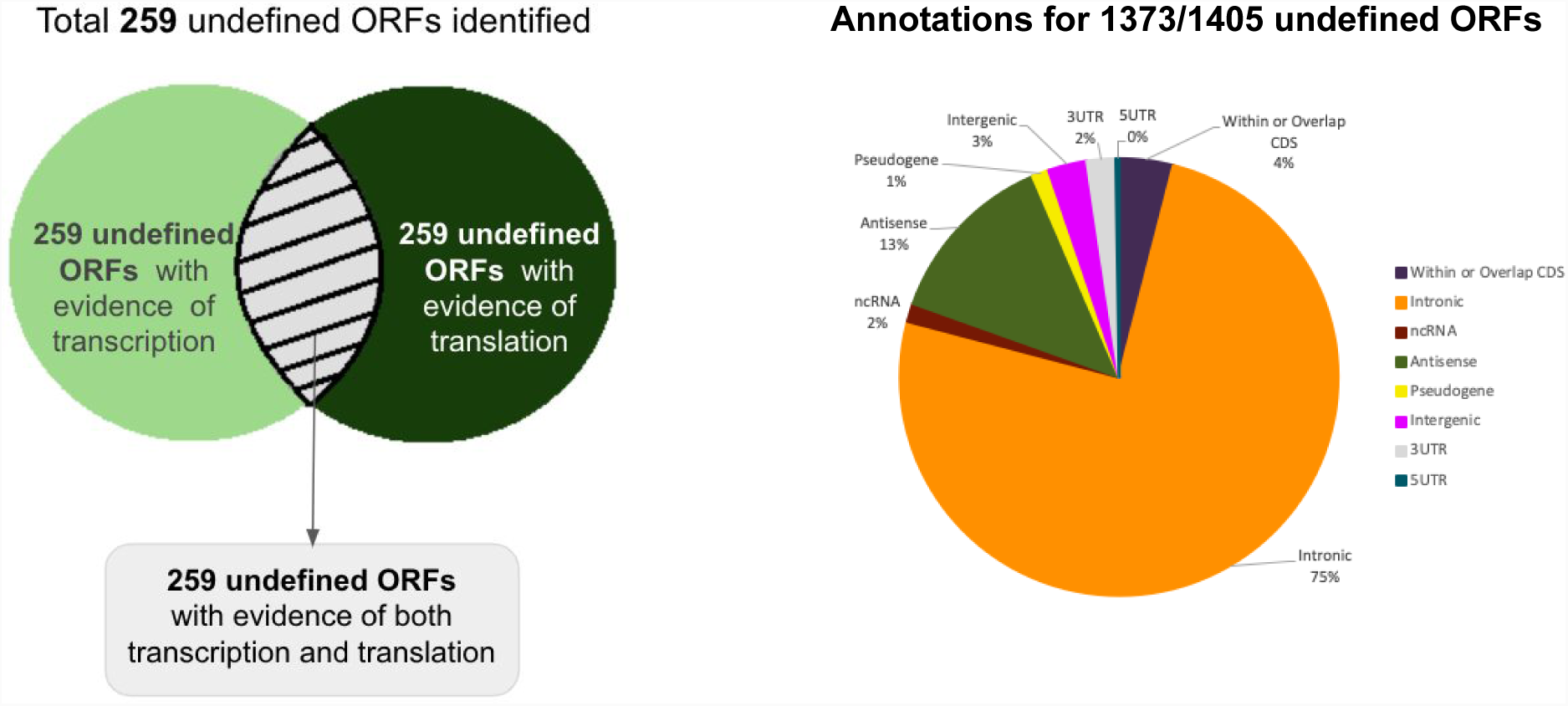
Identification of transcribed and translated products from known and novel ORFs in mouse B and T cells. (**A**) (Left panel) Cell and gender-specific categorisation of known genes based on expression of their transcripts in at least one specific cell type. (Right panel) Cell-specific categorisation of the known proteins. (**B**) (Left panel) A total of 4,325 sORFs in mouse B and T cells were identified to be either transcribed or translated. Of this, 2,721 sORFs show evidence of transcription, 1649 show evidence of translation, and 45 sORFs have evidence for both transcription and translation in our samples. Right panel shows a pie chart with the different genomic annotations for 2916/4325 sORFs. (**C**) (Left panel) A total of 3,205 altORFs in mouse B and T cells were identified to be either transcribed or translated. Of this, 3,198 altORFs display evidence of transcription, 9 of translation and 2 altORFs show evidence of both transcription and translation in our samples. Right panel shows a pie chart with the different genomic annotations for 2,621/3,205 altORFs (**D**) A total of 259 undefined ORFs in mouse B and T cells (Left panel) showed evidence of both transcription and translation. Right panel shows a pie chart with the different genomic annotations for 1,373/1,405 undefined novel ORFs is shown on the right.

We next demonstrated that the transcript abundances of protein coding genes, sORFs, altORFs, and undefined novel ORFs indicated that transcript abundance of novel ORFs are sufficient to distinguish B and T cell types like the known protein coding transcripts upon application of Principal Component Analysis (PCA) (Fig. S16). This observation suggests that transcriptional products of novel ORFs are unlikely to be a result of stochastic biological events.

### Investigating potential biological functions of novel ORFs

To infer whether the expression of novel ORF products could explain the cellular differences between B and T cells, we first checked whether the known protein coding transcripts and proteins could explain cellular differences. For this, Gene Ontology (GO) enrichment analysis was performed on transcripts and proteins that are uniquely expressed in either B or T cells, and transcripts that are differentially expressed in B and T cells. Fig. S17-20 show the GO-Slim biological process categories that are significantly enriched or depleted more than two-fold, in B-cells (grey bars) and in T-cells (black bars). Immune related functions are indeed enriched in both the known protein coding transcripts and proteins that are uniquely expressed in each cell type as well as in transcripts that are differentially expressed in both the cell types. But these uniquely expressed and differentially expressed protein coding transcripts and proteins are not sufficient to explain the cellular complexities and functions of B and T cells. Therefore, we hypothesized that the novel ORF encoded transcriptional and translational products may be involved in cellular functions that might contribute to the unique identities of B and T cells.

We performed the following analyses, (a) investigated whether the potential human orthologs of the identified novel ORFs are expressed in human tissues, (b) checked whether the expression of novel ORFs regulate the expression of known protein coding transcripts, (c) inferred the functions of novel ORFs from their sequences, (d) mapped mutations from COSMIC and HGMD databases to novel ORFs to check whether they harbour deleterious mutations, and (e) finally we investigated whether the translated products could form protein-like structures using the structural genomic approach EV-fold [13], to infer potential biological functions of novel ORFs,

We first identified human orthologs of all the identified sORFs, altORFs, and the undefined novel ORFs and then their expression in whole blood samples in GTEx data [16] was investigated. Among all the conserved sORFs, altORFs and other undefined ORF products we found evidence for expression of only 20 sORFs in the GTEx data (**Fig. 4A**). This is because GTEx data is predominantly based on poly A based RNA sequencing methodology and majority novel ORF transcripts may not have poly A and hence may not be sequenced and represented in GTEx. Abundance of the 20 sORFs, and the first quartile, median and third quartile expression values of all known protein coding transcripts are plotted in **Fig. 4A**. Six sORFs are expressed more than the median value of known protein coding transcripts, while two sORFs are very highly expressed and 12 sORFs are lowly expressed. This indicates that sORFs are expressed in human tissues and their expression levels are comparable to known protein coding transcripts and hence may have potential biological functions and can definitely not be ignored.

**Fig. 4.**
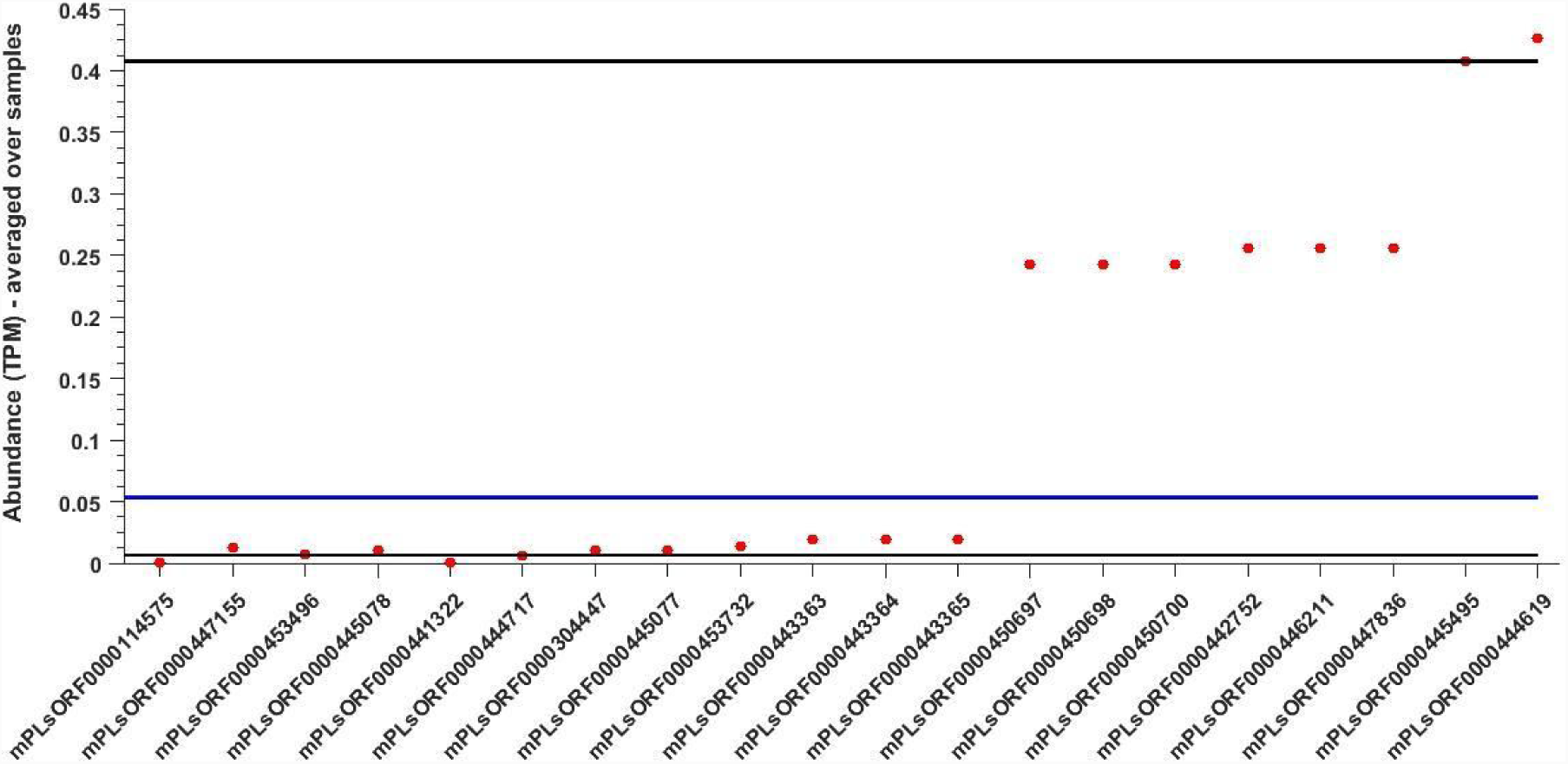
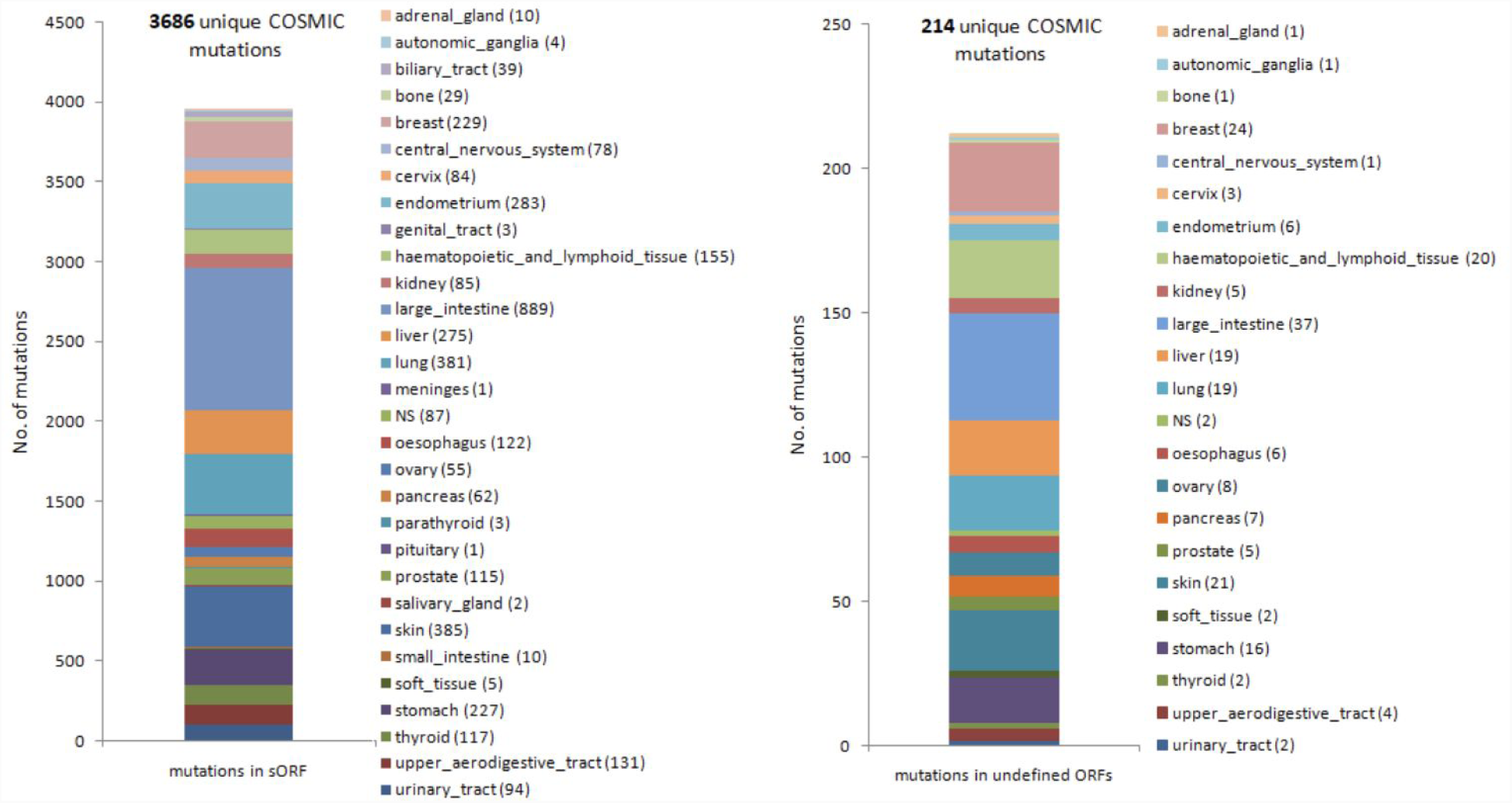
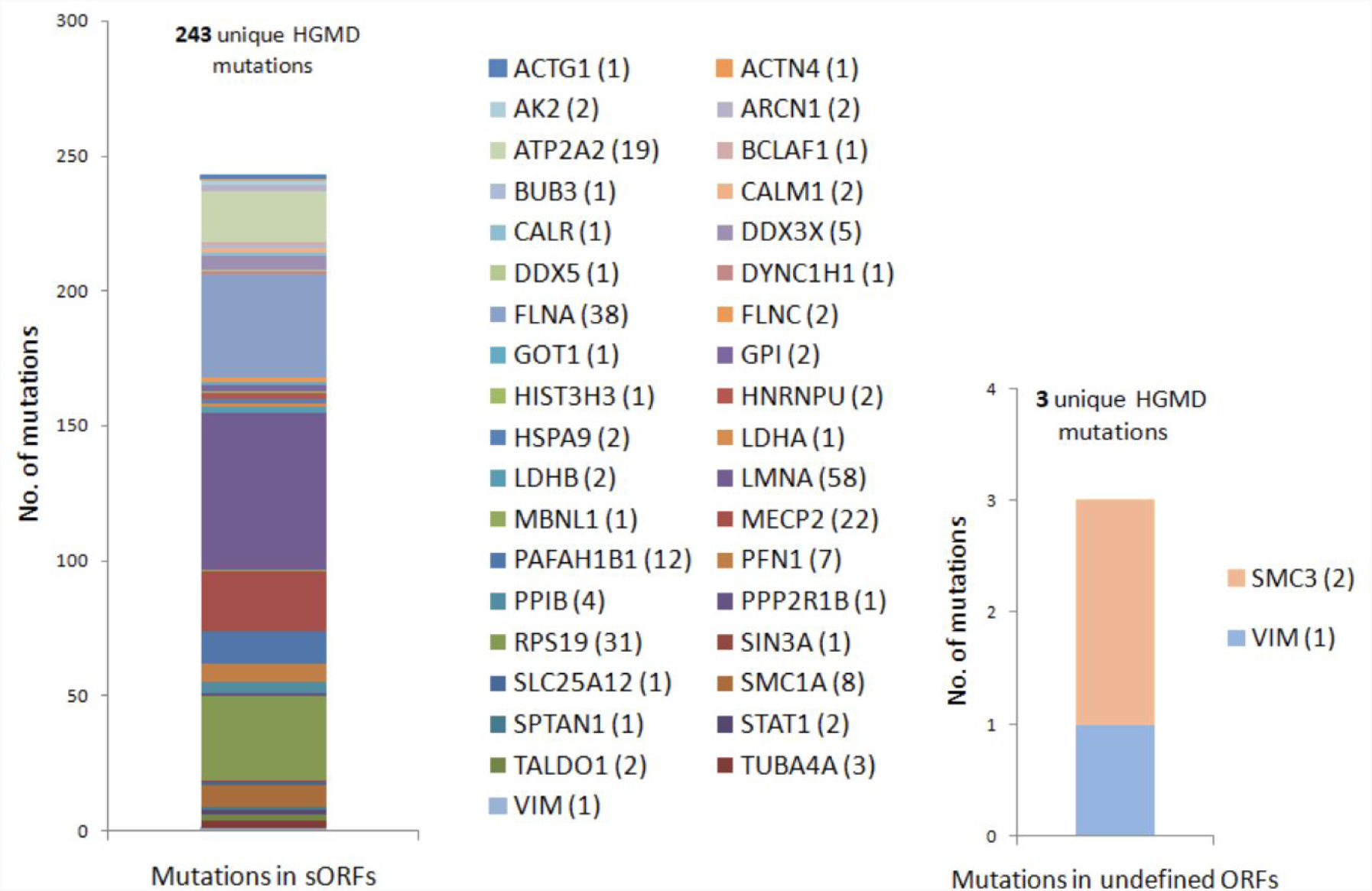
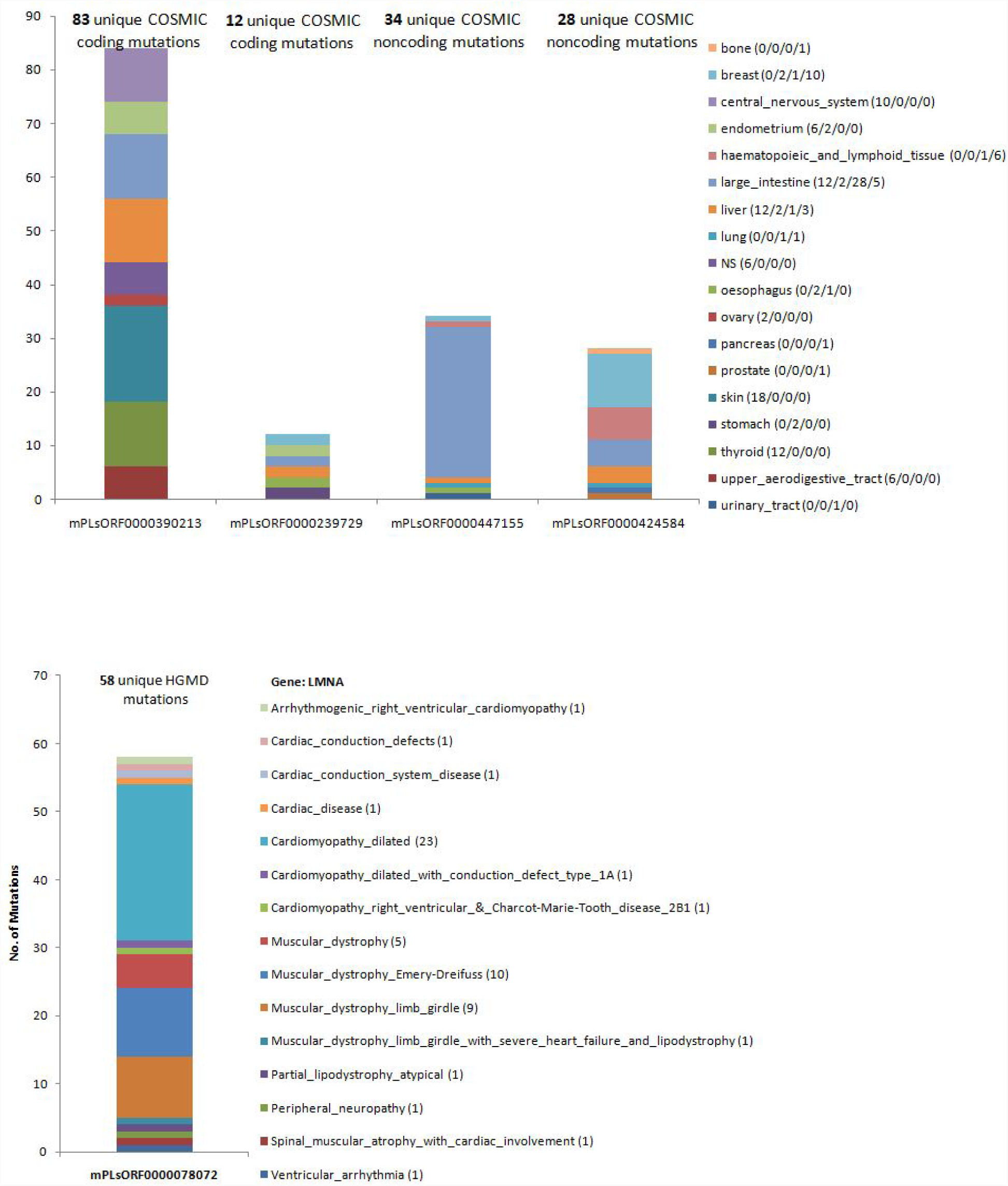
Investigation of biological functions of novel ORFs conserved in humans. **(A)** TPM abundance of human orthologues of 20 mouse sORFs in whole blood samples (from GTEx) plotted in the y-axis. For each sORF, average TPM is calculated over 407 whole blood samples and again averaged over all transcripts. The horizontal solid lines, from bottom up, are the first (black), second (blue) and third (black) quartiles calculated for the average expression of all protein coding transcripts in whole blood. **(B)** Results of mapping COSMIC mutations to sORFs and undefined ORFs that are conserved in the human genome identified using tblastn and LiftOver. **(C)** Results of mapping HGMD mutations to sORFs and undefined ORFs that are conserved in the human genome identified using tblastn and LiftOver. Disease phenotypes and the number of mapped mutations associated with the genes in the legend are expanded in Table S4 **(D)** Examples of COSMIC (top panel) and HGMD (bottom panel) mutations mapping to four sORFs. Numbers in brackets in the legends indicate the number of mutations from a particular primary tissue mapping to different sORFs from left to right.

To explore whether the expression of a sORF or an altORF alters the expression of an overlapping known protein coding genes, we checked for positive or negative correlation between a sORF or an altORF expression and respective overlapping protein coding gene expression, surprisingly we did not observe any correlation (Fig. S21 and Fig. S22). We also observed that a nearby gene’s length is not correlated with the number of sORFs (Fig. S23) or altORFs (Fig. S24) overlapping it. A maximum of 17 unique sORFs overlapped to ‘*Lpp*’ gene with length 15,647 bp; whereas 16 sORFs were mapped to *Eef1b2* gene of length 1939 bp. A list of genes with more than 10 unique sORFs mapping to them is shown in Table S2. Similarly, 16 unique altORFs were observed to overlap with ‘*Gvin1*’ gene with length of 9,005 bp; whereas, a gene longer than 15000 bp had just 2 unique altORFs overlapping it. A list of genes with more than 7 unique altORFs mapping to them is shown in Table S3. There are 212 genes overlapping sORFs and altORFs and functional annotation analysis of these gene revealed there was no immune related functional category enrichment (Fig. S25) for these genes. The above analysis demonstrated that sORFs and altORFs do not regulate the expression of protein coding genes overlapping them and could have independent functions of their own.

After establishing that novel ORF expression may be relevant and the expressed products may have significant biological functions, we analyzed the sequences of novel ORFs using Interproscan to obtain GO terms to give us a clue to their putative functions (Fig S26). GO terms of known proteins with transcriptional and/or translational evidence were also analyzed to validate the Interpro predicted GO terms for novel ORFs. Expected values based on GO terms from known genes with cutoffs of q < 0.01 and p < 0.01 were used to determine significantly enriched or depleted GO terms for sORFs. Fig. S27 shows the list of significantly enriched or depleted GO terms for sORFs. We then used GOSim to cluster GO terms based on functional similarities between gene products and the associated GO terms (Fig. S28 and Fig. S29). The results indicate that sORFs are more involved in cytoskeletal or structural functions of the cells than signaling or protein binding functions. These analyses were not done for altORFs and other undefined ORFs due a small number of annotated GO terms.

Although the Interpro scan analysis indicated putative functional enrichment for all sORFs, we were not able to identify specific functions for the other novel ORFs. Therefore, we then looked for indirect evidence for functions of the novel ORFs. To do this, we identified their corresponding conserved novel ORF regions in the human genome and then mapped mutations from COSMIC and HGMD datasets to identify whether the novel ORFs are disrupted in diseases. **Fig. 4B** and **Fig. 4C**. respectively show the number of unique COSMIC and HGMD mutations along with the disease origin of these mutations for sORFs (left panel) and undefined ORFs (right panel) that are conserved in the human genome. **Fig. 4D** top panel shows examples of four sORFs with mutations from multiple cancer types. Maximum number of mutations were observed for the sORF mPLsORF0000390213, and sORFs mPLsORF0000447155 is expressed in whole blood (**Fig. 4A**). **Fig. 4D** bottom panel shows an example of sORF with multiple HGMD mutations. These results indicate that although we are not able to absolutely infer the functions of these novel ORF products, we can conclusively show that they harbour disease associated mutations.

Finally, we investigated whether the novel ORF translated products have the potential to form protein-like structures. The primary reason why this had been overlooked is because it is assumed that they are not big enough to form structures. We checked the length distribution of all the sORFs (Fig. S30) and altORFs (Fig. S31) that we curated from different sources as mentioned above and observed that this is significant to form protein-like structures.

We predicted structures using the EVFold pipeline [17] that we set up in a cloud environment. Out of 45 sORFs that we identified in mouse B and T cells with both transcriptional and translational evidence, we were able to predict structures for 25 of them (Table S5). Next, we mapped mutations from COSMIC and HGMD to assess whether these mutations could affect the structures of these sORFs. **Fig. 5A** (left panel and right panel) are examples of sORFs for which we have structures and for which mutations were mapped. The exact number of mutations for these sORFs are represented in **Fig. 4D**. It could be seen that mutations that map to these sORFs indeed compromise their structures. **Fig. 5B** is a structure of a transcribed altORF. The overlapping gene of the altORF is Leucine Rich Repeat Containing 58 (*Lrrc58*). The biological function of the parent protein is uncertain and no gene ontology terms were assigned by the Interproscan. **Fig. 5C** shows (a) predicted structure of a translated product from the undefined novel ORF in an intergenic region in chr 14, (b) predicted structure of an undefined novel ORF insertion in Rps3a1 ribosomal protein (cyan) with the inserted fragment (red), and (c) predicted structure of an undefined novel ORF product antisense to *Raet1*. All of the above novel ORFs are marked and represented in Integrative Genome Viewer (IGV) in Fig. S32-S35.

**Fig. 5.**
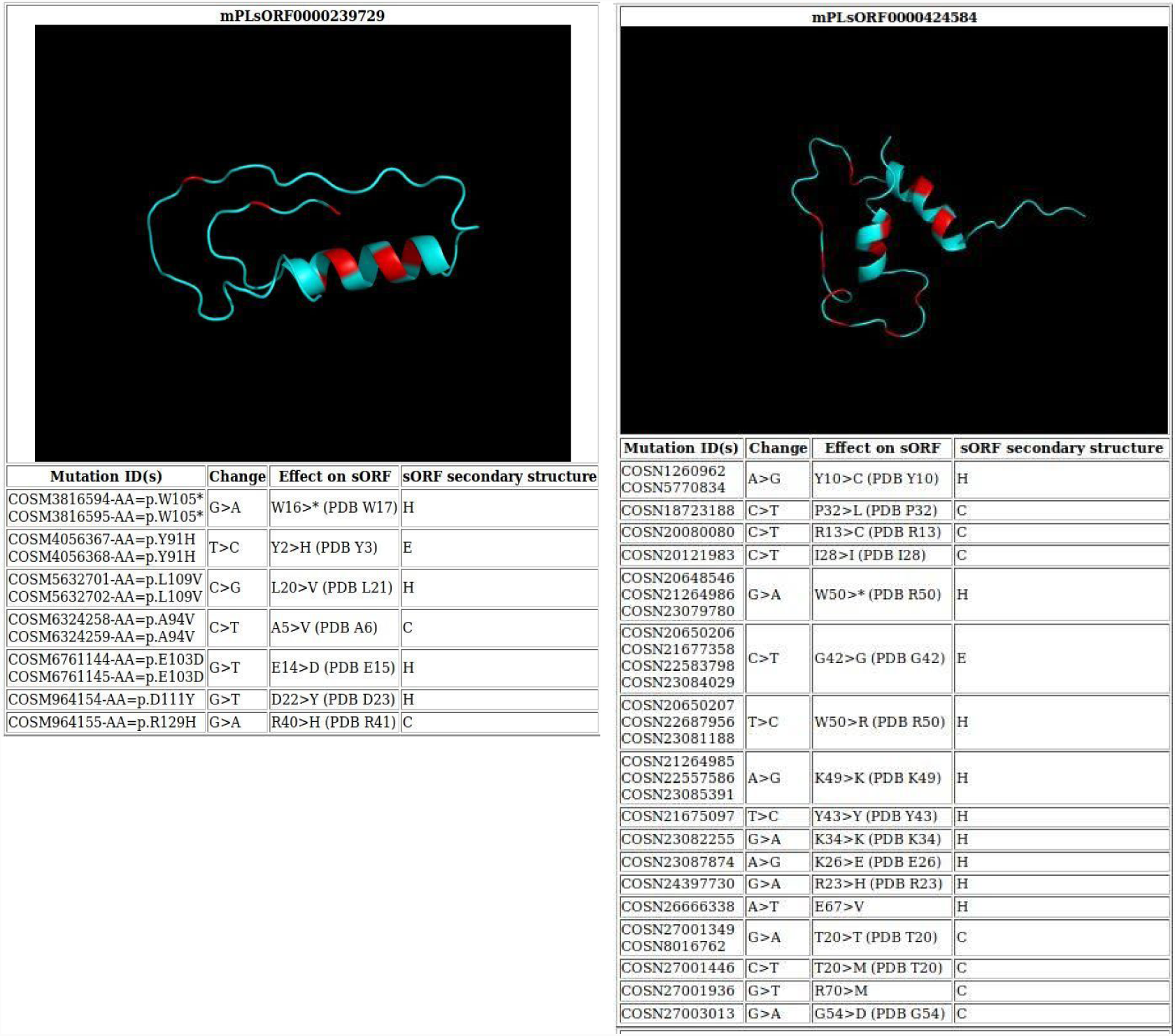
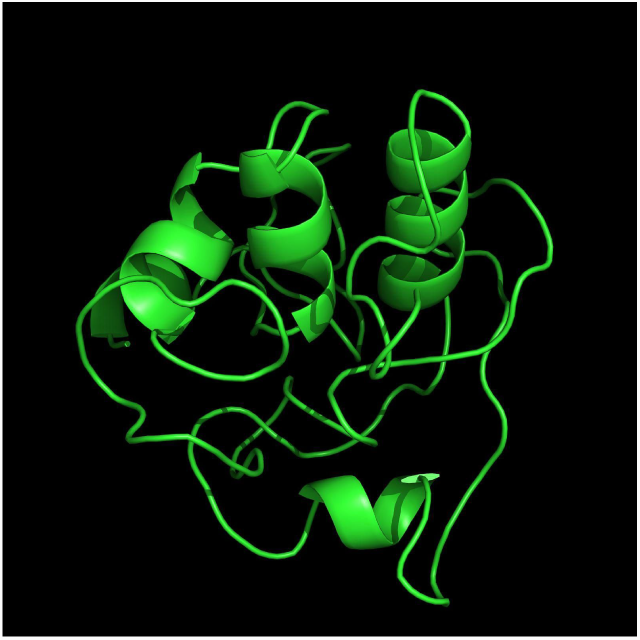
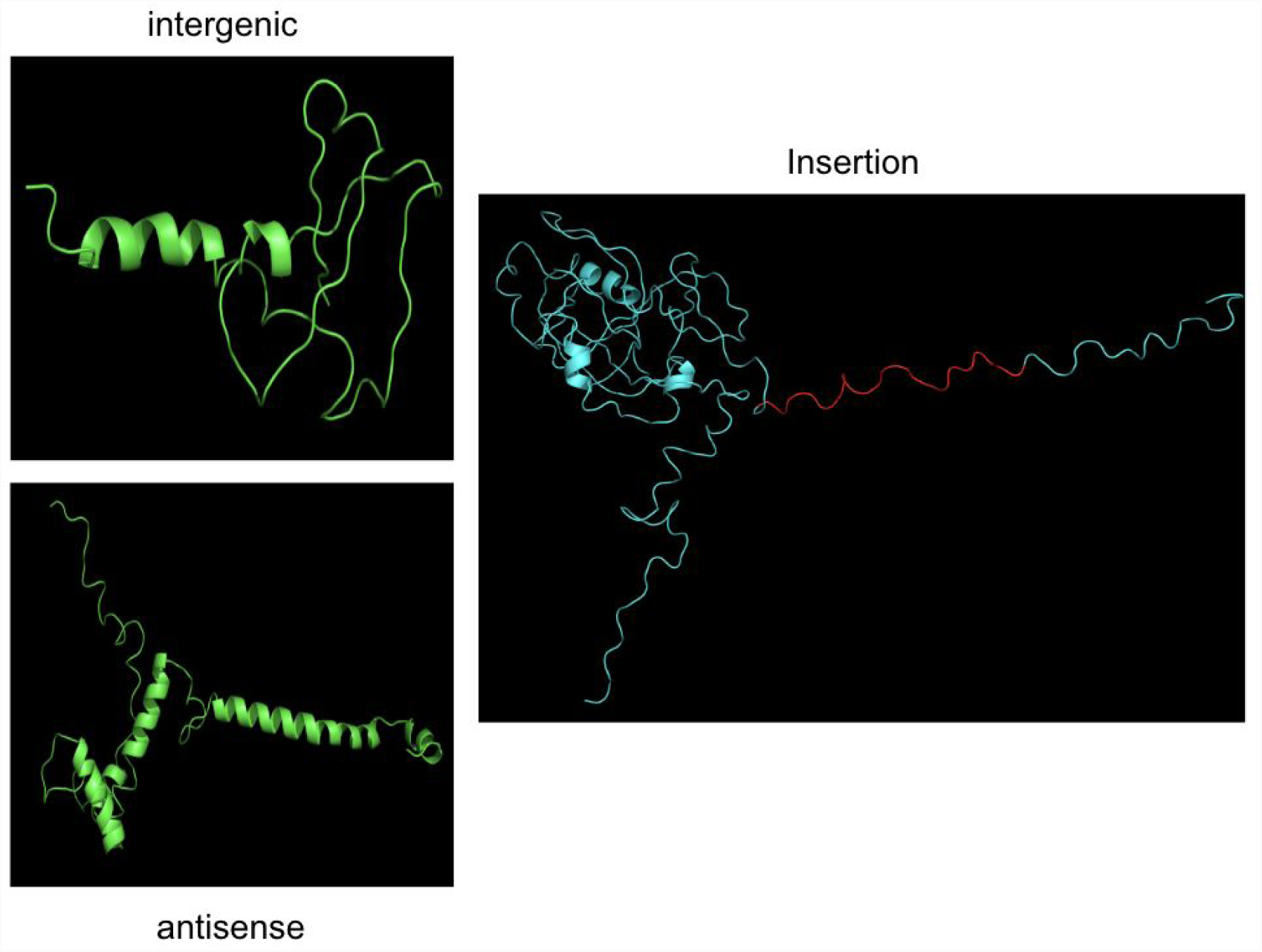
Potential structures of the novel ORFs from B and T cells. **(A left)** Predicted structure of mPLsORF0000239729 using EV-Fold displayed with pymol. Red regions on the structure indicated amino acids which are affected by COSMIC mutations. The table shows the list of cosmic mutation IDs along with the nucleic acid change, predicted amino acid change according to the standard amino acid code, and the predicted secondary structure of the protein at that position. H = Helix, C = Coil, E = Strand. **(A right)** Predicted structure of mPLsORF0000424584 using EV-Fold and as described for (A left). **(B)** Predicted structure of IP_233958.1 (species: Mus musculus) from the Roucou laboratory for which we have evidence of transcription. **(C)** Predicted structures of intergenic protein in chr 14 (left). The insertion ribosome_s40 (right) is the predicted structure of the original Rps3a1 protein (cyan) with the inserted fragment (red). The antisense protein (bottom) is the predicted structure of a predicted ORF antisense to *Raet1*.

## Conclusion

In this study we identified three types of novel ORFs; 4,325 sORFs, 3,205 altORFs, and 835 undefined novel ORFs with transcriptional and/or translational evidence in naive mouse B and T cells. We then investigated whether the transcribed and translated products from these novel ORFs had any potential functions by assessing whether the corresponding conserved novel ORFs in humans are expressed in whole blood cells. We identified 20 that are indeed expressed at an equivalent relative abundance to known protein coding genes. We found no correlation between their expression and that of their overlapping known protein coding genes.. Since the identified novel ORFs had molecular weight distributions similar to that of general proteome, we hypothesized that they might have independent functions of their own. Prediction of their potential functions determined enrichment for cytoskeletal pathways, while signaling pathways are depleted for sORFs. We also indirectly inferred potential functions of these novel ORFs by checking whether disease associated mutations from COSMIC and HGMD map to these regions. A total of 3,900 mutations from COSMIC and 246 mutations from HGMD mapped to these regions. Finally, we predicted structures for 24 sORFs (Table S5), 9 altORFs (Table S6), three undefined ORFs (**Fig. 5C**), and also mapped mutations to these structures and inferred that these novel ORF translational products can not only form protein-like products but that mutations mapped to them with potential to disrupt their putative functions.

The novel ORFs that we identified in this study, those that we identified from mouse neurons in our previous study [3], and those identified by others from yeasts [18] to humans [19] are located in long non coding RNAs, pseudogenes, 3’UTRs, 5’UTRs, in alternative reading frames to canonical protein coding regions, and other non-coding regions of the genome [3,20]. Among these the smallest ORF for which any known function is attributed is just six amino acids long and is in a 5’UTR. It regulates the expression of S-adenosylmethionine decarboxylase in response to polyamine levels [21]. More such investigations conducted in the recent past have indicated a diverse range of functions for sORFs. These include muscle regeneration [22,23], phagocytosis [24], DNA replication [25], cancer [26] and metabolism [27,28]. Despite these examples, the vast majority of novel ORF products have not been investigated rigorously and systematically because most of the past studies have focussed on only sORFs, which encode peptides that are smaller than 100 amino acids, that have been dismissed as functionally inconsequential. Systematically investigating these novel ORF regions is very important since here, we not only show that disease-and trait-associated mutations map to these regions but we also show that the pathogenicity scores of these mutations are more than the pathogenicity scores of mutations that map to known proteins, and that the translated products from novel ORF translated products can form protein-like structures.

## Acknowledgments

We thank Dr. Marco Chiapello former member of the Cambridge Center for Proteomics for help with the initial proteomic analysis; Tessa Bertozzi, for help with the B and T cells extraction from mice and Prof. Anne Ferguson-Smith for access to BLUEPRINT datasets. We would like to thank Seven Bridge Genomics (https://www.sevenbridges.com/) for letting us use their cloud platform. We would like to thank RosettaHuB (https://rosettahub.com) for helping us to build applications using Amazon Web Services. We would like to thank Dr. Chiatanya Athale and Dr. Girish Ratnaparkhi for letting us share their lab space in IISER-Pune.

## Author contributions

CE did the transcriptomic analysis, contributed to the mutation analysis, correlation analysis, proteogenomic analysis, and writing the manuscript. DC performed the structural genomic and functional analysis, contributed to the transcriptomic analysis, proteogenomic analysis, mutation analysis, and writing the manuscript. NM performed the mutation and pathogenicity analysis, contributed to the transcriptomic analysis and writing the manuscript. Sh. P did the correlation, whole blood expression, and GO analyses, contributed to writing the manuscript. RC performed the FACS and proteomic experiments and contributed to writing the manuscript. YU contributed to the proteogenomic analysis. AA and JN participated in the initial transcriptomic analysis. MTW contributed to the transcriptomic and proteomic analysis. CP contributed to the FACS analysis. KL contributed to the proteomic analysis. SP designed and supervised the project, analysed the data, and wrote the manuscript;

## Data and materials availability

Almost all processed data is in the main text or in the supplementary materials. Transcriptomic data can be obtained from GEO accession GSE94671 and GSE94676. Proteomics data will be deposited in PRIDE. All codes for this work can be obtained from https://github.com/PrabakaranGroup/t_and_b_cell_paper_scripts.

