## Supplementary material for "Translational products encoded by novel ORFs may form protein-like structures and have biological functions": Materials and Methods

### B and T cells total RNA sequencing data acquisition

B and T cells extracted from the spleen of six male and six female C57BL/6J mice were FACS analyzed to isolate resting B and naive CD4+ T cells. Total RNA was extracted from each of the 12 samples (three B-male, three B-female, three T-male and three T-female) and sequenced using Illumina HiSeq 2500. This work was done in Ferguson-Smith lab at the Department of Genetics, University of Cambridge. Data can be accessed at NCBI GEO database, accession GSE94671.

### Naive B and T cell separation for proteomics analysis

All steps were carried out fast, and cells were maintained in ice and ice cold buffers. Spleen from six male and six female C57BL/6J mice, age 12 weeks (Fig. S1), were collected in the cold 1x PBS (-Ca, -Mg) and gently crushed on a 40µM Nylon cell strainer (Fisherbrand, 22363547) using Iscove's Modified Dulbecco's Medium (IMDM, Sigma I3390), 10% heat-inactivated FBS (Sigma F9665), 1% antibiotic + antimicotic (Sigma A5955) media, 1% L-glutamine (Sigma G7513)) to isolate splenocytes. Samples were centrifuged for 5 min at 400 x g and the pellet was gently resuspended in 1x MojoSort buffer (Biolegend 480017). Cells were incubated with 2ml of RBC Lysis Buffer (Biolegend, 420301) for 3 min, and afterwards with 8ml IMDM-10. Cells were centrifuged for 5 min at 400 x g, resuspended in 1x MojoSort buffer to a final density 400µL of buffer per $10^{8}$ total cells. We obtained on average $1.8x10e^{8}$ splenocytes per spleen.

To cells, 360μL of antibody cocktail of biotin-conjugated monoclonal anti-mouse antibodies against CD8a, CD11b, CD11c, CD19, CD25, CD45R, CD49b, CD105, Ter-119, MHC class II, and TCRγ/δ from T cells isolation kit (Miltenyi,130-106-643) and cocktail of biotin-conjugated anti-mouse antibodies against CD43-Ly48, CD4-L3T4, and Ter-119 from B cells isolation kit (Miltenyi, 130-090-862) was added at a final concentration of 100μl biotin antibody per $10e^8$ cells and were incubated for 5 min in the refrigerator. Cells bound with biotin-antibody, 1.08ml of cold 1x MojoSort buffer and 720μl of α-Biotin microbeads were added (final 300μl buffer and 200μl microbeads per $10e^8$ cells) and incubated for additional 10 min in the refrigerator. Cells were then diluted with 15ml 1x mojo buffer and centrifuged for 5 min at 400 x g. Pellets were resuspended in 1.44ml 1X MojoSort buffer. Biotin-bound cells were depleted by passing through LD columns (Milteny, 130-042-901) in the magnetic field and flow-through was collected. Columns were washed twice with extra 1ml of buffer. Pooled unlabeled cells (total volume ~ 3.44ml per tube) represent the enriched T cells and B cells for subsequent FACS sorting.

Cells were counted as T Cells $2x10e^6$ cells per spleen, and B cells = $3x10e^7$ cells per spleen from $1.8x10e^8$ splenocytes. For sorting 100,000 cells per 100μl were prepared to establish the % population of naive T and B cells obtained from antibody-mediated sorts. To cells, 1μl antibody each of CD4-FITC, CD25-PE-Vio770 (PE-Cy7), CD44-APC, CD62L-PE was added for individual flow channels. Controls were prepared using 1μl of Streptavidin-V450, and a negative control with no antibody in cells. A sample was prepared with all above 5 antibodies. For B cells, Streptavidin-V450 and CD45R (B220)-PE antibodies were used in similar manner. Antibodies and cells were incubated on ice for 20 min, and then 850μl 1x mojosort buffer was added. Cells were centrifuged at 2000 rpm, for 5 min and resuspended in 500 μl buffer. Cells were processed through flow cytometer and following outputs were measured. We obtained 95% CD4+ cells that were live, single cells and streptavidin negative. CD25 channel filter removed 0.3% cells further, resulting in about 94% cells. A CD62+ and CD44+ gates resulted in 73 - 80% (CD4+ CD25- CD62+ CD44-) cells. For B cells, >99% of population processed through FACS sorter were selected for naive B cells (CD45R (B220)+). Samples were sorted with above settings using 1µl antibody per $1x10e^6$ cells, and cells were pelleted for proteomics workflow.

**Extraction of B and T cell proteome**

To extract total cellular proteome, cells were lysed in buffer (6M Urea, 2M Thiourea, 4% CHAPS, 5mM Magnesium Acetate, 30mM Tris pH 8.0), and 15µg protein in 5x Laemmli buffer with 5% b-mercaptoethanol was loaded on Mini-PROTEAN® TGX™ Precast Gels (BioRad). Gel lanes were cut into three sections for peptide extraction. Gel sections were cut into 1-2mm cubes, washed with 50% Acetonitrile and 100mM Ammonium bicarbonate solution until blue stain is washed. Gel pieces were treated with 100% Acetonitrile, and then reduced with 10mM DTT in 100mM Ammonium bicarbonate for reduction at 56℃ for 1 hour, and alkylated with 55mM Iodoacetamide in 100mM Ammonium bicarbonate in dark for 45 min at room temperature. Gel pieces were washed with 100mM Ammonium bicarbonate, and then treated with 50% Acetonitrile followed by 100% Acetonitrile. Subsequently, gel pieces were treated with diluted trypsin (5ng/ul) enzyme for overnight at 37℃. Peptides were extracted, dried, and dissolved in 3% Acetonitrile with 0.1% Formic Acid.

**Mass spectrometry analysis of the B and T cell proteome**

All LC-MS/MS experiments were performed using a Dionex Ultimate 3000 RSLC nanoUPLC (Thermo Fisher Scientific Inc, Waltham, MA, USA) system and a Q Exactive Orbitrap mass spectrometer (Thermo Fisher Scientific Inc, Waltham, MA, USA). Separation of peptides was performed by reverse-phase chromatography at a flow rate of 300 nL/min and a Thermo Scientific reverse-phase nano Easy-spray column (Thermo Scientific PepMap C18, 2microm particle size, 100A pore size, 75microm i.d. x 50cm length). Peptides were loaded onto a pre-column (Thermo Scientific PepMap 100 C18, 5microm particle size, 100A pore size, 300microm i.d. x 5mm length) from the Ultimate 3000 autosampler with 0.1% formic acid for 3 minutes at a flow rate of 10 microL/min. After this period, the column valve was switched to allow elution of peptides from the pre-column onto the analytical column. Solvent A was water + 0.1% formic acid and solvent B was 80% acetonitrile, 20% water + 0.1% formic acid. The linear gradient employed was 2-40% B in 30 minutes.

The LC elutant was sprayed into the mass spectrometer by means of an Easy-Spray source (Thermo Fisher Scientific Inc.). All m/z values of eluting ions were measured in an Orbitrap mass analyzer, set at a resolution of 70000 and was scanned between m/z 380-1500. Data dependent scans (Top 20) were employed to automatically isolate and generate fragment ions by higher energy collisional dissociation (HCD, NCE:25%) in the HCD collision cell and measurement of the resulting fragment ions was performed in the Orbitrap analyser, set at a resolution of 17500. Singly charged ions and ions with unassigned charge states were excluded from being selected for MS/MS and a dynamic exclusion window of 20 seconds was employed.

**Assembly and Analysis of B and T cell total RNA transcripts**

Quality of sequenced reads was determined using FastQC (Fig. S3). Primary assembly sequence and comprehensive gene annotation files for C57BL/6J, GENCODE release version M12, were used as the reference genome in our analysis. A genome index file to assist with read alignment was created using HISAT2-build, which extracts the exon and splice-site coordinates from the reference annotation. The paired-end sequenced reads where then aligned to the genome using HISAT2 run with default settings and the '–dta' option to ensure that strand information is retained after alignment. The output SAM files were converted to BAM format and the aligned reads were sorted based on genomic coordinates using Picard SortSam. Aligned reads in the BAM files and the reference genome were used to assemble sample-specific transcripts using StringTie run with default settings and the '–fr' option which assumes that reads were generated from a stranded library. The sensitivity and specificity of the StringtTie output relative to the full reference annotation and to a subset of protein-coding transcripts extracted from the reference annotation (defined by the "transcript_type "protein_coding"" tag) was assessed using GffCompare run using default settings and –T option to suppress output of mapping files.

The StringTie merge function was used to create a list of non-redundant transcripts in B and T cells using the 12 sample-specific GTF files. This merged transcript GTF file along with the 12 BAM files containing aligned reads were used for a second StringTie run with parameters '-Be' to calculate transcript FPKM values for each sample. Furthermore, we merged the information in the 12 CTAB files containing transcript FPKM values with the merged transcript file to create the final master transcriptomic file with ~164,000 transcripts. The master transcriptomic file was further analyzed as below.

We define unannotated transcripts in the master transcriptomic file as those without a corresponding gene name, and annotated/known transcripts as those which were assigned a reference gene by StringTie. Transcripts identified in unlocalised contigs in chromosome 1 and chromosome 4 with the names 'GL4XXXX' and 'JH5XXXX' respectively, were removed. Additionally, the master transcript file was filtered to remove transcripts with '0' FPKM values for all the 12 samples. This filtering gave us 109,441 transcripts. The remaining transcripts were categorised into four sub-groups: B-male, B-female, T-male or T-female, based on whether at least one out of three samples corresponding to a sub-group had a non-zero FPKM value. Finally, the transcripts were categorised into B or T cell-specific transcriptomic datasets based on whether a transcript was present in at least one of the two sub-groups corresponding to a particular cell type. This resulted in 101,767 B cell-specific transcriptomic dataset and 99,552 T cell-specific transcriptomic dataset.

**Creation of B and T cell-specific nucleotide proteogenomic database**

Transcript coordinates in the B and T cell-specific transcriptomic datasets were used to extract the corresponding nucleotide sequence from the reference genome using Bedtools Getfasta available in CGC. Bedtools Getfasta was run with default settings and with the name parameter = "True", which ensures that the name column of the input BED file is used as the header for the output FASTA file. Furthermore, transcripts with length > 100,000 nt were split into components of length less than 100,000 nt to facilitate downstream analysis using Mascot. The output FASTA files generated are our B and T cell-specific nucleotide proteogenomic database.

**Creation of sORF and altORF amino acid databases**

Prabakaran Lab mouse sORF (mPLsORF) database was created using information curated from two sources: sORFs.org [12] and SmProt [11]. sORFs.org contains 1,127,154 mouse sORFs, which have been either computationally predicted or experimentally verified. We exported mouse sORFs from sORFs.org with default filters except for FLOSS classification, which was set to 'GOOD' and 'EXTREME'. SmProt contains a list of computationally predicted small peptides identified in several species including mouse. We extracted 15,581 mouse sORFs from SmProt with filter parameters set to 'ALL'. The downloaded information from SmProt did not provide chromosome information for sORFs. A macros code was, therefore, run on the SmProt website to specifically extract chromosome information for sORFs.

Both databases had several duplicate entries which were removed by filtering them based on their chromosome location and amino acid sequence. We assigned unique sORF ids of the format 'mPLsORFXXXXXXXXXX', where X denotes a number, to each sORF entry and created our sORF database with the following columns: Organism_name, Source_database, Chromosome_number, Start_coordinate, End_coordinate, Strand, Amino_acid_sequence. There are still a few sORFs in our database with the same chromosome coordinates, but these duplicates were not removed because their corresponding amino acid sequences were different. Our final in house curated sORF database contains a total of 454,120 sORFs (Fig. S10).

We downloaded mouse altORF coordinates from Roucou's lab [13]. Few altORFs had multiple chromosome numbers assigned to it. These were removed from our dataset to generate a a final list of 2,15,320 altORFs (Fig. S11) for downstream analysis.

**Investigating sORF and altORF transcription in B and T cells**

GTF files for sORFs in mPLsORF database, altORFs in mouse altORF database and transcripts in our B and T cell-specific transcriptomic datasets were created and sorted first according to their chromosome number and then according to their start coordinates in ascending order. Bedtools intersect was used to identify coordinate overlap between sORFs or altORFs and B or T cell transcripts. The following parameters were used for this run: parameter '-f' = 0.99, signifying only sORFs/altORFs that overlap 99% of the transcript coordinates will be called; parameter -'wo' to generate information on the sORF or altORF, the transcript it matches to and the total number of nucleotide overlap between the two; parameter – 's' to only map sORFs to transcripts if they are from the same strand. altORFs were run twice once with and the other without the '–s' parameter because some altORF in the database do not contain strand information. altORF and sORF ids were extracted from the output TXT files generated by bedtools intersect and filtered to create a unique list of sORFs or altORFs with evidence of transcription. For sORFs, one sORF maps to multiple transcripts, and multiple sORFs map to one transcript. Similar cases are also observed for altORFs too.

**Proteogenomic workflow to identify sORF, altORF, and undefined novel ORFs**

Thermo raw mass spectrometry files were submitted to four databases search as described in Fig. 2a utilizing Proteome Discoverer v2.1 and Mascot 2.6. Briefly, an average of 383216 mass spectra were obtained from each sample. All mass spectra were initially searched independently against three amino acid databases - Uniprot database, sORF database, and altORF database and against the cRAP database of common contaminants. The spectra identification was performed with the following parameters: MS/MS mass tolerance was set to 0.8 Da, and the peptide mass tolerance set to 10ppm. The enzyme specificity was set to trypsin, and two missed cleavages were tolerated. Carbamidomethylation of cysteine was set as a fixed modification, whilst variable modifications consisted of: oxidation of methionine, phosphorylation of serine, theronine and tyrosine, and deamidation of asparagine and glutamine. High confidence peptide identifications were determined using Percolator node, where false discovery rate estimation (FDR) < 0.01. A minimum of two high confidence peptides per protein was required for identification.

Out of 383216 mass spectra, 165418 mass spectra mapped to Uniprot database; out of 383216 mass spectra, 67091 mass spectra mapped to sORF database; out of 383216 mass spectra, 32269 mass spectra mapped to altORF database. We then filtered the entries to remove 'cRAP', which are contaminants introduced during the experiment. Only those proteins with 'Medium/High' FDR values were retained. Finally, entries with no abundance values for all the four sub-groups were removed. After filtering for these parameters a total of 2031 known proteins, 1649 sORFs, and 9 altORFs were identified to be translated (**Fig. 2c**).

All unmatched mass spectra from each step were then exported, combined into a single mgf and duplicates were removed. B-cell specific mgf file contained 111,227 spectra and T-cell specific mgf contaniend 100,942 spectra. These files were then re-searched against B or T-cell specific nucleotide proteogenomic databases in six frames. 18,545 mapped to B-cell specific nucleotide proteogenomic database, 7,384 spectra mapped T-cell specific nucleotide proteogenomic database. Spectral matches were then filtered and validated by two independent approaches. The first validation was done with Mascot Decoy analysis in Mascot and a second independent validation was done with Percolator analysis in Proteome Discoverer (Thermo Scientific). Transcripts that were only identified by both the validation methodologies and with at least two peptides matching them were considered as translated. A total of 259 transcripts from both B and T cell nucleotide proteogenomic databases were identified to be translated with evidence of at least two peptides out of a total of 766 peptides mapping to them and these 259 regions were further analysed as discussed below

**Further processing proteogenomic results**

Of the 259 transcripts identified to be translated, 176 transcripts were identified in B cells and 86 transcripts were identified in T cells. These transcript regions varied in length, with the largest being 1.4 million bases, and because two peptides were separated by vast distances in single transcripts it was difficult to identify any undefined ORFs in this region. So, we decided to investigate undefined ORFs based on individual peptides within these transcripts. To do this, we aligned the peptide and searched the genome up and downstream of the peptide until a stop or start codon was encountered. Out of 766 peptides 689 peptides were unique and 632 peptides aligned mouse genome with evalue < 0.01. Of these 632 peptides we could annotate 617 peptides into 835 undefined novel ORF regions (**Fig. 3D**). The genomic coordinates of these undefined ORFs (+/- 500 bp up and downstream) were subsequently classified using Ensembl API (GET overlap/region) to identify neighbouring genomic features (genes, transcript, exon, cds) in the mm10 genome. A small portion of these ORFs could not be classified due to the genomic features from Ensembl disagreeing at different levels.

**Differential expression (DE) analysis of B and T cell transcripts**

The 12 CTAB outputs from the StringTie run with parameters '-Be', generated a list of sample-specific transcript FPKM values, which were used as inputs for DE analysis. Ballgown's 'stattest' function performs a $log_2$ transformation on the library-normalised FPKM values, fits the normalized values to a standard linear models and calculates p and q values for the transcripts. Here, transcripts with q values < 0.01 were called differentially expressed. Finally, the list of DE transcripts was filtered using Benjamin-Hoschberg corrected p-values at a cutoff of 0.05

DE transcripts that were identified after filtering were split into four categories: protein-coding transcripts, identified using transcript annotation information from Ensembl BioMart (Ensembl Biomart, Ensembl genes 91); sORF transcripts identified as DE transcripts that mapped to sORFs; altORF transcripts which are DE transcripts that mapped to altORFs; and finally, undefined transcripts that belonged to none of these three categories. DE sORFs and altORFs were categorised into B-male, B-female, T-male, T-female, B cells or T cells depending on their FPKM values.

**Principal Component Analysis (PCA) of B and T cell transcripts**

PCA to determine if sORF/altORF transcript abundances could distinguish between cell types and different genders of the same cell type. For this, we used FPKM values of transcripts that map to sORFs and altORFs for each of the 12 samples, $log_2$ normalised the data using a pseudo-count of 1 ($log_2$(number+1)) and centred the data around its median by subtracting the median from each data point. The final normalised dataset was then run using the R function prcomp with scale and centre parameters set to "FALSE". PCA plots for B vs T cells and male vs female cells were plotted along two principal component axes: PC1 and PC2. Similarly, PCA plots were generated for known transcripts.

**Gene Ontology (GO) Enrichments analysis**

To investigate whether just the transcriptome and proteome information is sufficient to differentiate between the B and T cells, enrichment analysis was performed using PANTHER Overrepresentation Test (release 2018-02-03) with annotation version 13.1. The genes expressed uniquely in a cell-type were compared to a reference list of all the genes expressed in that particular cell type. Similarly, proteins translated uniquely in a cell were compared against all the proteins expressed in that cell. Additionally, DE genes between B and T cells and also the genes that express both sORFs and altORFs were compared against all the genes expressed in both the cell types. Fisher's Exact test with false discovery rate correction was chosen for the enrichment of GO-Slim biological process categories. In all the analyses, genes with alternatively spliced transcripts; having transcripts expressed in more than one categorized cell type (B-male, B-female, T-male or T-female), was not considered for the gene ontology analysis.

**Correlation expression analysis of sORFs and altORFs with genes that overlap them**

The effect of expression of a sORF or an altORF on the expression of an overlapping known gene was checked by correlation expression analysis. Only those sORFs overlapping the exon, intron or an untranslated region of a known gene were considered in the analysis. Overlapping genes for altORFs were documented by the Roucou's lab. Whereas, overlapping genes of sORFs was determined by mapping the coordinates of the sORF to all the known genes coordinates in the mouse genome (Ensembl's Grcm38 annotation file). The gene within which the sORF transcript lies was chosen as an overlapping gene. Bedtools intersect with parameters '-s',' -f 0.99' and ' –wo' was used for coordinate mappings between the sORF transcript and the known gene. The abundance of the nearby or overlapping genes that are not expressed in B and T cells was assumed to be zero. Pearson correlation of abundances (FPKM) of all the sORFs/ altORFs to their respective overlapping gene was calculated. For the scatter plots the abundances were $\log_{10}$ normalized using a pseudo-count of 0.001 ($\log_{10}$(abundance + 0.001)) and the scale was adjusted to fit in the first quadrant.

**Correlation of length of a gene to the number of sORFs or altORFs overlapping with it**

The correlation of the gene's length to the number of unique ORFs overlapping with it was investigated. The length of the longest transcript of the gene was considered to be the gene's length. The longest transcripts length was determined by using a standalone python script GTFtools-0.6.5 [3] and by using Ensembl's Biomart [4].

**Expression analysis of sORFs in GTEx datasets**

We checked whether the mouse sORFs identified to be transcribed and/or translated in our naïve B and T samples are expressed in Whole Blood. The RNAseq data for the 407 Whole Blood samples collected from the deceased adult human donors was downloaded from GTEx. The RSEM quantified transcript abundances (TPMs) was used in the analysis.

We first identified the human orthologue transcripts of the mouse sORFs by mapping the sORFs transcripts against human genome (build GRCh38) using NCBI's tblastn with attributes -word_size 2. But, as the GTEX's RNASeq datasets are mapped against GRCh37 annotations, we used LiftOver to determine the corresponding human sORF coordinates in the GRCh37 based human genome.

Next, we checked the evidence of transcription of these sORFs in whole blood by mapping the sORF coordinates with the coordinates of all the transcripts expressed in the GTEx datasets using bedtools intersect with parameters -s, -f 0.99 and --w0. Of all the mouse sORFs transcribed and translated in our B and T cells, 20 were mapped to the GTEx transcripts. As tblastn maps a mouse sORFs to multiple human locations, for 7 of the 20 sORFs, each sORF was mapped to multiple transcripts from GTEx. The calculated abundance of each sORF is the expression value first averaged over the 407 whole blood samples, and then averaged over its respective transcripts (if any). Finally, we next checked the expression of these sORFs relative to the expression of all the known protein coding transcripts. Of the total 196520 transcripts expressed in all the tissues of the GTEx dataset, 145641 transcripts were annotated as 'protein-coding'. Three quartiles of the distribution of average abundance of each protein coding transcript in the Whole blood samples was calculated and plotted as in Fig. 4a.

**Structure Prediction of sORFs, altORFs, and translated products from undefined novel ORFs.**

EVFold pipeline was setup according to instructions on the github repository (https://github.com/debbiemarkslab/EVcouplings) on an Ubuntu AWS instance. This included the installation of the following software Hmmer suite 3.0, PLMC, CNS solve 1.2, HH-suite, Psipred, Maxcluster64. The database used was the recommended Uniref90 downloaded from https://www.uniprot.org/downloads.

**Mapping and visualisation of disease associated mutations in sORFs**

We developed computational strategies using bedtools intersect to map mutations from the HGMD and COSMIC database on to the protein and protein-like products from the noncoding regions. For that we had to first identify human homologous sequences. Briefly, LiftOver and NCBI tblastn, with attributes -word_size 2, was used to map mouse sORF to human genome, build Hg38. tblasn results further filtered using the following parameters constraints pident>80 & ppos>80 & ((mismatch*100)/qend) <10 & ((qstart*100)/qend) < 25 & qcovs > 80 & gaps <=2 & gapopen <=1).

LiftOver and tblastn mapped 4325 mouse sORFs to 1339 and 3429 regions for build hg38, respectively. Only 1339 regions for hg38 were mapped commonly from both LiftOver and tblastn. Hg38 mapped coordinates were scanned against Cosmic and HGMD variant databases using bedtools intersect without strand specification. Mapped mutations from each region were then compared to the coding sequence of each sORF to determine potential changes to amino acid sequence using Biopython. For the sORFs with predicted structures available, the mutations were mapped onto the PDB file and visualised with Pymol as red coloured residues.

**GO analysis of sORFs and altORFs against known Proteins**

Using interproscan-5.29-68 (downloaded from https://www.ebi.ac.uk/interpro/download.html) , we annotated sORFs and altORFs for which we have transcriptional and/or translational evidence in at least 1 sample. In order to allow a fair comparison, known proteins downloaded from uniprot with transcriptional and/or translational evidence were also annotated with interproscan serving as a reference point. Non automated annotation was not used as this information is not available for the majority of sORFs and altORFs. Proteins in the reference genome were referenced using uniprot accession IDs and the genes mapped to these IDs were obtained using the uniprot online mapping service (https://www.uniprot.org/mapping/). Analysis was performed on presence or absence of GO term annotation rather than the number of times the gene or protein might have been annotated with the same GO term.

Chi-squared tests were then performed with expected values based on the known protein proportions. The Bioconductor qvalue package was used to calculate q-values to be used for FDR correction. Cutoffs of $q < 0.01$ and $p < 0.01$ were used to select significantly enriched or depleted GO terms in sORFs. This analysis was not carried out for altORFs due to the low number of annotated GO terms. The significant GO terms were then clustered using the Bioconductor GOSim package using default settings of getTermSim.
