## Supplementary material for "Translational products encoded by novel ORFs may form protein-like structures and have biological functions": Figures

### Supplementary Figure 1

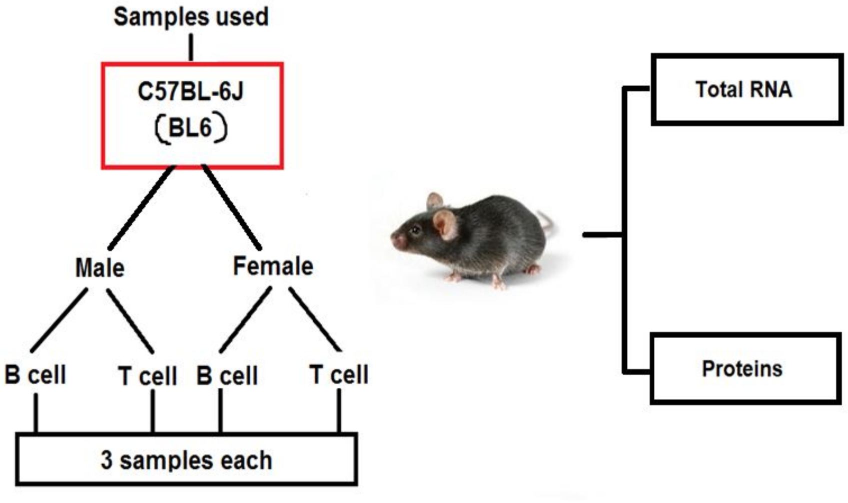


**Fig. S1. Extraction of total RNA and proteins from mouse B and T cells.** Naive B and T cells were isolated from spleen of two sets of six male and six female C57BL/6J mice that were 12 weeks old using FACS. From one set, total RNA was extracted from each of the 12 samples (three B-male, three B-female, three T-male and three T-female) and sequenced. From another set, proteins were extracted and proteins from the same sub-group (B-male, B-female, T-male or T-female) were pooled together for mass spectrometry analysis. Hence, for RNA there are three biological replicates; whereas, for proteins there is only one biological replicate.

**Supplementary Figure 2**

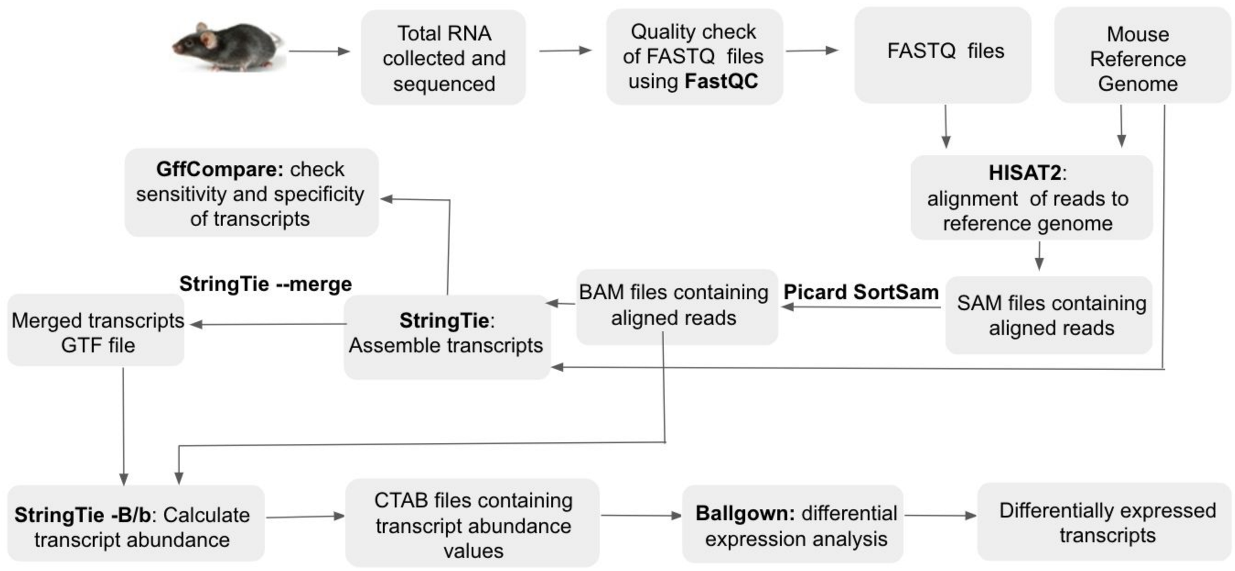


**Fig. S2. Workflow of transcript assembly and differential expression analysis.** Total RNA sequenced using Illumina HiSeq 2500 platform were assessed for their quality using FastQC. Read alignment was done using HISAT2, with FASTQ files and reference genome (GENCODE version M12) as inputs. The resultant SAM files containing the aligned reads were converted to BAM using Picard SortSam. Sample BAM files along with the reference genome were used as inputs for transcript assembly using StringTie, for which the quality was assessed using GffCompare. Transcripts assembled across the 12 samples were merged using the StringTie merge function for accurate transcript identification and downstream analysis. StringTie run with the -B/b parameter, using the sample BAM files and the merged transcript list as the reference genome, produced 12 CTAB files for each sample containing details of sample-specific transcript expression levels. These CTAB files were utilised for differential expression analysis using Ballgown. For further details pertaining to each step, refer to the materials and method section.

### Supplementary Figure 3

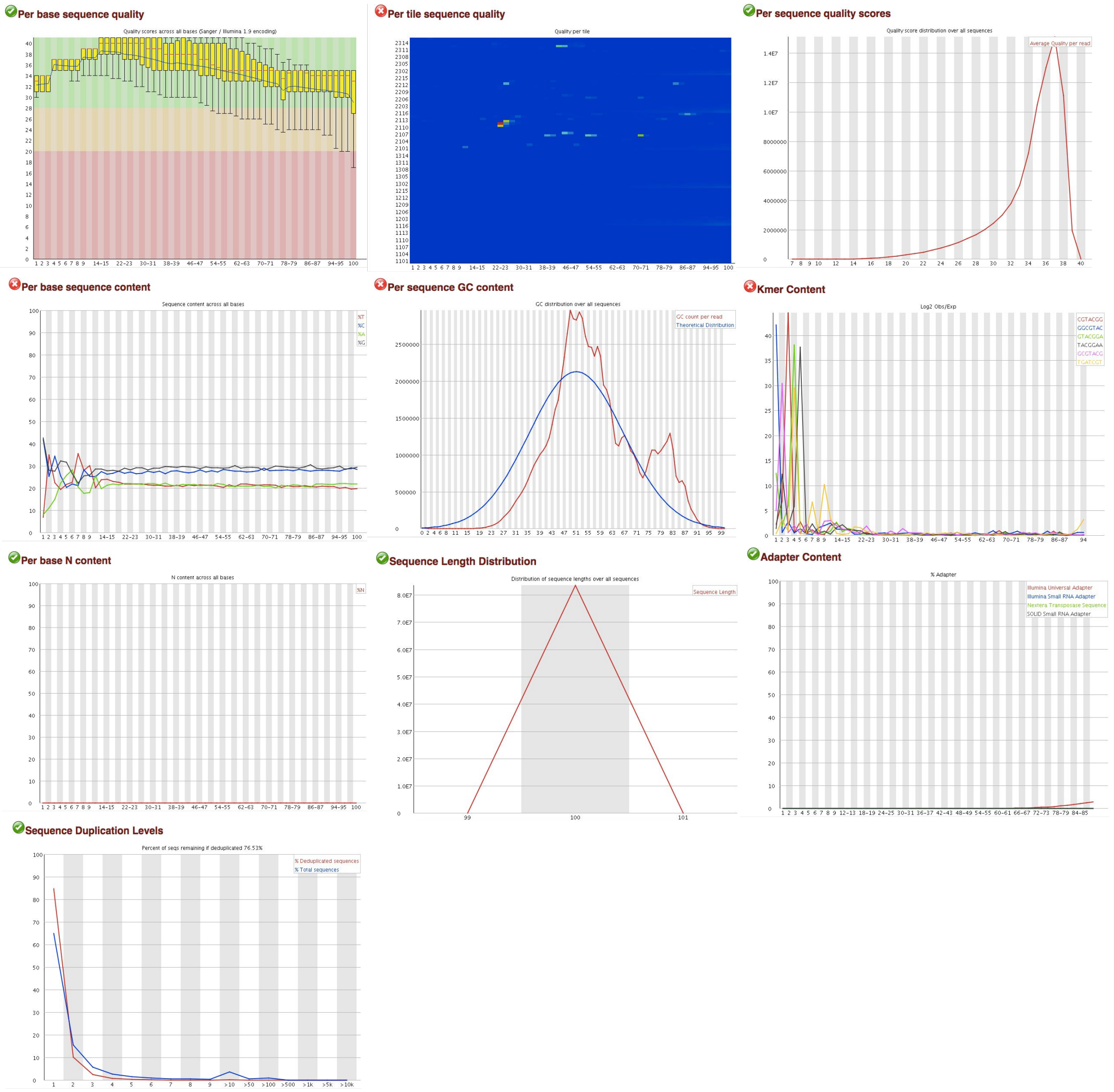


**Fig. S3. Representative raw read quality metrics generated by FastQC.** The 10 panels shown are representative FastQC report for the first set of reads for one of the female T cell sample. The ticks or crosses present in each frame represent how the data compare to the quality thresholds defined by FastQC (www.bioinformatics.babraham.ac.uk/projects/fastqc/Help) and do not necessarily imply that the data are inappropriate for analysis for the purposes of this project. In particular, the expected distribution of the per sequence GC content (blue line) does not refer specifically to the mouse transcriptome.

**Supplementary Figure 4**

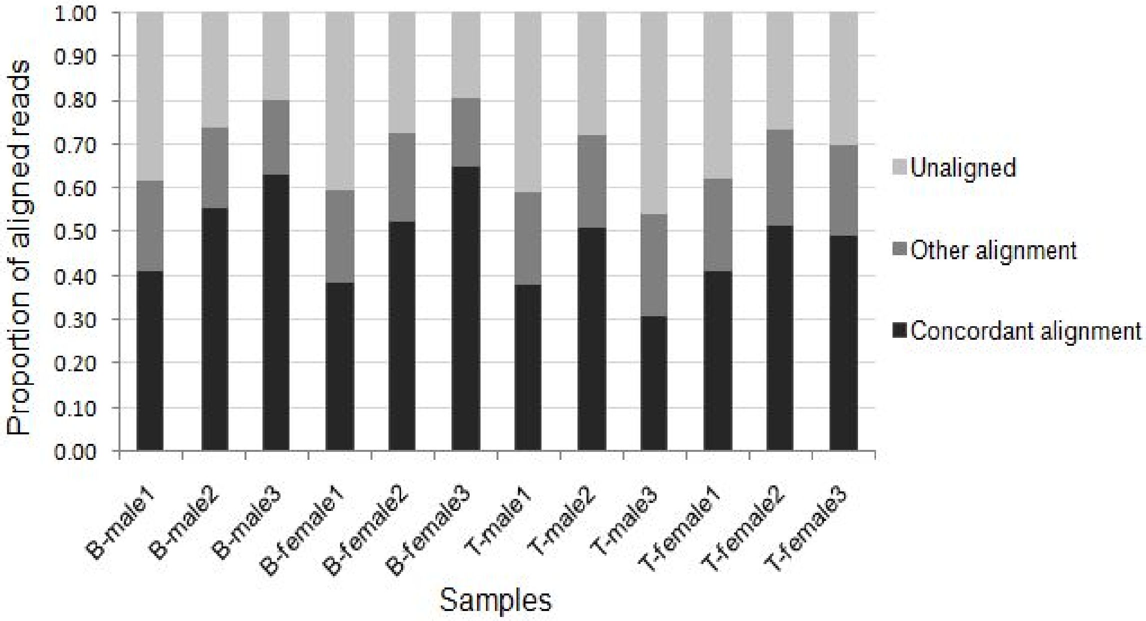


**Fig. S4. Results of HISAT2 read alignment.** The proportion of aligned reads (y-axis) corresponding to each of the 12 samples (x-axis) is represented by the bar graph. Read alignments were grouped into three categories: Concordant alignment (black) includes reads that aligned concordantly exactly one time; Other alignment (dark grey) includes reads that aligned concordantly more than one time, discordantly one time and where one of the mate pair aligned one or more times; unaligned (light grey) includes reads where none of the mate pairs aligned to the reference genome. On average ~68% alignment was achieved.

### Supplementary Figure 5

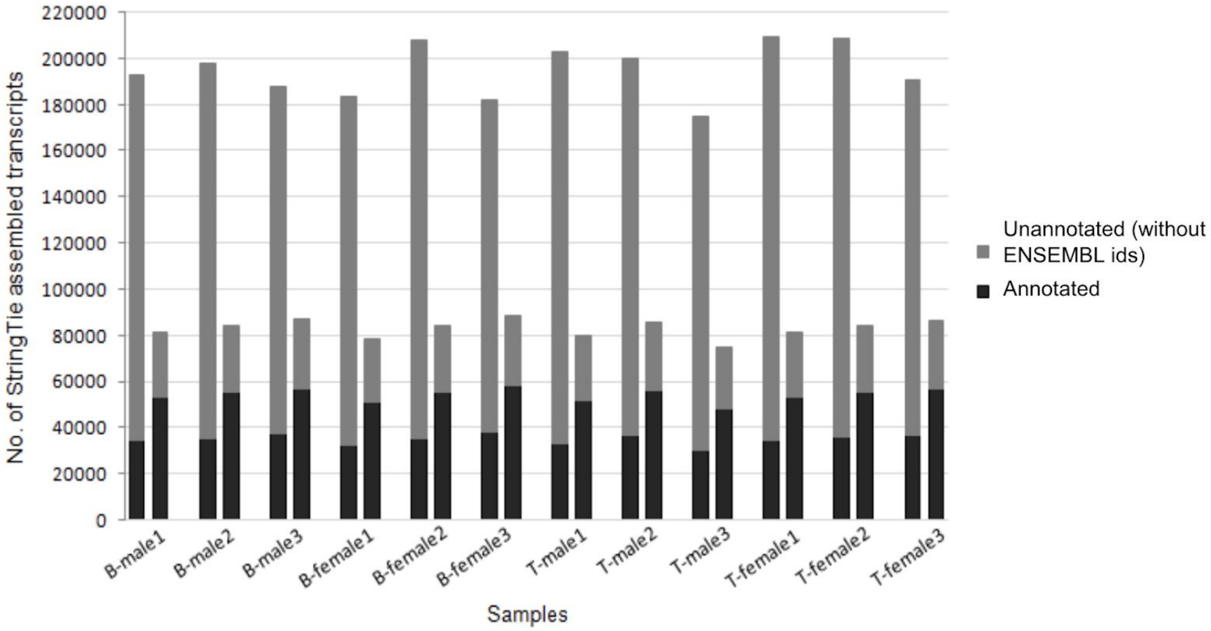


**Fig. S5. Results of transcript assembly using StringTie.** Number of assembled transcripts (y-axis) in each sample (x-axis) identified after two StringTie runs. Bars on the left for each sample correspond to transcripts identified after first StringTie run. Transcripts identified after applying the StringTie merge function is denoted by the bar on the right for each sample. Transcripts were called annotated (black) if StringTie assigned an ENSEMBL transcript id to the assembled transcript, else they were denoted as unannotated (dark grey). For each sample, a reduction in the number of total transcripts is observed after the merge step. Furthermore, there is an increase in the number of annotated transcripts and decrease in the number of unannotated transcripts indicating that several unannotated transcripts after the first StringTie run are incompletely assembled transcripts.

**Supplementary Figure 6**

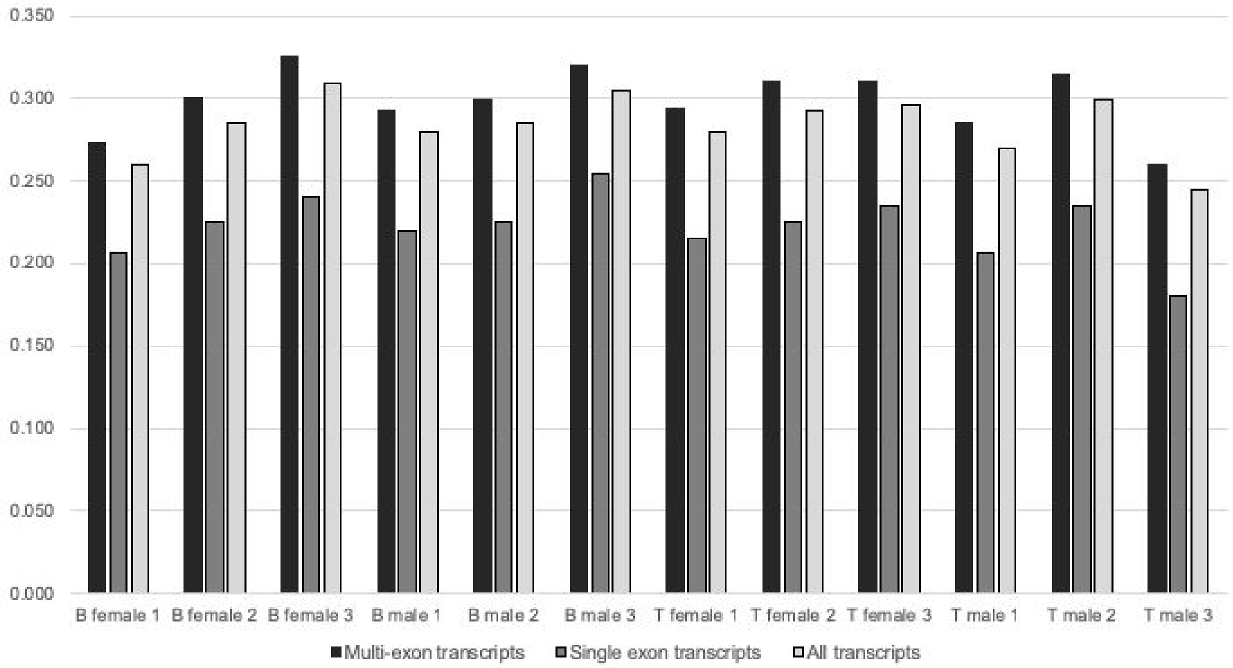


**Fig. S6. The sensitivity of transcript assembly by StringTie.** Sensitivity is defined as the proportion of reference transcriptome that exactly matches the assembled transcriptome. Values for sensitivity with respect to multi-exon transcripts, single exon transcripts, and all transcripts are shown. StringTie produces a mean of 194,316 assembled transcripts across all samples, with a mean of 34,600 such transcripts matching the total number of transcripts in the reference annotation (a mean sensitivity of 28.3%). Relative to the 53,961 protein-coding transcripts present in the reference, a mean of 15,452 assembled transcripts similarly showed matching intron chains (a mean sensitivity of 28.6%). These values represent sensitivity relative to the whole transcriptome and not the subset of expressed genes in B and T cells; by definition, these values are an underestimate of the actual sensitivity.

**Supplementary Figure 7**

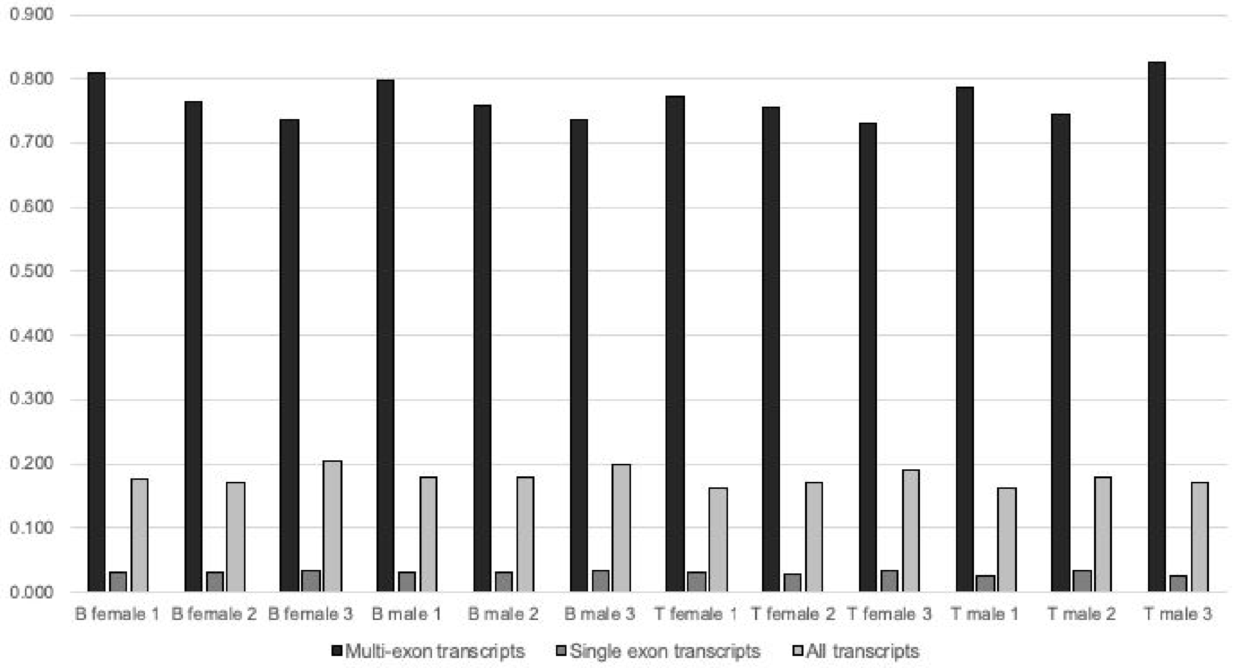


**Fig. S7. The specificity of transcript assembly by StringTie.** Specificity is defined as proportion of assembled transcripts that exactly match the reference transcriptome. StringTie performs better when assembling multi-exon transcripts compared to single exon transcripts: compared to a mean of 23.4% for multi-exon transcripts, a mean of 96.7% of all single exon transcripts predicted by StringTie are not present in the annotated reference, so can be considered novel if not due to transcriptional noise or misalignment. Considering the magnitude of multiple concordant and other non-concordant alignment of the raw reads, it is possible that these predictions are a result of alignment inaccuracies rather than the performance of StringTie itself.

**Supplementary Figure 8**

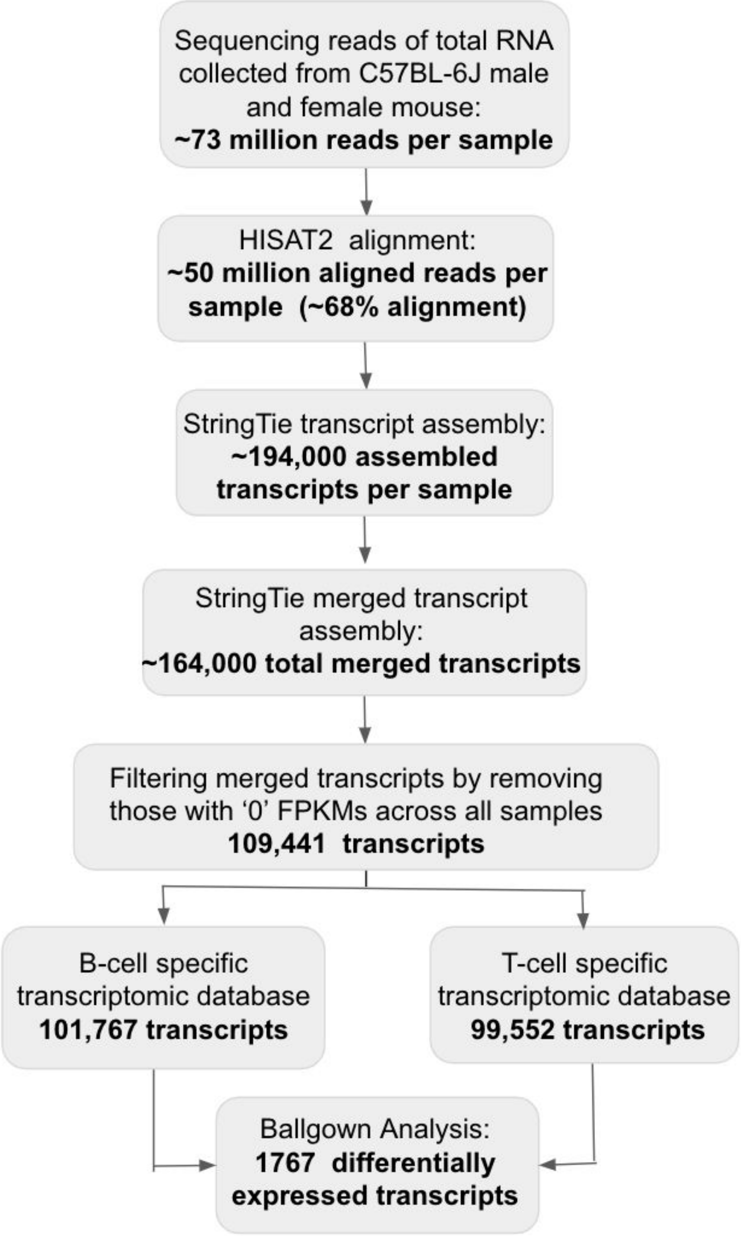


**Fig. S8. Differential expression analysis of transcripts.** The flowchart represents the different steps used to identify differentially expressed transcripts and a summary of the results obtained. Differential Expression analysis with Ballgown identified 1767 transcripts to be differentially expressed between B and T cells.

**Supplementary Figure 9**

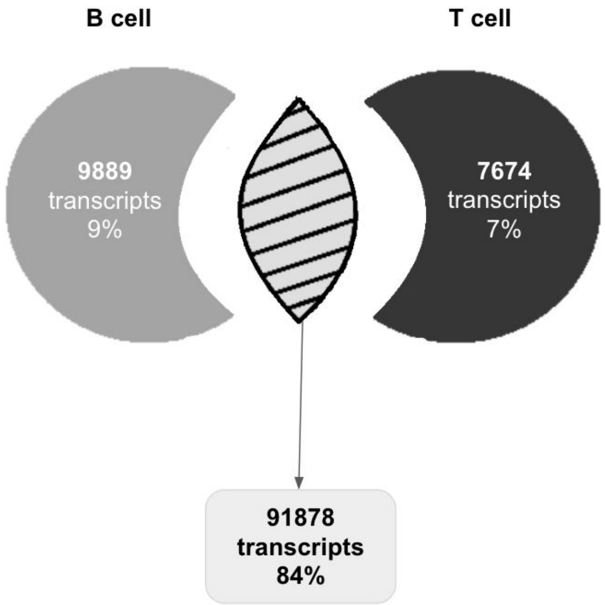


**Fig. S9. Venn diagram of transcripts identified.** The 109,441 transcripts identified in our analysis were categorised into B or T cell transcripts based on their expression levels. If the transcript expression levels corresponding to at least one of the three samples in a subgroup (B-male, B-female, T-male or T-female) was non zero, then the transcript was considered to be expressed in that particular cell. With this rule identified, 9,889 transcripts (9%) unique to B cells, 7,674 transcripts (7%) unique to T cells and 91,878 transcripts (84%) common to both B and T cells.

**Supplementary Figure 10**

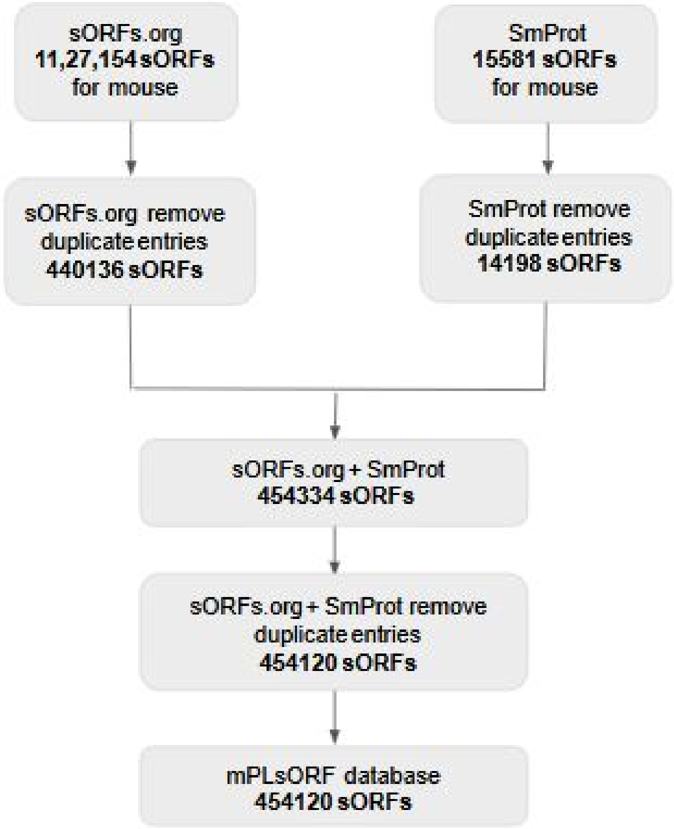


**Fig. S10. Creation of a sORF database.** Mouse sORFs for this work were obtained from two sources: sORFs.org containing 1,127,154 sORFs and SmProt containing 15,581 sORFs. Every entry in each of these datasets were individually filtered to remove duplicates resulting in 440,136 sORFs in sORFs.org and 14,198 sORFs in SmProt resulting in 454,120 sORF entries. Each of these sORFs were then assigned a unique identifier and relevant information about the sORFs including their genomic coordinates, strand information, source database, amino acid sequence and genomic annotation in our mPLsORF database.

**Supplementary Figure 11**

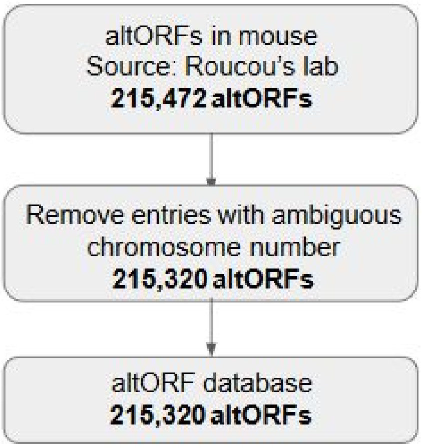


**Fig. S11. Creation of altORF database.** Information for the 215,472 mouse altORFs was downloaded from Xavier Roucou's lab, which were processed to remove entries with more than one designated chromosome. Strand information was ascertained separately and added to the database. This analysis resulted in a total of 215,320 altORFs that were used for our analysis .

### Supplementary Figure 12

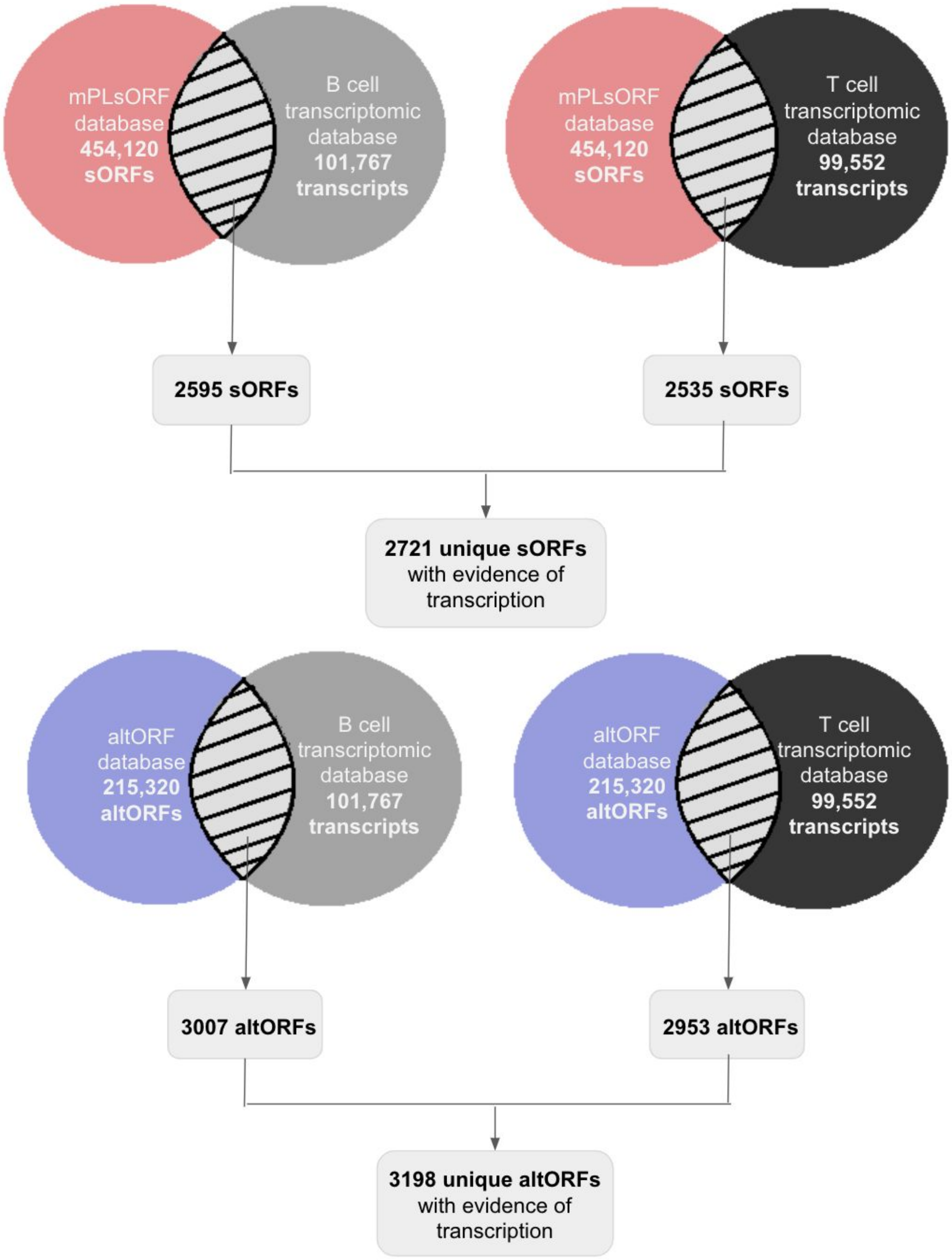


**Fig. S12. Evidence for transcription of altORFs and sORFs in B and T cells. (Top pane)** Genomic coordinates of 454,120 sORF entries were compared with our B and T cell specific transcriptomic dataset using getfasta bedtools intersect with parameters -f set to 0.99, -wo and -s, 2,595 and 2,535 sORFs were identified in B and T cells respectively. Comparing both these sets a total of 2,721 unique sORFs were identified with evidence of transcription in B and T cells (top panel). Similarly, 215,320 altORF entries were compared with our B and T cell specific transcriptomic fasta file and 3,007 and 2,953 altORFs were identified in B and T cells respectively. Comparing both sets a total of 3198 altORFs were identified with evidence of transcription in B and T cells (bottom panel).

**Supplementary Figure 13**

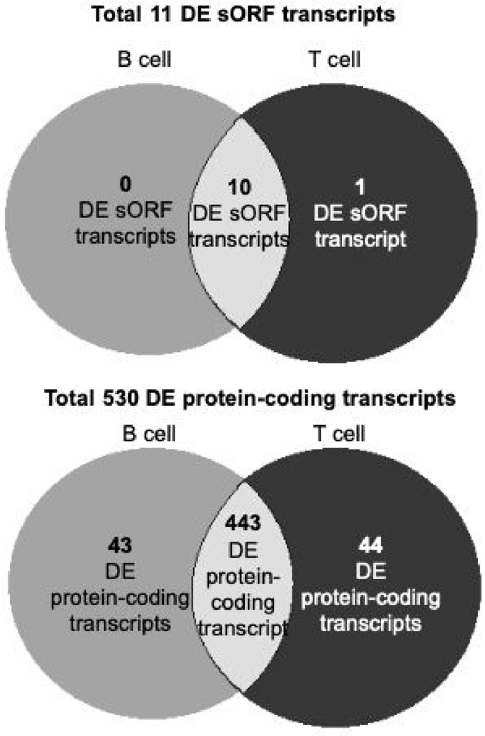


**Fig. S13. Classification of differentially expressed transcripts into B and T cells.** We identified 1767 DE transcripts. Of these, 11 were DE sORF transcripts, 145 were DE altORF transcripts and 530 were DE protein-coding transcripts. We further classified each of these sets of DE transcripts into B and T cell based on the transcript expression levels. One DE sORF, 7 DE altORF, 44 DE protein-coding transcripts were found to be unique to T cell. Nine DE altORF and 43 DE protein-coding transcripts were found to be unique to B cell. Ten DE sORFs, 129 DE altORFs and 443 DE protein-coding transcripts were found common to both B and T cells. Similarly, of the 1,767 DE transcripts identified in B and T cells, 204 are unique to B, 185 unique to T and 1,378 common to both B and T cells.

### Supplementary Figure 14

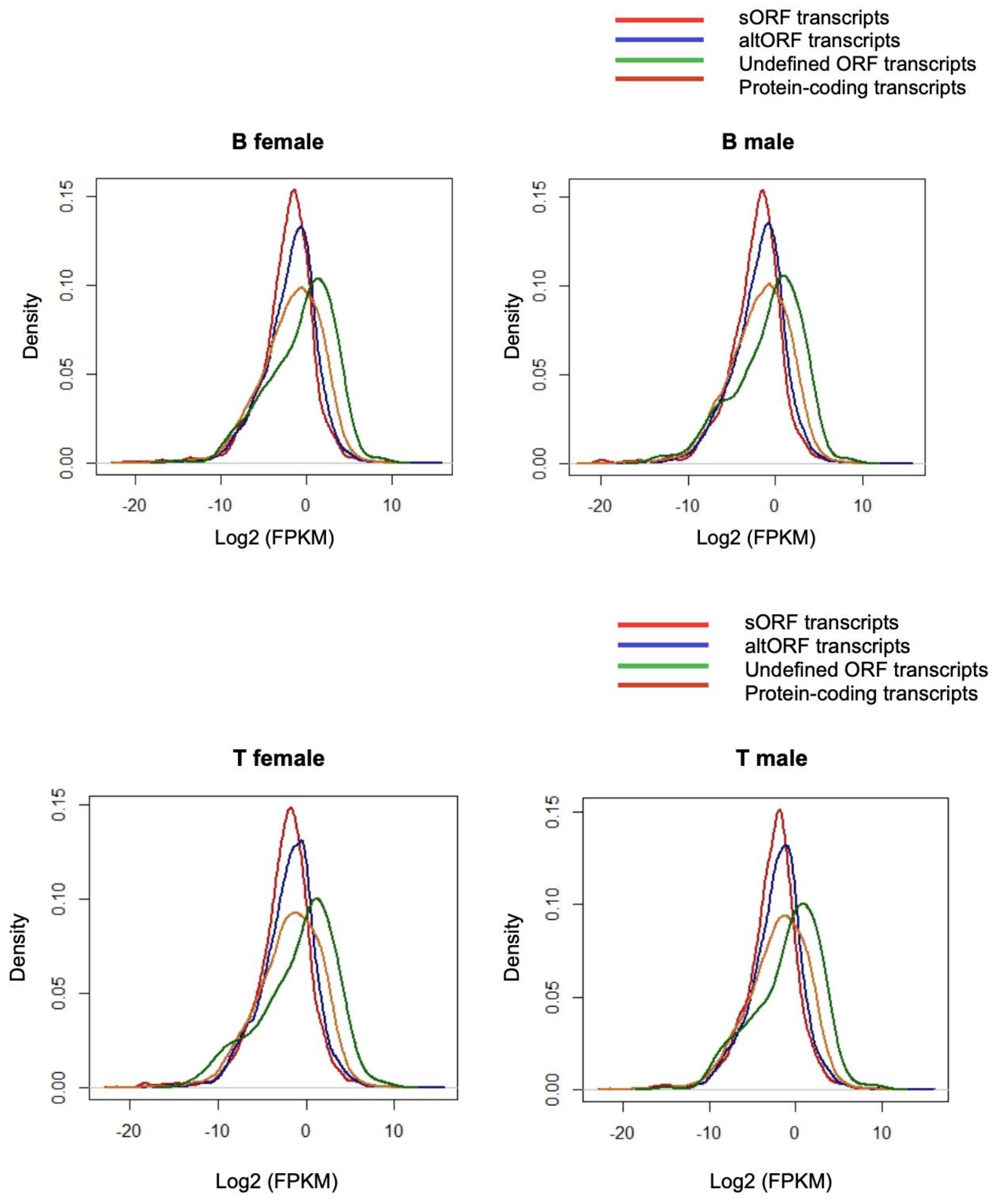


**Fig. S14. Transcript abundance distribution plots for different cell groups.** Transcript abundance plots calculated as $log_2$ of FPKM values (x-axis) is plotted against density (y-axis) to represent the distribution of transcript abundances of sORF (red), altORF (blue), undefined ORFs (green) and protein-coding transcripts (dark orange) for B male, B female, T male and T female.

**Supplementary Figure 15**

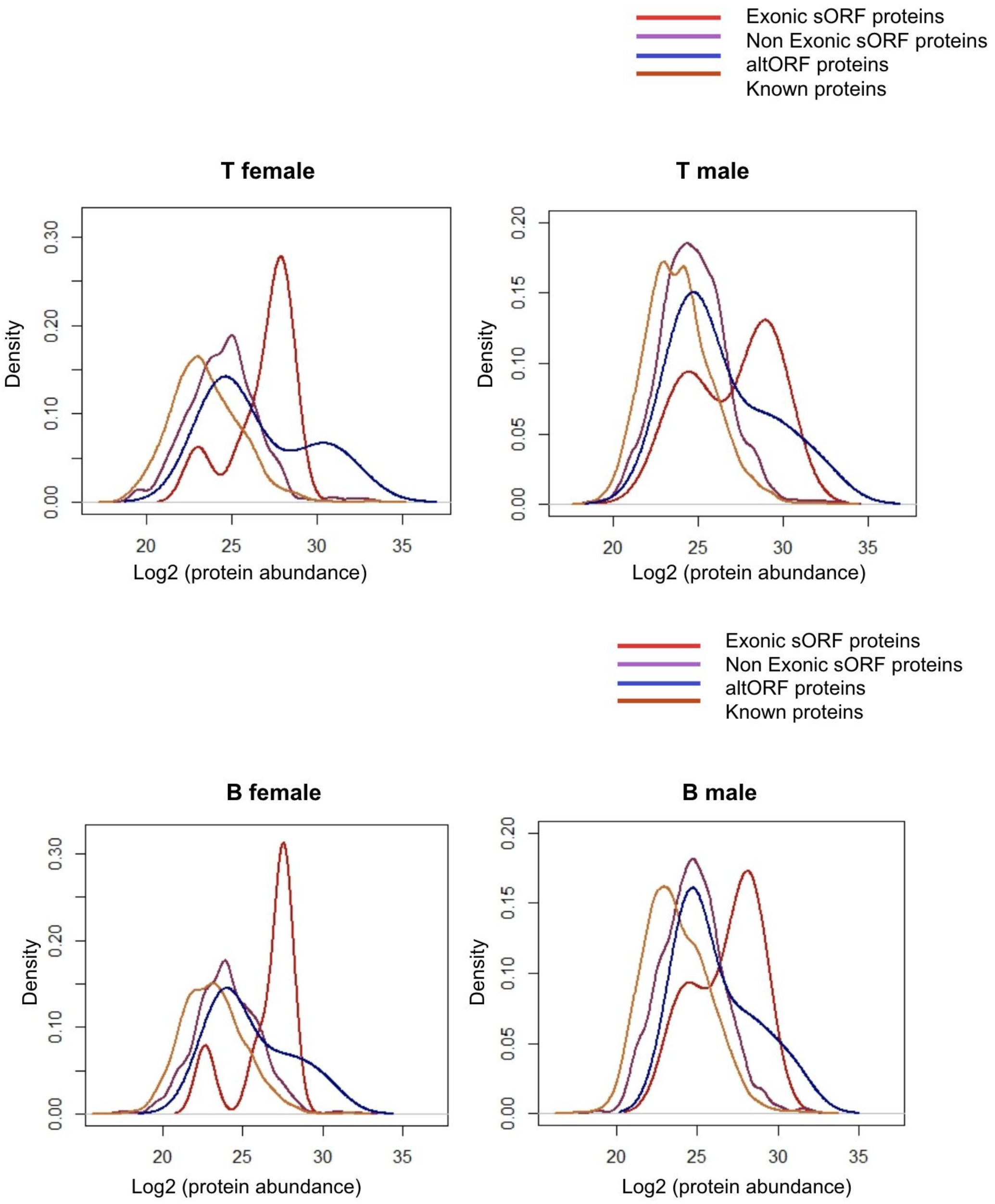


**Fig. S15. Protein abundance distribution plots for different cell groups.** Protein abundance plots calculated as $\log_2$ of protein abundance (x-axis) is plotted against density (y-axis) to represent the distribution of protein abundances of exonic sORF (red), non exonic sORF (violet), altORF (blue) and known proteins (dark orange) for B male, B female, T male and T female.

### Supplementary Figure 16

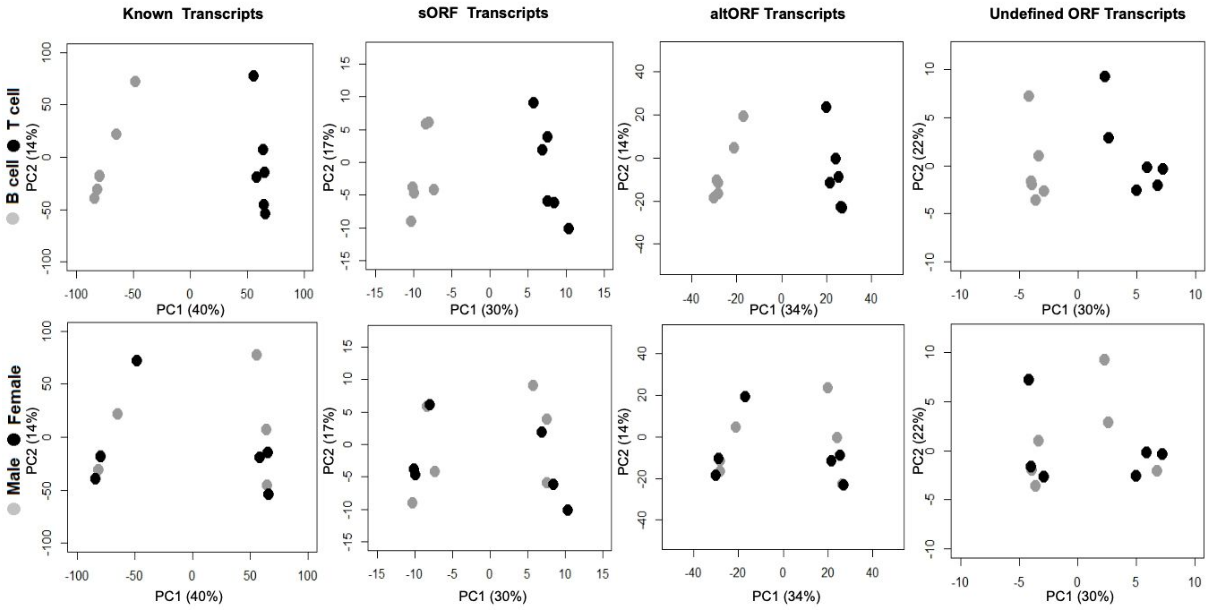


**Fig. S16. PCA of known protein coding, sORF, altORF and undefined novel transcripts.** PCA of transcript expression levels of known, sORF, altORF and undefined novel transcripts were performed. Only the first two principal components have been plotted and the amount of variance explained by each is mentioned in brackets. Top panel highlights that the transcript abundances of not only known transcripts but sORF, altORF, and undefined novel transcripts can distinguish B cells (grey) and T cells (black). Bottom panel indicates that the transcript abundances are unable to distinguish between male (grey) and female (black) of a particular cell type.

**Supplementary Figure 17**

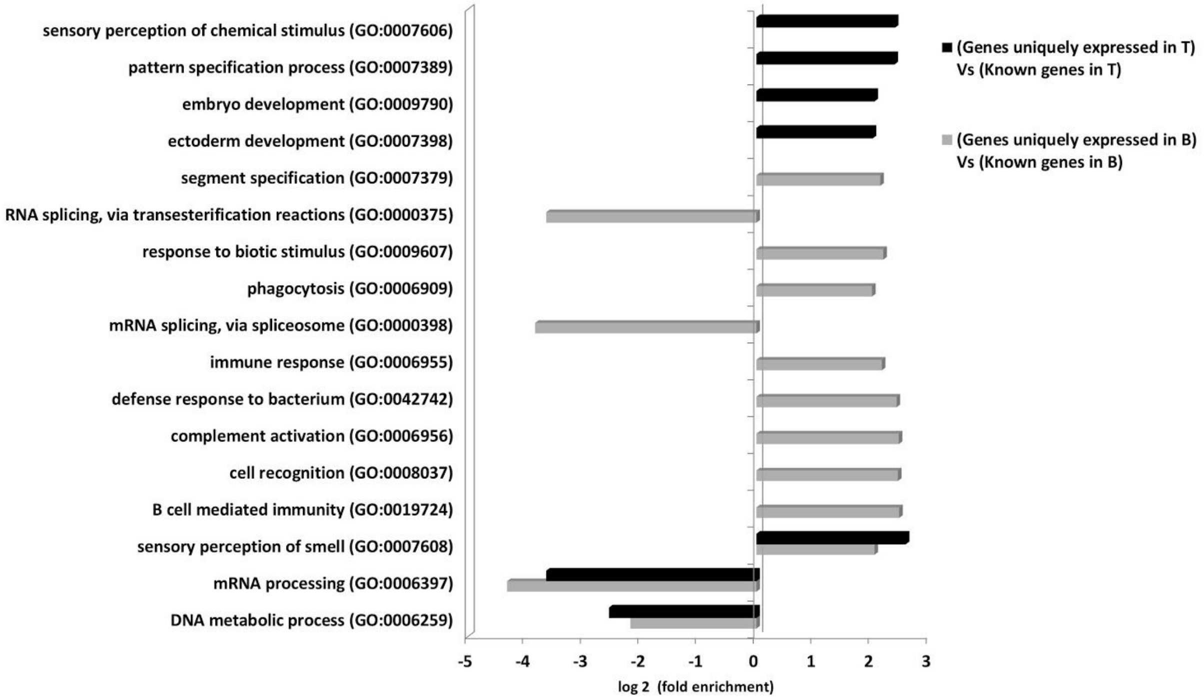


**Fig. S17. Functional classification of the genes expressed uniquely in B and T cells.** GO enrichment analysis was performed on the genes categorised to be expressed uniquely in either B or T cells. The figure includes the GO Slim biological process categories that are significantly (FDR correction < 0.005) enriched or depleted more than two fold, in B cells (grey bars) and in T cells (black bars), relative to the expected number based on all the known genes expressed in that respective cell type. GO analysis indicates that the genes expressed uniquely in both B and T cells are enriched with immune related functions. The 'B-cell mediated immunity' and 'immune response' category is two fold enriched only for the genes expressed uniquely in B cells.

**Supplementary Figure 18**

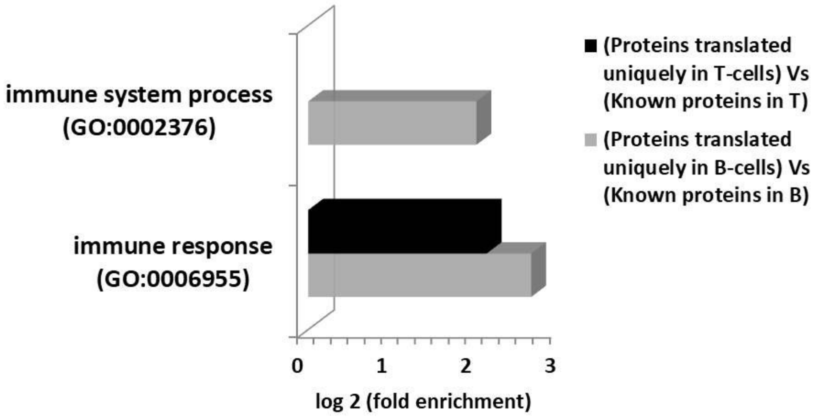


**Fig. S18. Functional classification of the proteins expressed uniquely in B and T cells.** GO enrichment analysis was performed on proteins categorised to be translated uniquely in a cell type. The figure includes the GO Slim biological process categories that are significantly (FDR correction < 0.1) enriched or depleted more than two-fold, in B cells (grey bars) and in T cells (black bars), relative to the expected number based on all the known proteins translated in that respective cell type. Only the immune related GO terms are enriched more than two fold for the B and T cell uniquely expressed proteins.

**Supplementary Figure 19**

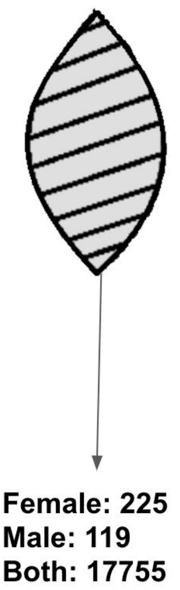


**Fig. S19. Genes expressed in both B and T cells.** Of the total known genes which are expressed in both B and T cells; 225 genes have expression uniquely in female, 119 in male and 17755 genes are expressed in both male and female mouse samples.

### Supplementary Figure 20

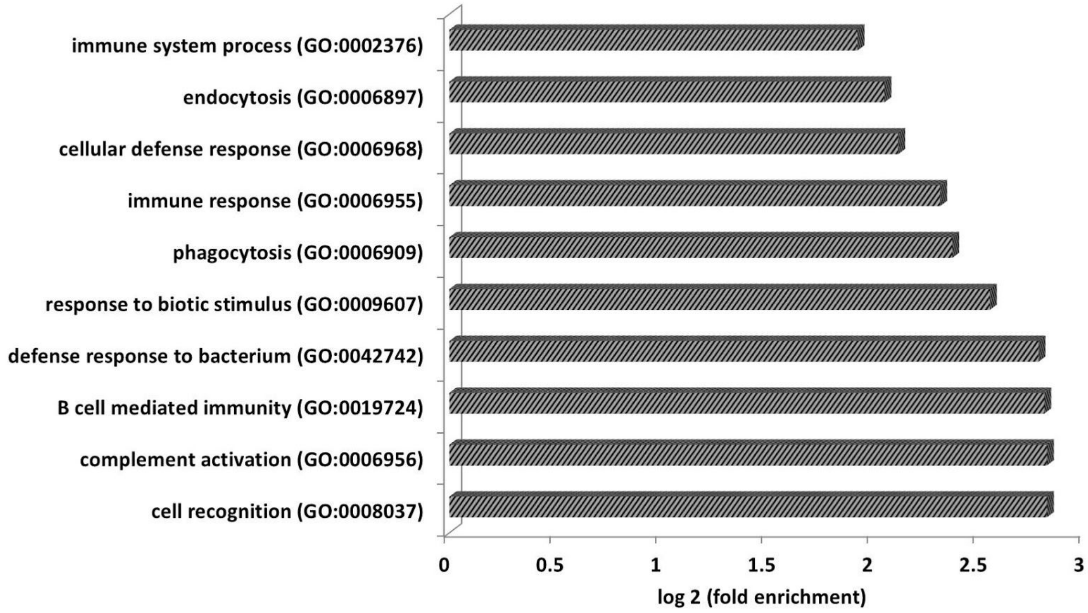


**Fig. S20. Functional classification of the differentially expressed genes in B and T cells.** GO enrichment analysis was performed on the genes that are differentially expressed in B and T cells. The figure depicts the GO Slim biological process categories that are significantly (FDR correction < 0.005) enriched or depleted more than two fold, relative to the expected number based on all the known genes expressed in both the cells. GO analysis indicates that the genes which are differentially expressed between B and T cells are involved in immune response.

### Supplementary Figure 21

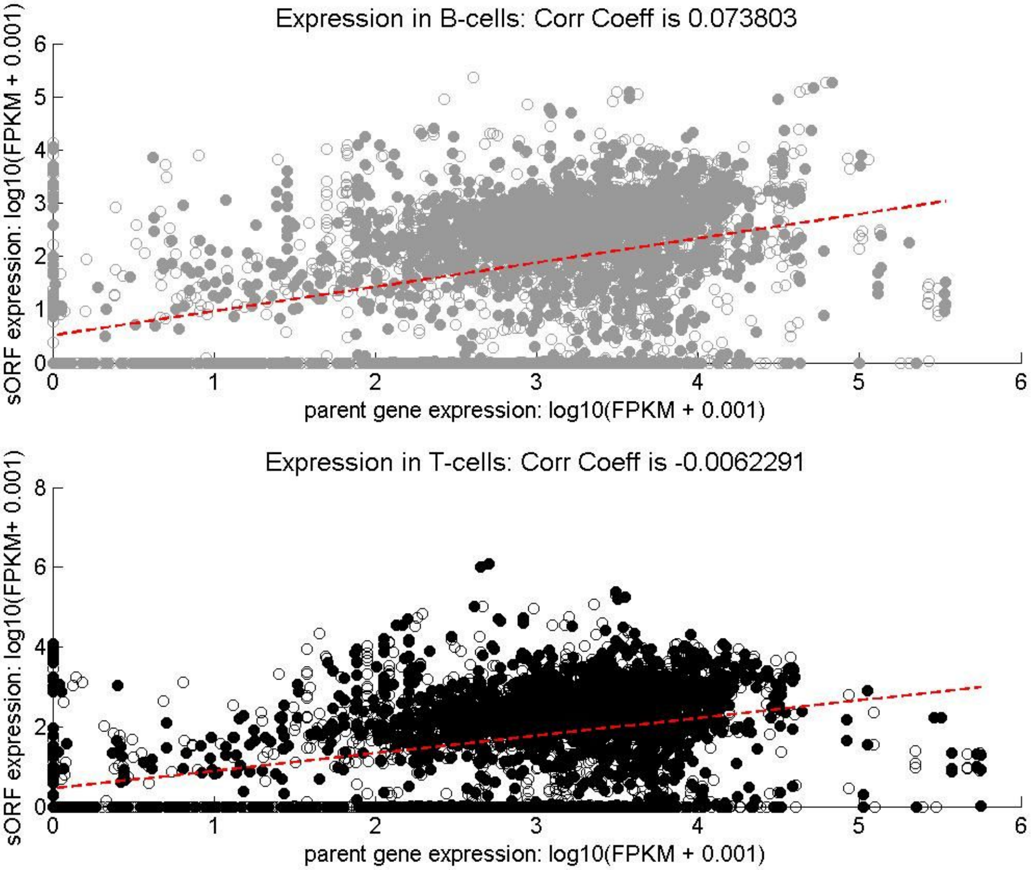


**Fig. S21. Correlation plot of sORF vs its respective overlapping gene abundance in (a) B cells (top panel) and in (b) T cells (bottom panel).** An overlapping gene within which a sORF transcript lies was extracted from the ensembl's GRCm38 annotation file using bedtools intersect. Each point in these plots depict the abundance of a sORF (y-axis) plotted against the abundance of its overlapping gene (x-axis). The log scaled abundances are rescaled to fit in the first quadrant. Filled circles represent the abundances from female mice whereas the unfilled circles represent abundances from male samples. From the correlation analysis we observe that in both B and T cells expression of a sORF is uncorrelated with the expression of its overlapping gene (Correlation coefficients are almost 0).

**Supplementary Figure 22**

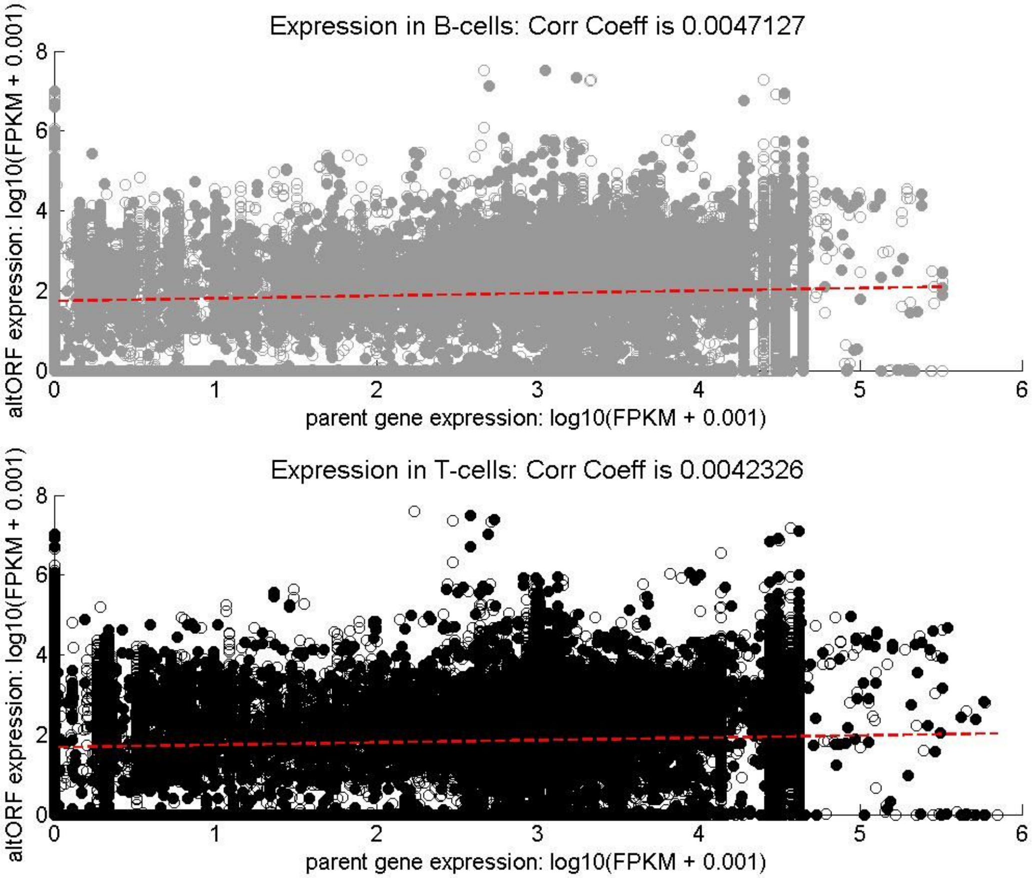


**Fig. S22. Correlation plot of an altORF vs abundance of an overlapping it (a) B cells (top panel) and in (b) T cells (bottom panel).** Information of the nearby gene name is obtained from Dr. Roucou's lab. Each point in these plots depict the abundance of an altORF (y-axis) plotted against the abundance of the overlapping gene (x-axis). The log scaled abundances are rescaled to fit in the first quadrant. Filled circles represent the abundances from female mice whereas the unfilled circles represent abundances from male samples. From the correlation analysis we observe that in both B and T cells expression of altORFs is independent to the expression of its nearby or overlapping (Correlation coefficients are almost 0).

### Supplementary Figure 23

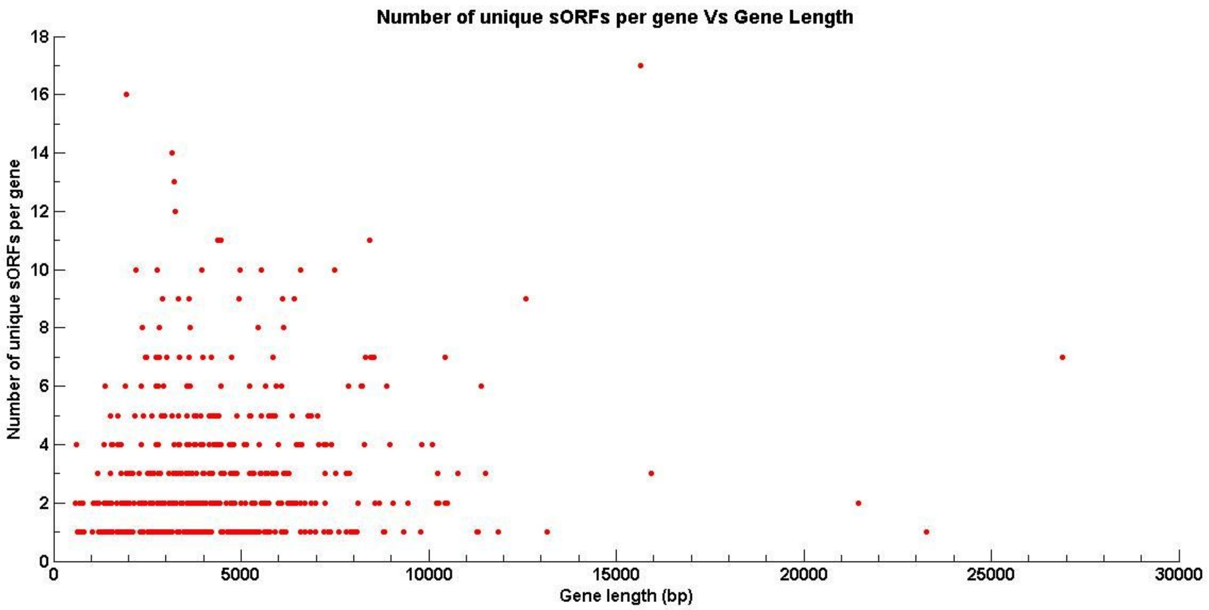


**Fig. S23. The correlation of the number of unique sORFs to the length of an overlapping known gene**. Each point in this plot depicts the number of unique sORFs (y-axis) overlapping with a known gene vs that gene's length (x-axis). The length of the longest transcript of a known gene, determined by ensembl's biomart and by GTFtools-0.6.5, is considered to be that gene's length.

**Supplementary Figure 24**

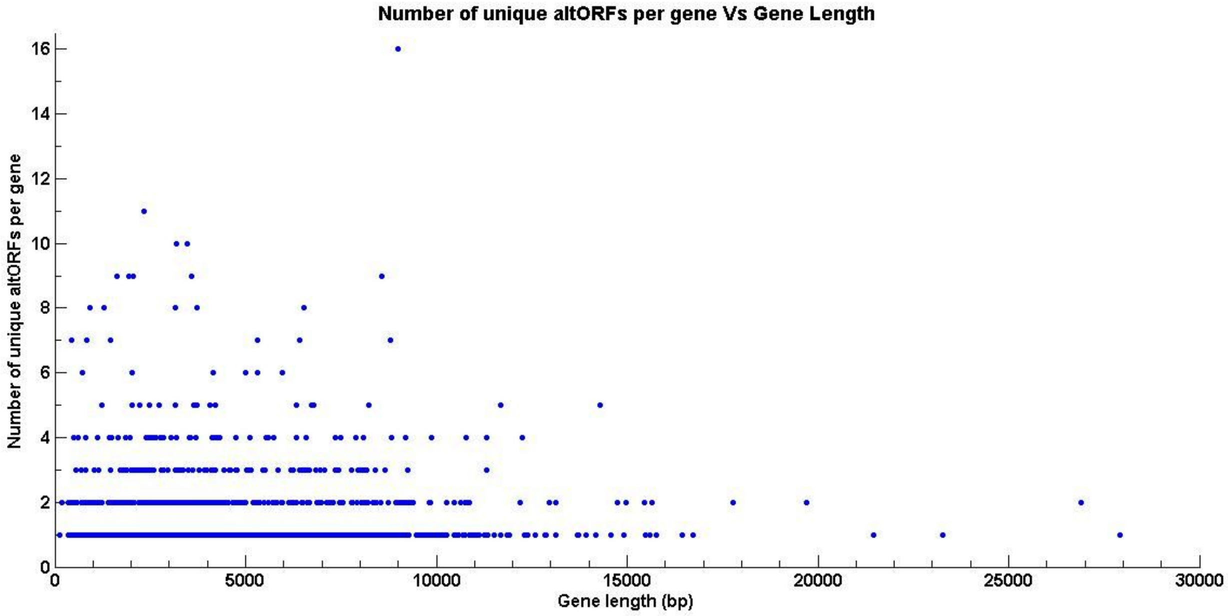


**Fig. S24. Correlation plot of number of unique altORFs to the length of an overlapping known gene.** Each point in this plot depicts the number of unique altORFs (y-axis) mapped to a known gene overlapping with it vs that gene's length (x-axis). The length of the longest transcript of a gene, determined by ensembl's biomart and by GTFtools-0.6.5, is considered to be that gene's length.

### Supplementary Figure 25

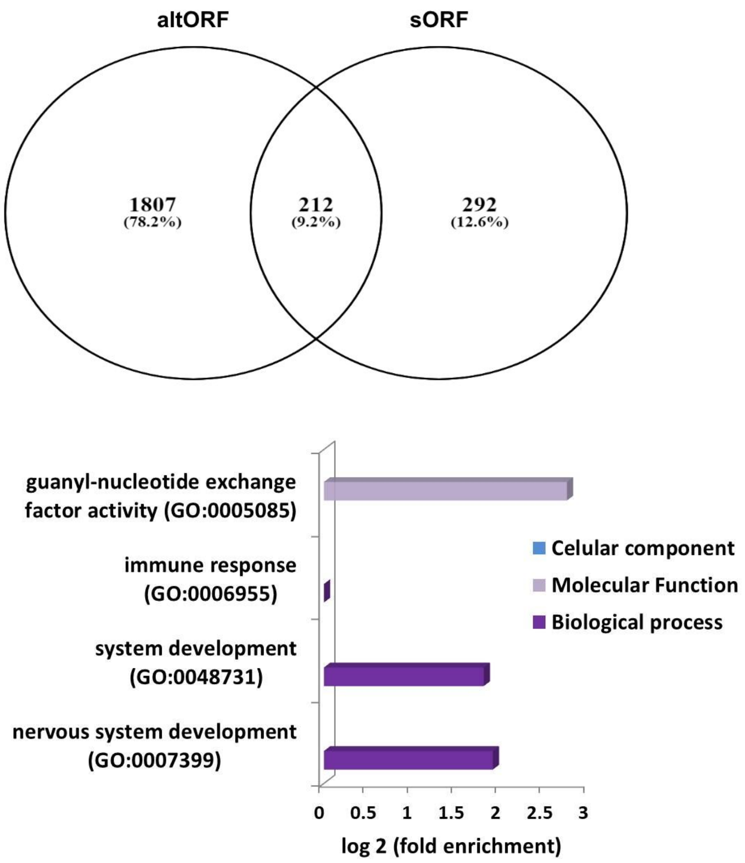


**Fig. S25. Functional annotation analysis of the known genes that overlap with both sORFs and altORFs. (Top panel)** 212 known genes overlap with both sORFs and altORFs, while 1807 and 292 genes overlap with only altORFs and sORFs respectively. **(Bottom panel)** GO enrichment analysis was performed on the 212 genes overlapping with both sORFs and altORFs. The figure includes the GO Slim biological process and molecular function categories that are significantly (FDR correction < 0.1) enriched or depleted, relative to the expected number based on all the known genes expressed in both the cells. None of the genes which overlap with both sORFs and altORF are involved in immune related biological processes (observed number of genes for immune response category is 0).

**Supplementary Figure 26**

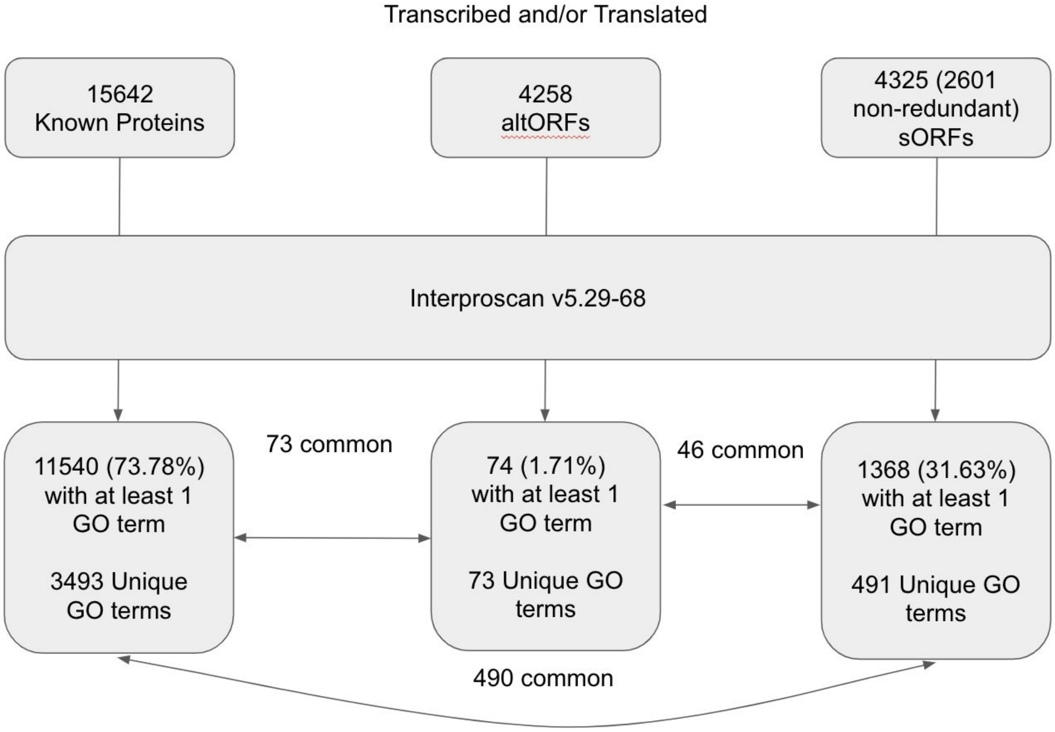


**Fig. S26. Workflow of GO annotation of altORFs and sORFs in preparation for GO analysis.** A list of amino acid sequences for all known proteins, altORFs, and sORFs with either transcription or translation evidence in at least 1 of the 12 samples was compiled and analyzed using Interproscan v5.29-68. GO annotation for the known proteins were also generated this way to ensure equal comparison between altORFs and sORFs which are unlikely to have GO annotations from other sources such as experimental methods. The list of 3493 GO terms from these known proteins was then used as the background list for subsequent GO enrichment analysis of sORFs. 490 GO terms present in known proteins were shared with sORFs, and all 73 terms found in altORFs were also present in the known proteins. However, only 46 terms were shared between sORFs and altORFs.

**Supplementary Figure 27**

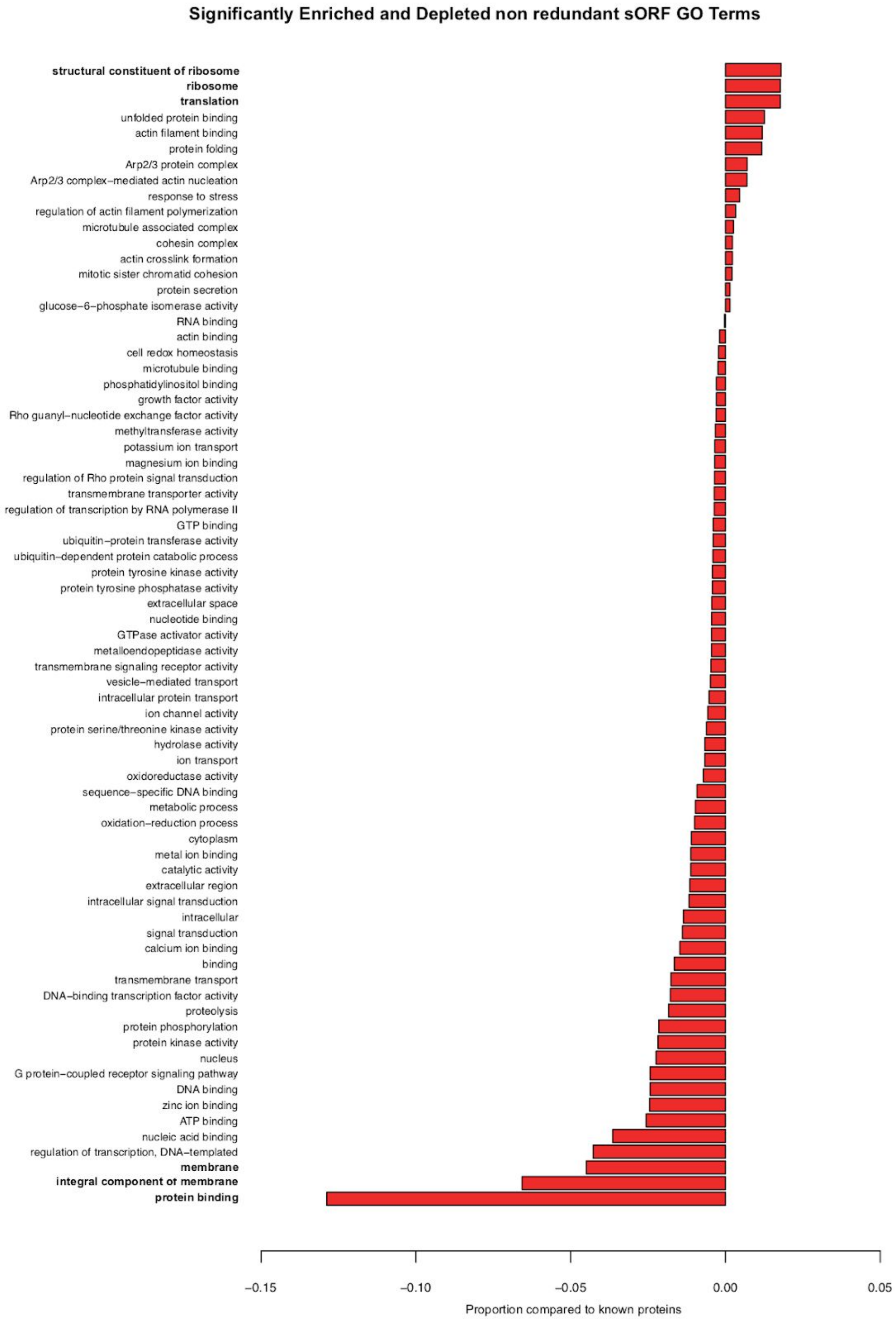

**Fig. 27. List of significantly enriched or depleted GO terms in sORFs after removal of redundant sORFs as compared to GO terms from known proteins.** Redundant sORFs were identified as sORFs with genomic coordinates that were exactly the same or were encompassed by another sORFs genomic coordinates. Counts of GO terms in known proteins and the non-redundant sORFs were tabulated and normalised against the number of known proteins and non redundant sORFs analyzed respectively to obtain proportions. The bioconductor package q-value and R base chisq.test with simulated p-values was then used to calculate GO terms that were significantly different between the 2 groups based on a p-value of $< 0.01$ and q-value of $< 0.01$. The proportions shown in the graph are obtained from subtracting the known protein GO term proportion from the non-redundant sORF GO term proportion. The top three enriched and depleted categories are highlighted in bold.

**Supplementary Figure 28**

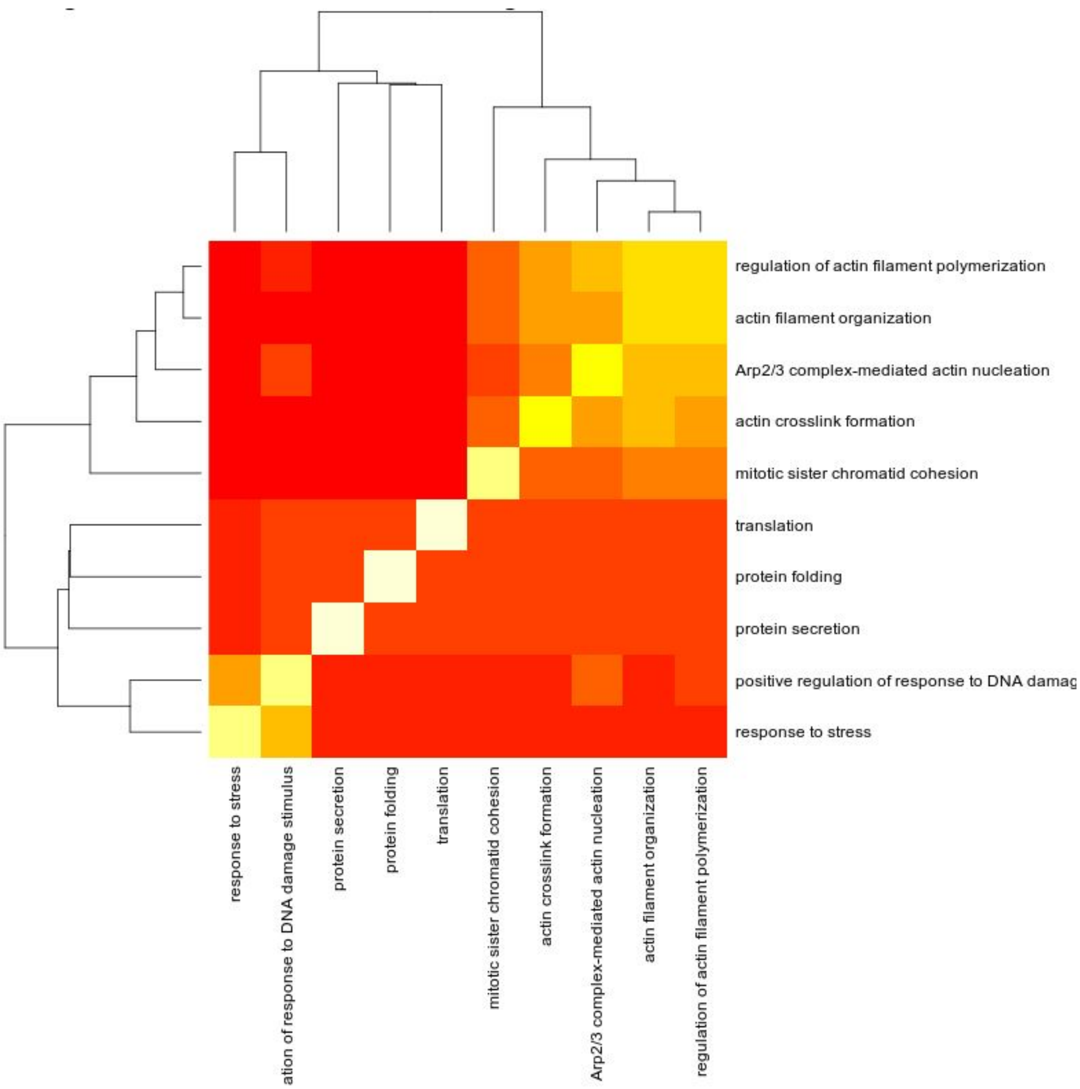


**Fig. S28. Clustering of significantly enriched GO Terms in non redundant sORFs.** Significantly enriched GO terms in non redundant sORFs were identified using a p-value < 0.01, q-value < 0.01, and proportion more than known protein GO Term proportion. Distances between GO terms were calculated using getTermSim function from bioconductor GOSim package using default settings. The distance metric was then used to cluster the terms to enable easier interpretation by grouping similar GO terms.

### Supplementary Figure 29

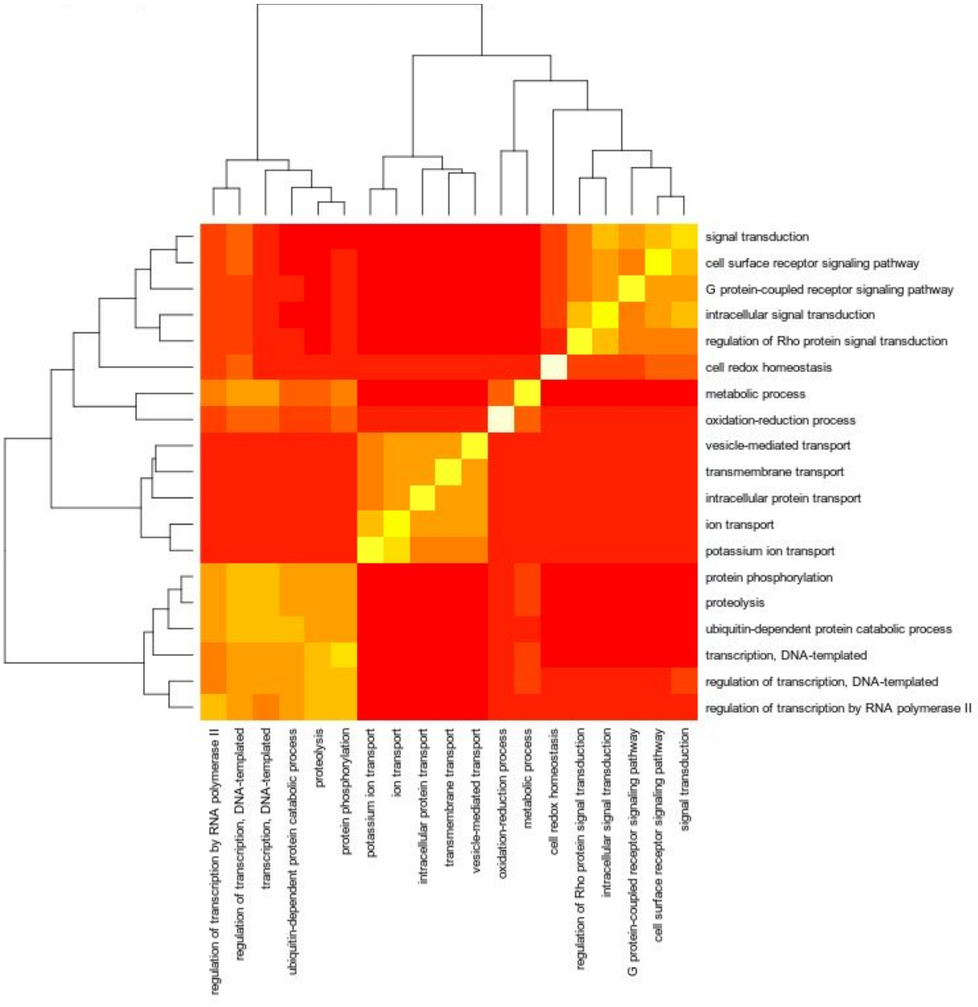


**Fig. S29. Clustering of significantly depleted GO Terms in non-redundant sORFs.** Significantly depleted GO terms in non-redundant sORFs were identified using a p-value < 0.01, q-value < 0.01, and proportion less than known protein GO Term proportion. Distances between GO terms were calculated using getTermSim function from bioconductor GOSim package using default settings. The distance metric was then used to cluster the terms to enable easier interpretation by grouping similar GO terms.

**Supplementary Figure 30**

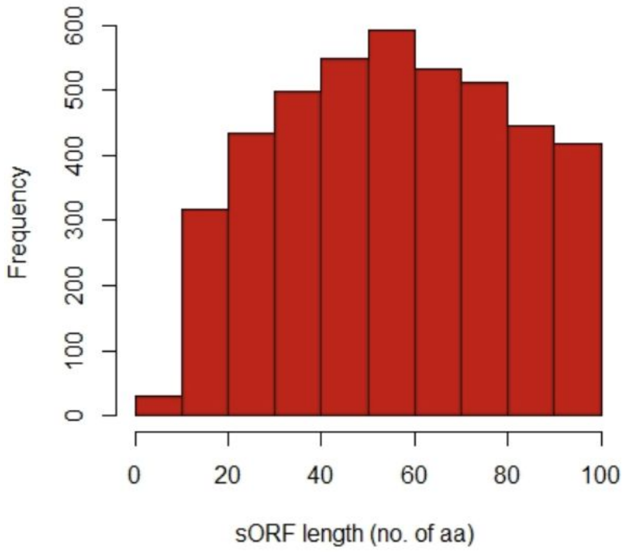


**Fig. S30. Histogram of sORF lengths for sORFs identified in B and T cells.** sORF lengths (x-axis) defined as the number of amino acid in the translated products of the sORF plotted against the frequency of a particular sORF length (y-axis), for sORFs identified in mouse B and T cells. sORF lengths vary from 2-100 amino acids with maximum containing 50-60 amino acids.

**Supplementary Figure 31**

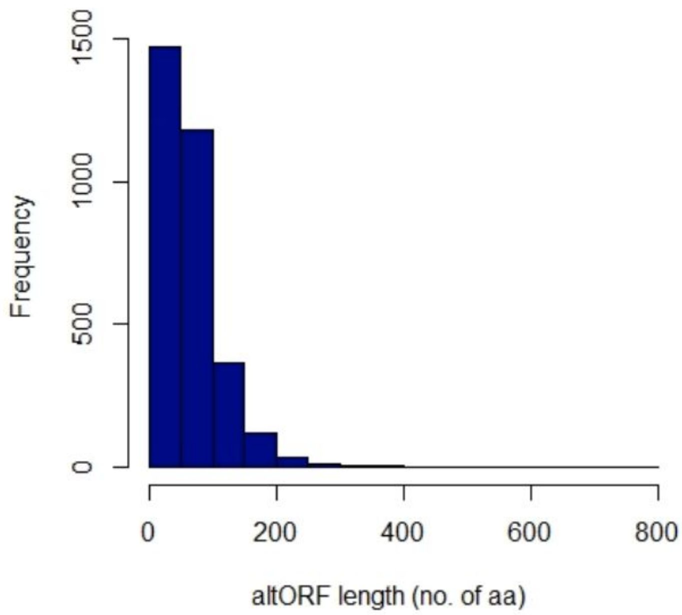


**Fig. S31. Histogram of altORFs lengths for altORFs identified in B and T cells.** altORF lengths (x-axis) defined as the number of amino acids (aa) in the translated products of the altORF plotted against the frequency of a particular altORF length (y-axis), for altorfs identified in mose B and T cells. altORF lengths vary from 29-1538 aa with maximum in the 0-50 bin and specifically in the 29-50 aa range.

### Supplementary Figure 32

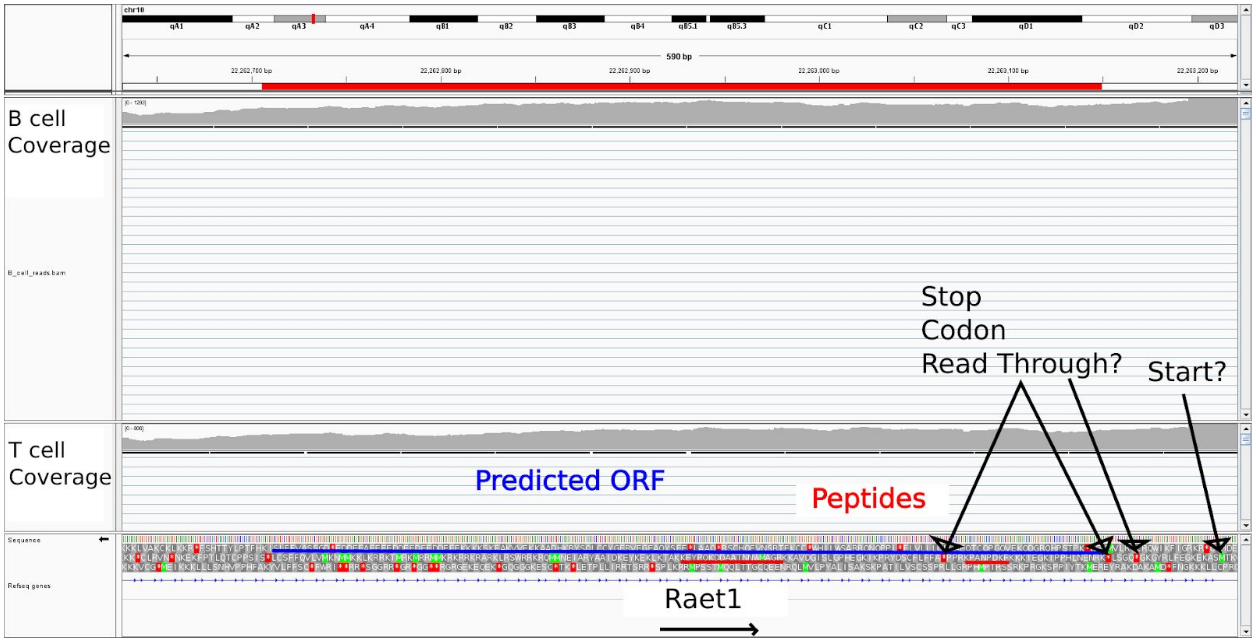


**Fig. S32. Undefined novel ORF antisense to *Raet1* pseudogene**. Predicted undefined novel ORF (blue line) was identified by proteogenomics analysis. The novel ORF had two distinct peptides (red lines) mapped to it. The novel transcript has two stop and two start codons.

### Supplementary Figure 33

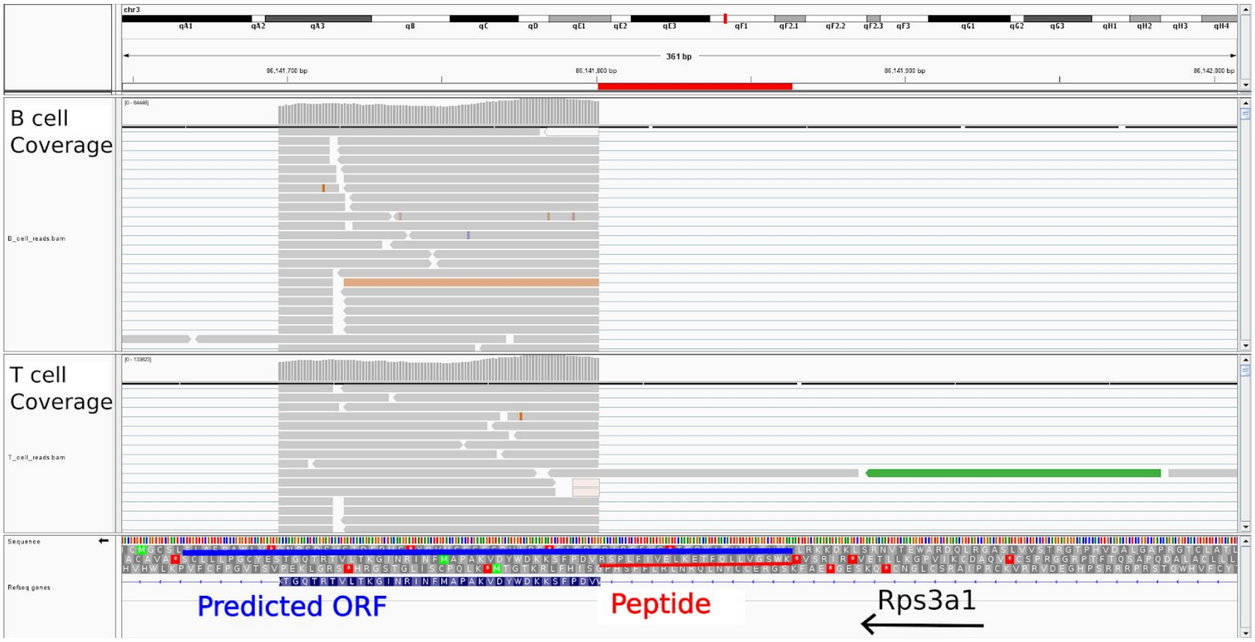


**Fig. S33. Undefined novel ORF as an intron insertion or novel exon of *Rps3a1*.** An undefined novel ORF (blue line) spans the exon of *Rps3a1*, a known gene which is a constituent of the 40s component of the ribosome. The identified peptides (red line) map to an unannotated exon and is in the same frame, suggesting it may be incorporated into the exon as an insertion.

### Supplementary Figure 34

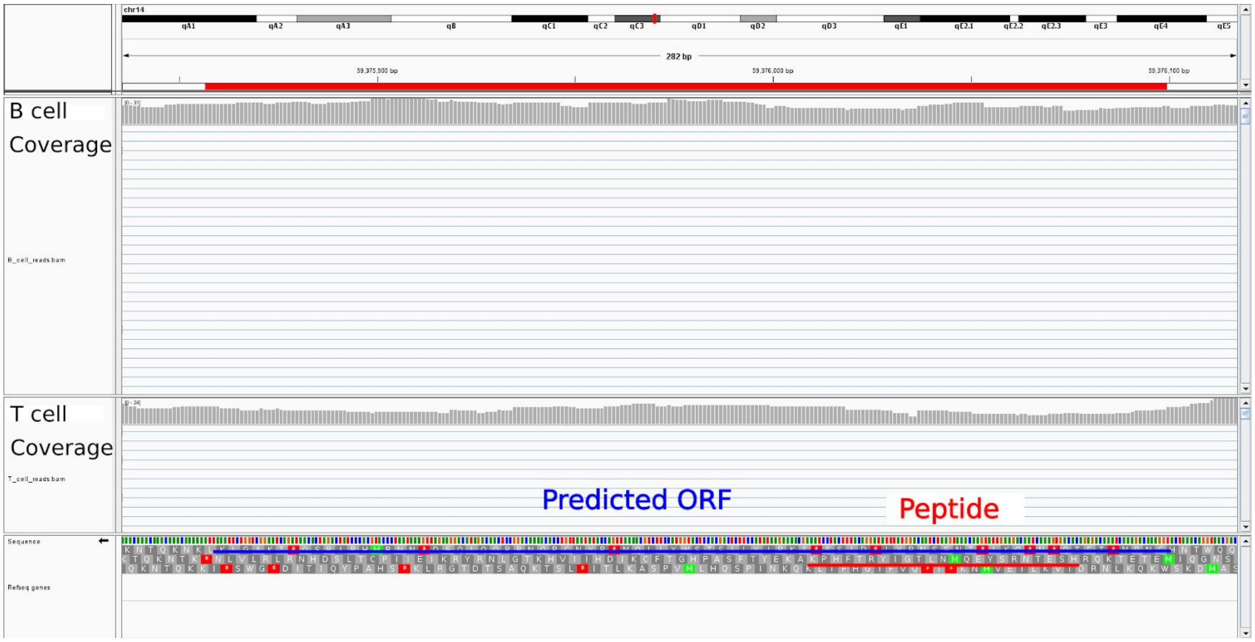


**Fig. S34. Undefined novel ORF in intergenic region on Chr 14.** An undefined novel ORF (blue line) in intergenic regions of Chr 14 was identified by proteogenomic analysis and by extension of the aligned peptide fragments (red line) both up and downstream until a stop codon or a start codon was encountered.

### Supplementary Figure 35

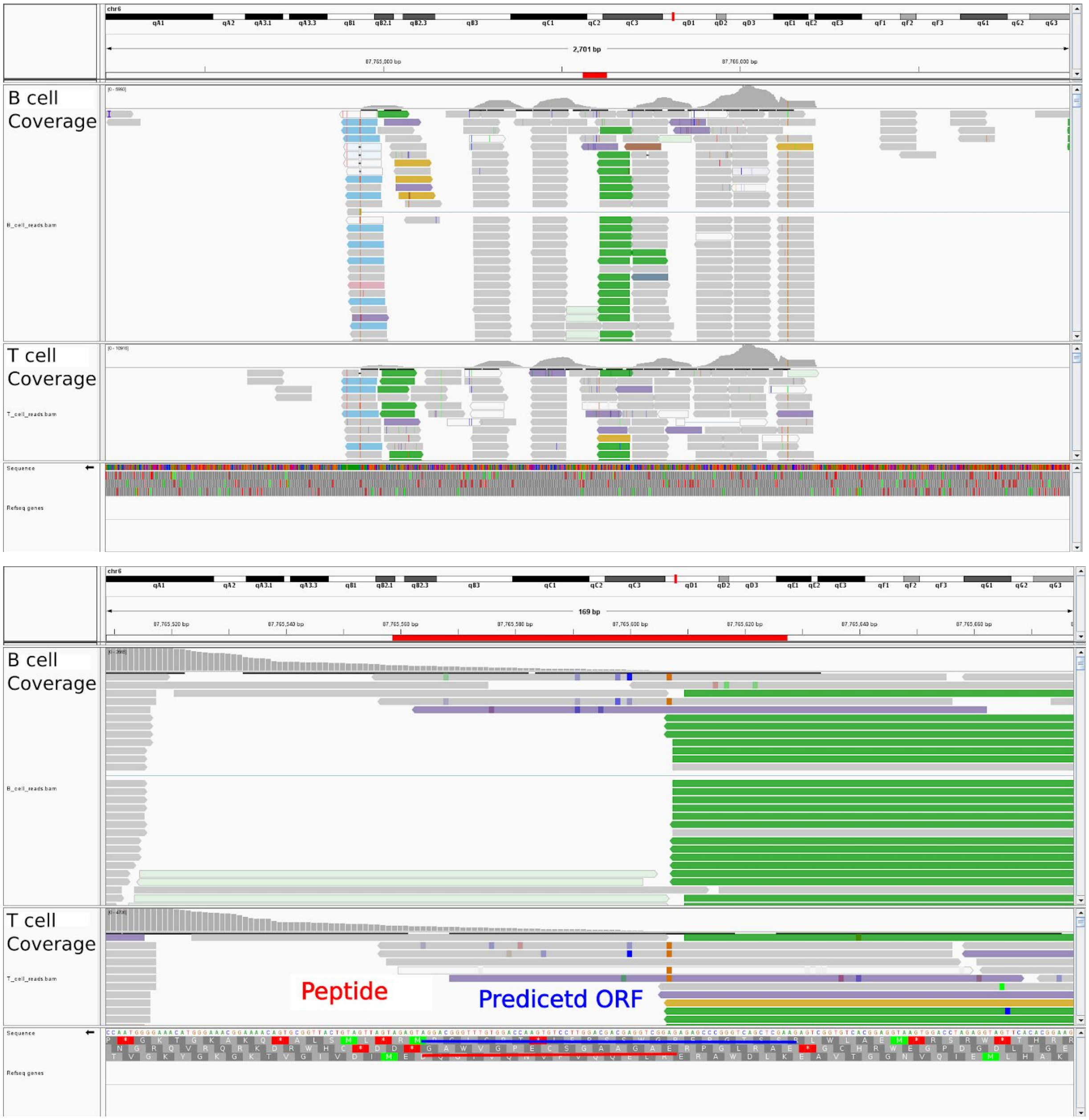


**Fig. S35. An undefined novel ORF is a processed pseudogene ENSMUSG00000068262.** The predicted ORF (blue line) was generated by extension of peptide aligned fragments both up and downstream until a stop codon or a start codon was encountered. **Top panel** provides a zoomed out view up of aligned peptides showing where it lies within a relatively low transcribed region of the pseudogene. **Bottom panel** provides a zoomed in view of the pseudogene.

**Supplementary Table 1**

| Sample ID | GEO Sample ID | Sample identify |
|---|---|---|
| 10966_8_1 | GSM2480724 | B cell female |
| 10966_8_2 | GSM2480725 | B cell female |
| 10966_8_3 | GSM2480726 | B cell female |
| 11048_5_13 | GSM2480727 | B cell male |
| 11048_5_14 | GSM2480728 | B cell male |
| 11048_5_15 | GSM2480729 | B cell male |
| 11049_4_4 | GSM2480730 | T cell female |
| 11049_4_5 | GSM2480731 | T cell female |
| 11049_4_6 | GSM2480732 | T cell female |
| 11049_5_16 | GSM2480733 | T cell male |
| 11049_5_17 | GSM2480734 | T cell male |
| 11049_5_18 | GSM2480735 | T cell male |

**Table S1. Information and Sample id for mice used in transcriptomic analysis**

Sample IDs used in the transcriptomic analysis, their corresponding GEO sample id and sample information is provided in the table. GEO sample id can be accessed using GEO accession: GSM2480756.

**Supplementary Table 2**

| Gene mapped to Grcm38 | number of unique sORFs per gene | Gene Length |
|---|---|---|
| Lpp | 17 | 15647 |
| Eef1b2 | 16 | 1939 |
| Mgat1 | 14 | 3151 |
| Gpm6a | 13 | 3215 |
| Heatr3 | 12 | 3251 |
| Mgat5 | 11 | 8436 |
| Rack1 | 11 | 4456 |
| Ppp1r3b | 11 | 4386 |
| Zmiz1 | 10 | 7482 |
| Eef2k | 10 | 6580 |
| Fus | 10 | 5536 |
| Rabgap1 | 10 | 4967 |
| Acsl3 | 10 | 3950 |
| Olfr56 | 10 | 2767 |
| Akt1s1 | 10 | 2207 |

**Table S2. List of top few genes with the number of unique altORFs mapped to them**

Tabulated above are the list of genes, their length and the number of unique sORFs overlapping with them. A maximum of 17 unique sORFs are observed to overlap to the gene '*Lpp*' having length of 15647 bp.

**Supplementary Table 3**

| Roucou's lab gene | number of unique altORFs per gene | Gene-length |
|---|---|---|
| Gvin1 | 16 | 9005 |
| Duxbl3 | 11 | 2336 |
| 6530403H02Rik | 10 | 3191 |
| Pira2 | 10 | 3462 |
| Gm14308 | 9 | 1628 |
| Gm3488 | 9 | 1945 |
| Gm3558 | 9 | 2055 |
| Gm21685 | 9 | 3589 |
| Tcf4 | 9 | 8577 |
| Gm3893 | 8 | 909 |
| Il11ra2 | 8 | 1299 |
| Gm2808 | 8 | 3159 |
| Amy2a2 | 8 | 3717 |
| Fubp1 | 8 | 6523 |

**Table S3. List of top few genes with the number of unique altORFs mapped to them**

Tabulated below are the list of genes, their length and the number of unique altORFs overlapping with them. A maximum of 16 unique altORFs overlap to the gene 'Gvin1' having length of 9005 bp

### Supplementary Table 4

| Gene | Disease Phenotype | Mutation counts |
|---|---|---|
| ACTG1 | Baraitser-Winter_syndrome | 1 |
| ACTN4 | Glomerulosclerosis_focal_and_segmental | 1 |
| AK2 | Reticular_dysgenesia | 2 |
| ARCN1 | Craniofacial_syndrome | 2 |
| ATP2A2 | Schizophrenia | 1 |
| ATP2A2 | Darier_disease | 17 |
| ATP2A2 | Acrokeratosis_verruciformis | 1 |
| BCLAF1 | Colorectal_cancer | 1 |
| BUB3 | Variegated_aneuploidy | 1 |
| CALM1 | Catecholaminergic_polymorphic_ventricular_tachycardia | 2 |
| CALR | Schizoaffective_disorder | 1 |
| DDX3X | Intellectual_disability | 5 |
| DDX5 | Fibrosis_risk_association_with | 1 |
| DYNC1H1 | Malformations_of_cortical_development | 1 |
| FLNA | Thoracic_aortic_aneurysms_and_dissections | 1 |
| FLNA | Thoracic_aortic_aneurysms | 1 |
| FLNA | Otopalatodigital_syndrome_2 | 6 |
| FLNA | Otopalatodigital_syndrome_1 | 4 |
| FLNA | Mental_retardation_X-linked | 1 |
| FLNA | Melnick-Needles_syndrome_epilepsy_&_heterotopia_periventricular_nodular | 1 |
| FLNA | Lower_resp._tract_infection_bilateral_lung_emphysema_with_basal_atelectasis_bronchospasm_and_pulmonary_artery_hypertension | 1 |
| FLNA | Heterotopia_periventricular_with_skeletal_dysplasia | 1 |
| FLNA | Heterotopia_periventricular_nodular | 7 |
| FLNA | Heterotopia_periventricular | 12 |
| FLNA | Heterotopia_nodular | 1 |
| FLNA | Frontometaphyseal_dysplasia | 1 |
| FLNA | FG_syndrome | 1 |
| FLNC | Frontotemporal_dementia_behavioural_variant | 1 |
| FLNC | Cardiomyopathy_hypertrophic | 1 |
| GOT1 | Aspartate_aminotransferase_deficiency | 1 |
| GPI | Glucosephosphate_isomerase_deficiency | 2 |
| HIST3H3 | Intellectual_disability | 1 |
| HNRNPU | Lennox-Gastaut_syndrome | 1 |
| HNRNPU | Epileptic_encephalopathy | 1 |
| HSPA9 | Parkinson_disease | 1 |
| HSPA9 | EVEN-PLUS_syndrome | 1 |
| LDHA | Lactate_dehydrogenase_deficiency | 1 |

| | | |
|---|---|---|
| LDHB | Lactate_dehydrogenase_deficiency | 2 |
| LMNA | Ventricular_arrhythmia | 1 |
| LMNA | Spinal_muscular_atrophy_with_cardiac_involvement | 1 |
| LMNA | Peripheral_neuropathy | 1 |
| LMNA | Partial_lipodystrophy_atypical | 1 |
| LMNA | Muscular_dystrophy_limb_girdle_with_severe_heart_failure_and_lipodystrophy | 1 |
| LMNA | Muscular_dystrophy_limb_girdle | 9 |
| LMNA | Muscular_dystrophy_Emery-Dreifuss | 10 |
| LMNA | Muscular_dystrophy | 5 |
| LMNA | Cardiomyopathy_right_ventricular_&_Charcot-Marie-Tooth_disease_2B1 | 1 |
| LMNA | Cardiomyopathy_dilated_with_conduction_defect_type_1A | 1 |
| LMNA | Cardiomyopathy_dilated | 23 |
| LMNA | Cardiac_disease | 1 |
| LMNA | Cardiac_conduction_system_disease | 1 |
| LMNA | Cardiac_conduction_defects | 1 |
| LMNA | Arrhythmogenic_right_ventricular_cardiomyopathy | 1 |
| MBNL1 | Myotonic_dystrophy | 1 |
| MECP2 | Rett_syndrome_preserved_speech_variant | 1 |
| MECP2 | Rett_syndrome_atypical | 1 |
| MECP2 | Rett_syndrome | 12 |
| MECP2 | Non-fatal_non-progressive_encephalopathy | 1 |
| MECP2 | Neonatal_encephalopathy_severe | 1 |
| MECP2 | Mental_retardation_X-linked | 3 |
| MECP2 | Mental_retardation | 1 |
| MECP2 | Autism_spectrum_disorder | 1 |
| MECP2 | Autism | 1 |
| PAFAH1B1 | Subcortical_band_heterotopia | 1 |
| PAFAH1B1 | Miller-Dieker_lissencephaly_syndrome | 1 |
| PAFAH1B1 | Lissencephaly_isolated | 10 |
| PFN1 | Amyotrophic_lateral_sclerosis_association_with | 1 |
| PFN1 | Amyotrophic_lateral_sclerosis | 6 |
| PPIB | Osteogenesis_imperfecta_recessive | 1 |
| PPIB | Osteogenesis_imperfecta_II | 1 |
| PPIB | Osteogenesis_imperfecta | 2 |
| PPP2R1B | Breast_cancer | 1 |
| RPS19 | Diamond-Blackfan_anaemia | 31 |
| SIN3A | Intellectual_disability_mild | 1 |
| SLC25A12 | AGC1_deficiency | 1 |
| SMC1A | Developmental_delay_epilepsy_delayed_speech_&_encephalopathy | 1 |
| SMC1A | Cornelia_de_Lange_syndrome | 7 |
| SPTAN1 | Intellectual_disability | 1 |
| STAT1 | Mycobacterial_infection | 1 |
| STAT1 | Impaired_mycobacterial_immunity | 1 |

| | | |
|---|---|---|
| TALDO1 | Transaldolase_deficiency | 2 |
| TUBA4A | Amyotrophic_lateral_sclerosis | 3 |
| VIM | Congenital_cataract | 1 |

**Table S4. List of genes, associated disease phenotype and number of HGMD mutations corresponding to the phenotype.**

Table to accompany **Fig. 4C**. The list of gene names mentioned in the legend of Fig.4C is listed in column 1 and the disease phenotype associated with it in column 2. Column 3 highlights the number of HGMD mutations associated with the particular gene and disease phenotype.

**Supplementary Table 5**

| | |
|---|---|
| **mPLsORF0000299804_significant_ECs_0** 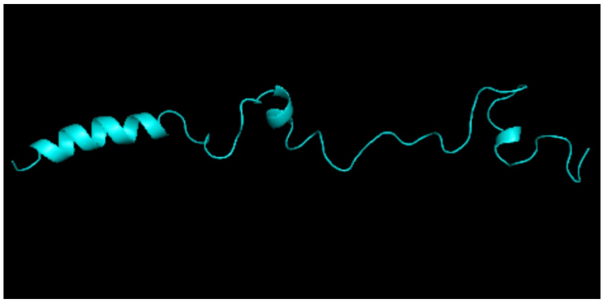 | **mPLsORF0000442197_significant_ECs_0** 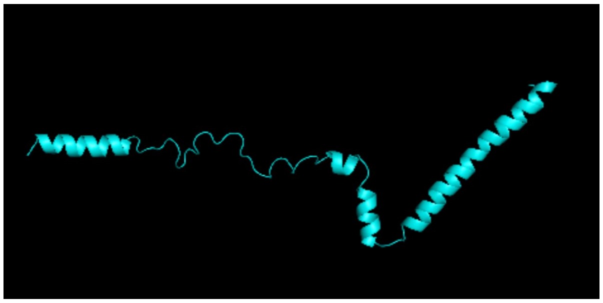 |
| **mPLsORF0000453204_68_8_hMIN** 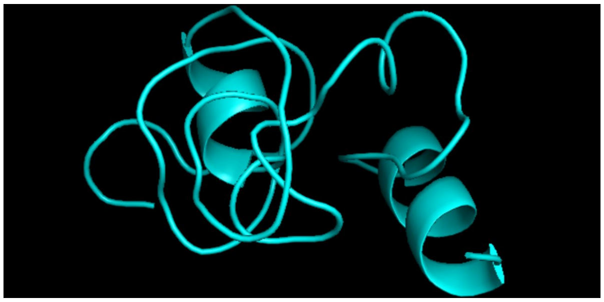 | **mPLsORF0000445046_35_7_hMIN** 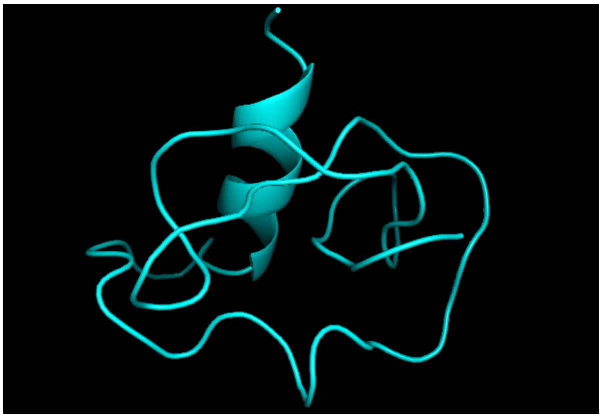 |
| **mPLsORF0000452527_significant_ECs_0** 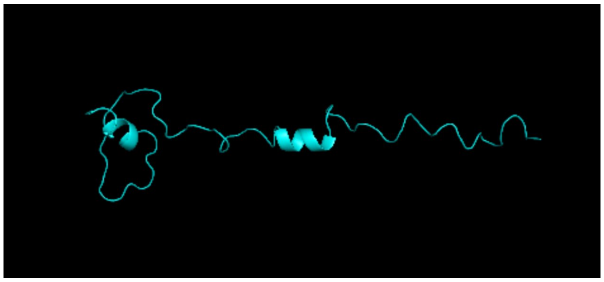 | **mPLsORF0000137329_55_2_hMIN** 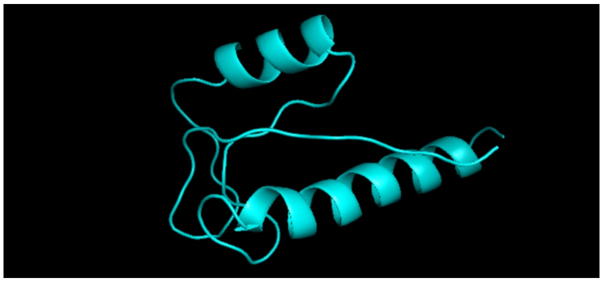 |

| | |
|---|---|
| **mPLsORF0000453225_56_10_hMIN** 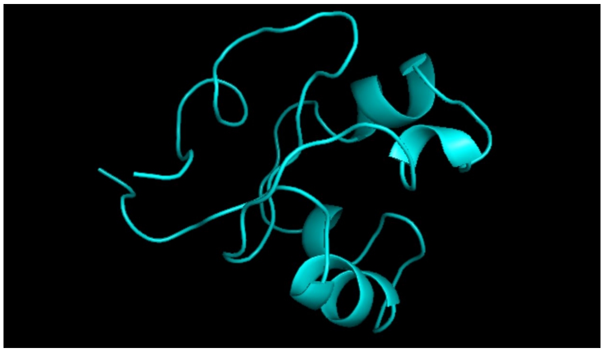 | **mPLsORF0000449632_76_3_hMIN** 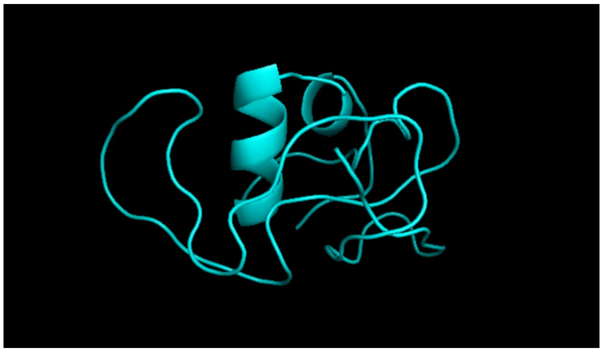 |
| **mPLsORF0000443365_43_8_hMIN** 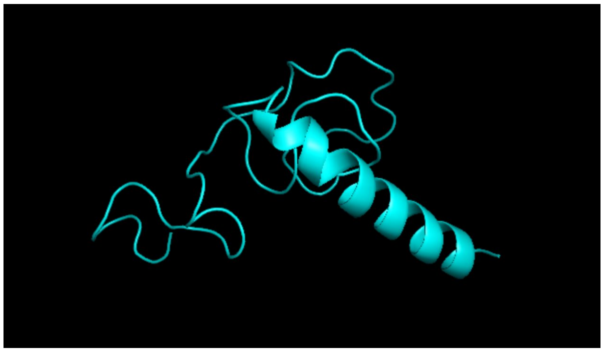 | **mPLsORF0000450681_significant_ECs_0** 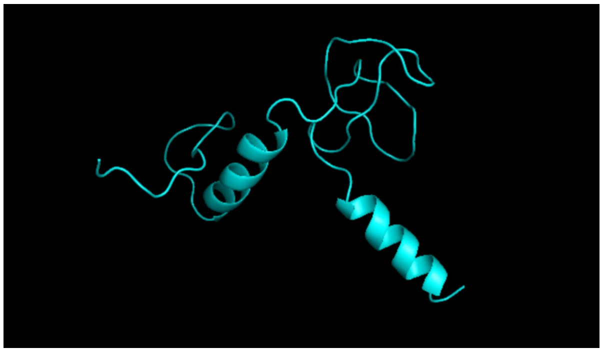 |
| **mPLsORF0000451320_43_8_hMIN** 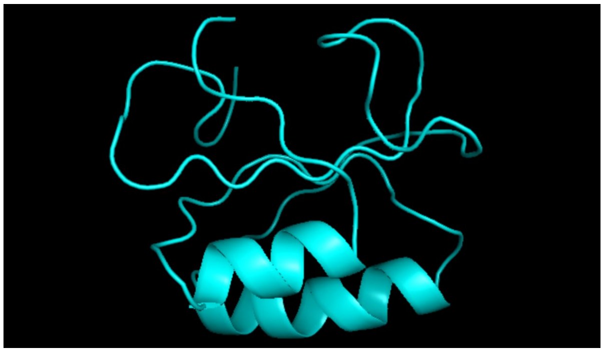 | **mPLsORF0000114575_63_6_hMIN** 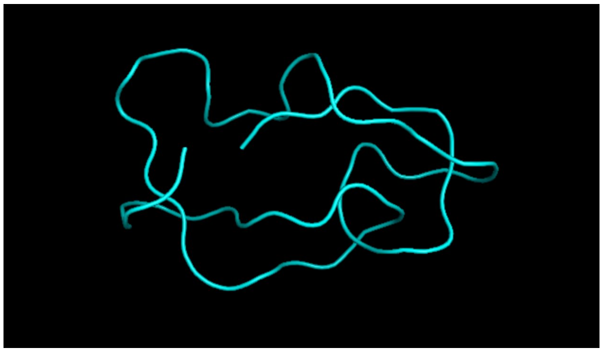 |

| | |
|---|---|
| **mPLsORF0000443648_61_4_hMIN** 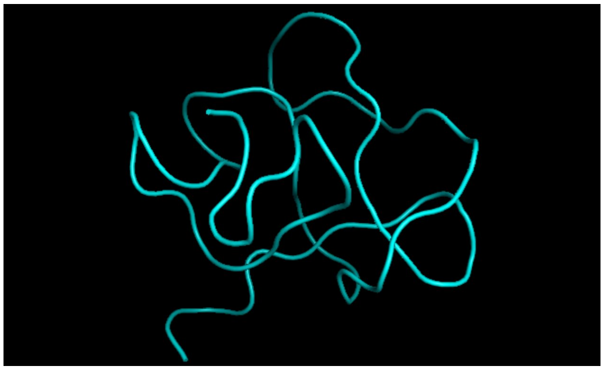 | **mPLsORF0000446045_54_9_hMIN** 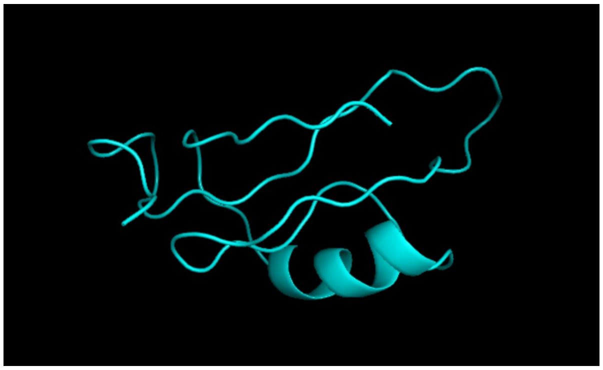 |
| **mPLsORF0000446380_83_10_hMIN** 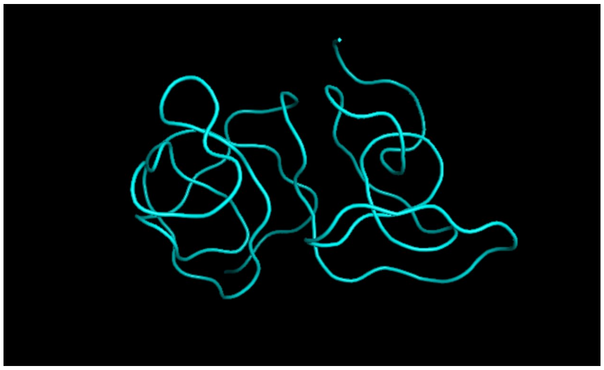 | **mPLsORF0000446071_63_4_hMIN** 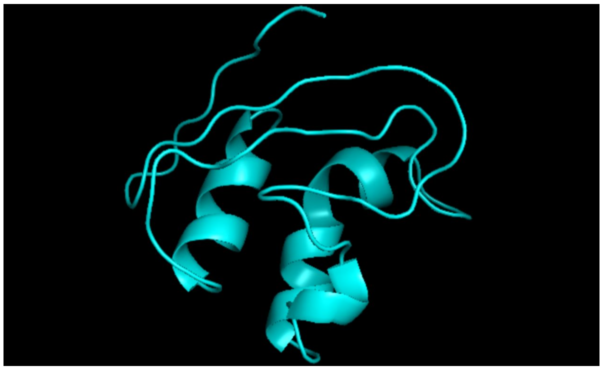 |
| **mPLsORF0000440295_45_4_hMIN** 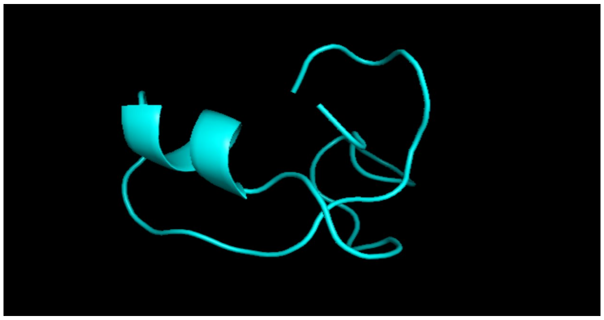 | **mPLsORF0000059717_67_1_hMIN** 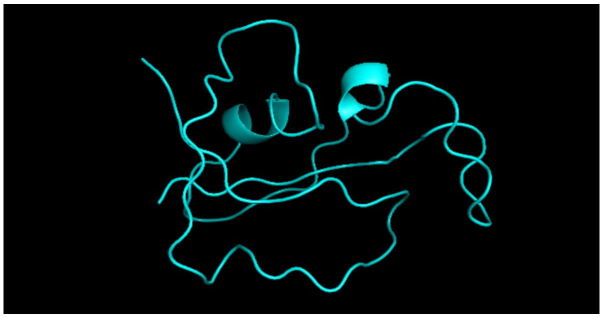 |

| | |
|---|---|
| **mPLsORF0000443953_significant_ECs_0** 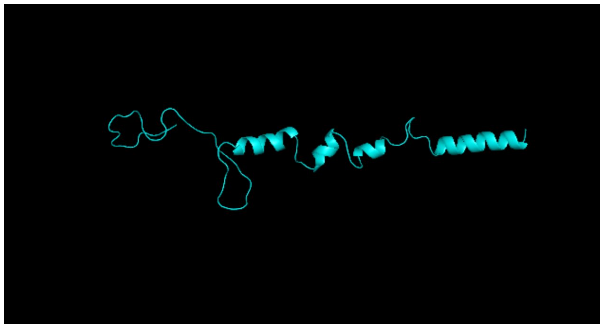 | **mPLsORF0000447578_24_9_hMIN** 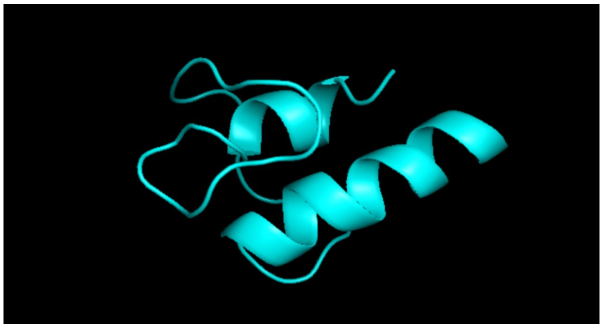 |
| **mPLsORF0000239729_significant_ECs_0** 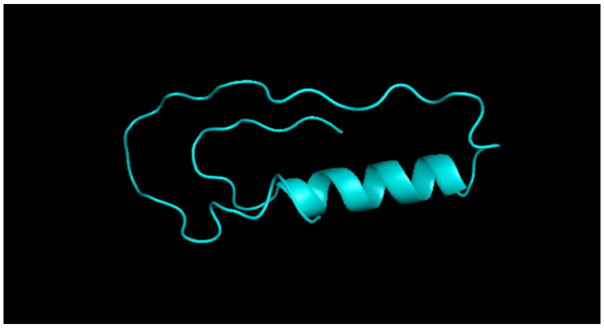 | **mPLsORF0000444809_35_5_hMIN** 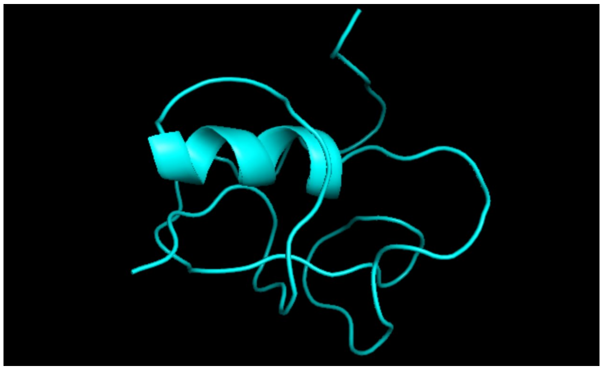 |
| **mPLsORF0000445338_27_2_hMIN** 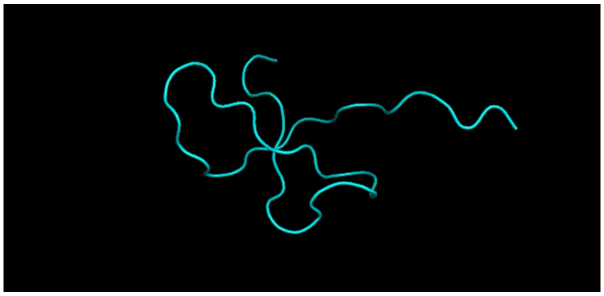 | **mPLsORF0000444619_significant_ECs_0** 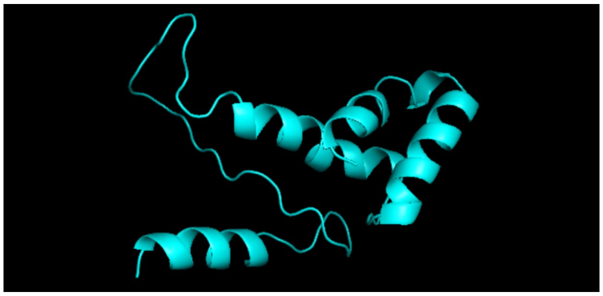 |

**Table S5. Predicted structures of sORFs with both transcriptional and translational evidence.** Structures of 24 sORFs for which we had both transcriptional and translational evidence and predicted with EV fold pipeline are displayed in the table.

**Supplementary Table 6**

| | |
|---|---|
| **IP_220049** 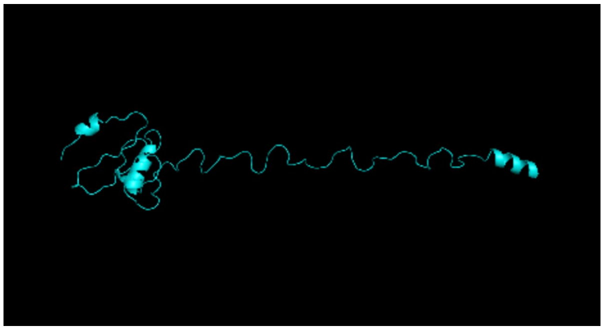 | **IP_189960** 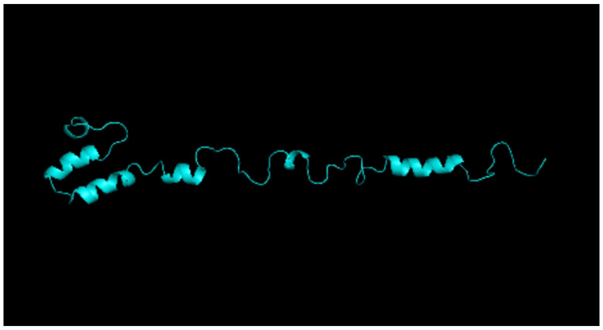 |
| **IP_195050** 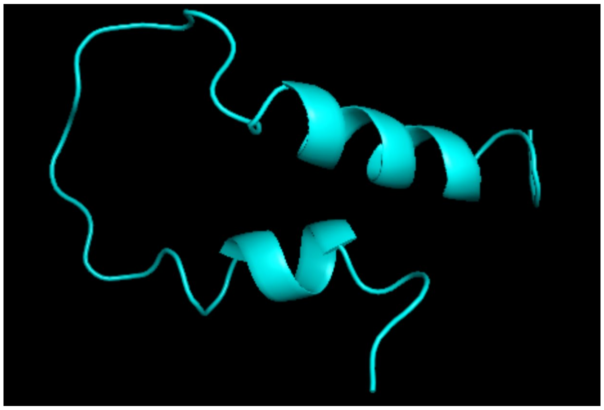 | **IP_195051** 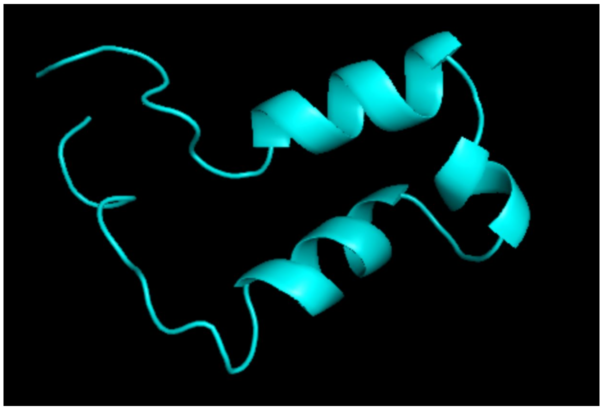 |
| **IP_159154** 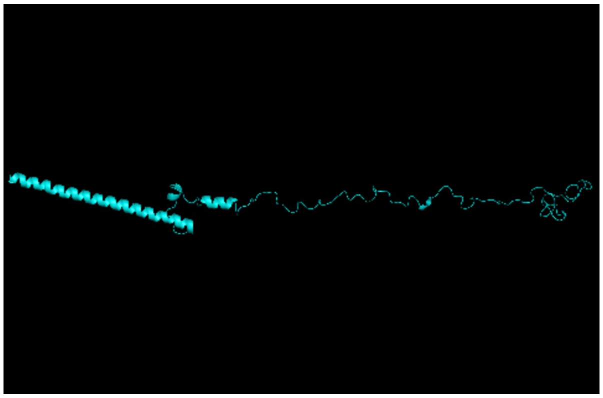 | **IP_275965** 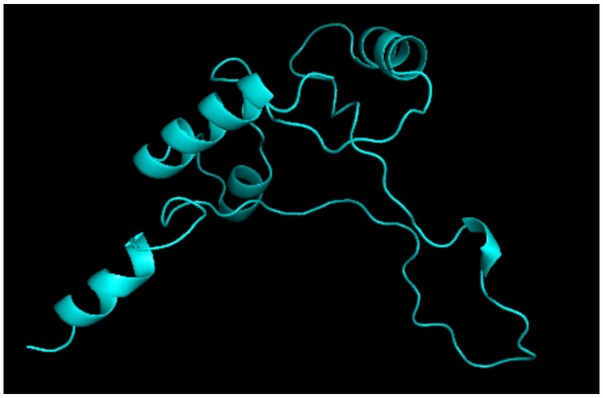 |

| | |
|---|---|
| **IP_233967** 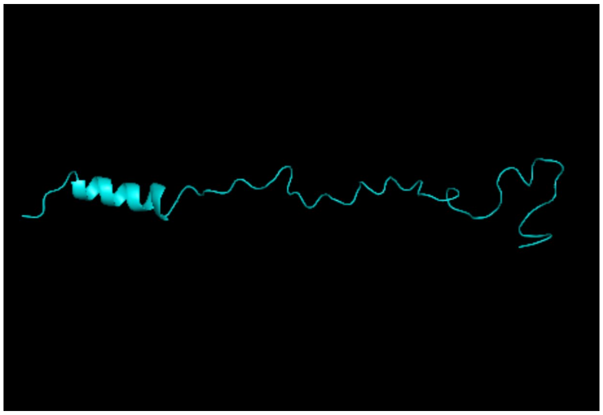 | **IP_233958** 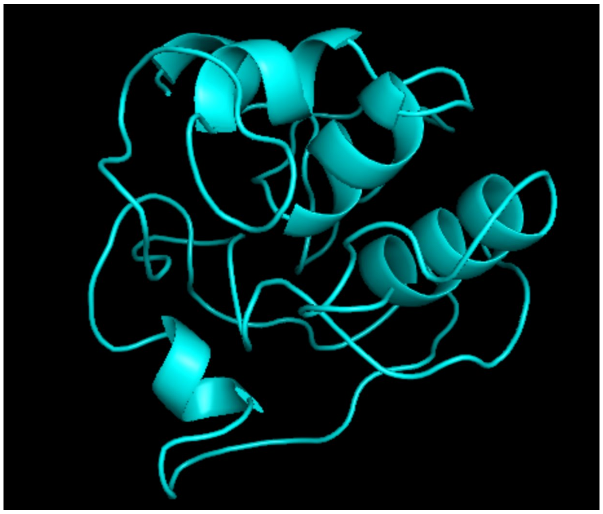 |
| **IP_199907** 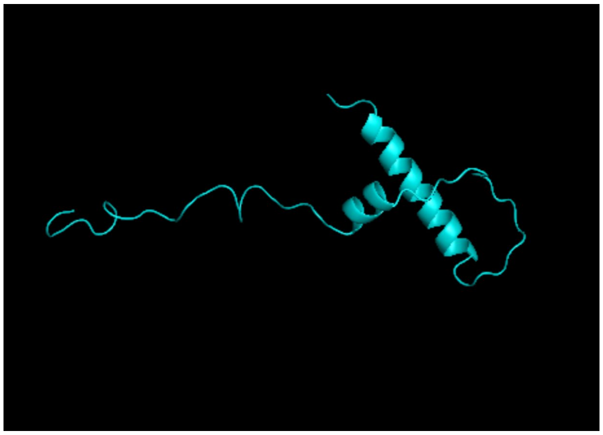 | |

**Table S6. Predicted structures of nine altORFs with both transcriptional and translational evidence.** Structures of nine altORFs for which we had both transcriptional and translational evidence and predicted with EV fold pipeline are displayed in the table.
